# Replisome loading reduces chromatin motion independent of DNA synthesis

**DOI:** 10.1101/2023.03.07.531331

**Authors:** Maruthi K. Pabba, Christian Ritter, Vadim O. Chagin, Janis Meyer, Kerem Celikay, Jeffrey H. Stear, Dinah Loerke, Ksenia Kolobynina, Paulina Prorok, Alice Kristin Schmid, Heinrich Leonhardt, Karl Rohr, M. Cristina Cardoso

**Affiliations:** Department of Biology, Technical University of Darmstadt, Germany; Biomedical Computer Vision Group, BioQuant, IPMB, Heidelberg University, Germany; Institute of Cytology RAS, St. Petersburg, Russia; EMBL Australia Node in Single Molecule Science, UNSW Sydney, Australia; Department of Physics & Astronomy, University of Denver, Denver, CO, USA; Department of Biology II, Ludwig Maximilians University, Munich, Germany

**Author notes:** These authors contributed equally to this work (Maruthi K. Pabba, Christian Ritter, Vadim O. Chagin). Further information and requests for resources and reagents should be directed to and will be fulfilled by the lead contact M. Cristina Cardoso; Karl Rohr.

**Keywords:** Aphidicolin, cell cycle, chromatin tracking, diffusion, DNA labeling, DNA replication, genome architecture, mean square displacement, S-phase

## Abstract

Chromatin has been shown to undergo diffusional motion, which is affected during gene transcription by RNA polymerase activity. However, the relationship between chromatin mobility and other genomic processes remains unclear. Hence, we set out to label the DNA directly in a sequence unbiased manner and followed labeled chromatin dynamics in interphase human cells expressing GFP-tagged PCNA, a cell cycle marker and core component of the DNA replication machinery. We detected decreased chromatin mobility during the S-phase compared to G1 and G2 phases in tumor as well as normal diploid cells using automated particle tracking. To gain insight into the dynamical organization of the genome during DNA replication, we determined labeled chromatin domain sizes and analyzed their motion in replicating cells. By correlating chromatin mobility proximal to the active sites of DNA synthesis, we showed that chromatin motion was locally constrained at the sites of DNA replication. Furthermore, inhibiting DNA synthesis led to increased loading of DNA polymerases. This was accompanied by accumulation of the single-stranded DNA binding protein on the chromatin and activation of DNA helicases further restricting local chromatin motion. We, therefore, propose that it is the loading of replisomes but not their catalytic activity that reduces the dynamics of replicating chromatin segments in the S-phase as well as their accessibility and probability of interactions with other genomic regions.

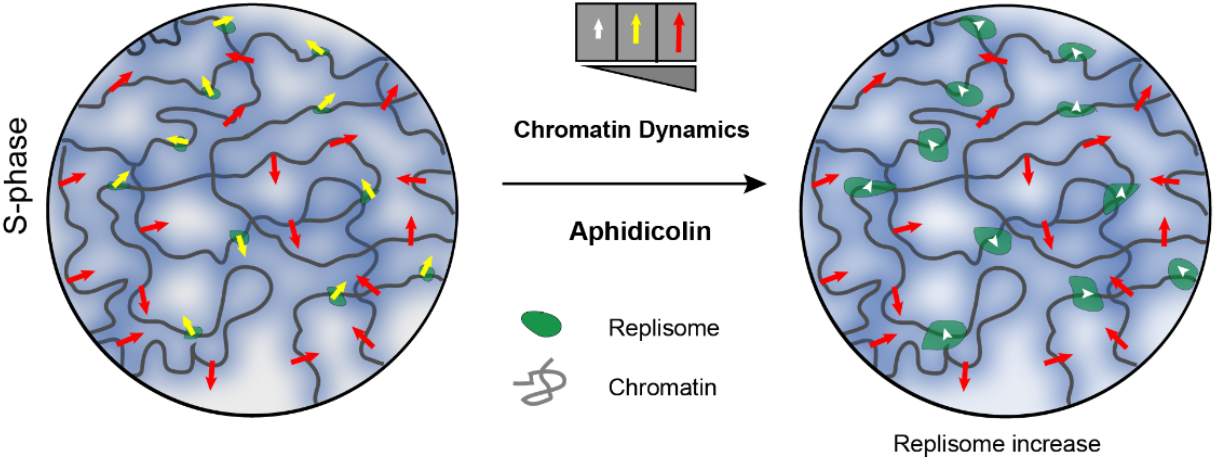

**Highlights:**

- Direct and sequence unbiased labeling of DNA genome-wide
- DNA labeled chromatin is more mobile in G1/G2 relative to the S-phase
- Restriction of chromatin motion occurs proximal to sites of DNA replication
- Loading of replisomes, even in the absence of processive DNA synthesis, restricts chromatin motion

## INTRODUCTION

Dynamic yet functionally stable organization of cellular processes is a crucial feature of biological systems, which allows them to respond to external stimuli and survive. The eukaryotic nucleus is a complex subcellular organelle where DNA metabolism including its replication, repair and transcription, occurs. Eukaryotic DNA is organized in the nuclear space by interactions with histones and architectural proteins to form a hierarchy of domains and compartments of the interphase chromatin. Nuclear architecture is dynamically modulated due to the binding of biomolecules and epigenetic changes of the chromatin. It is also interdependent with DNA metabolism mediated by the action of enzymes on the chromatin. The maintenance of the DNA (including its replication and repair) and its transcription into RNA are spatio-temporally organized within the cell nucleus.

Analysis of the local chromatin dynamics in live cells revealed that an essential aspect of interphase chromatin is its mobile nature (Gasser, 2002; Marshall et al., 1997). The movement of chromatin loci was shown to be consistent with an anomalous (constrained) diffusion model (Scipioni et al., 2018; Shukron et al., 2019). This model indicates that a single chromatin locus is corralled within a sub-micron radius and exhibits random diffusion motion and will execute multiple random jumps into neighboring compartments (Bronshtein et al., 2016; Chubb et al., 2002; Heun et al., 2001; Levi et al., 2005; Marshall et al., 1997). This behavior, which we refer to as local chromatin diffusion (LCD), has been described in multiple systems, suggesting that it is likely to represent a fundamental aspect of chromatin dynamics in eukaryotes.

According to the current paradigm, the 4D organization of the chromatin inherently includes its physical properties as a long polymer (Esposito et al., 2021, 2019), while stochastic thermodynamically driven events are likely to play a key role in the domain organization of the chromatin (Conte et al., 2020; Shin and Brangwynne, 2017) and in the regulation of genomic processes (Hnisz et al., 2017; Kilic et al., 2019; Laghmach et al., 2021; Nozaki et al., 2017; Spegg and Altmeyer, 2021; Uchino et al., 2022).

Some studies have reported that chromatin mobility is enhanced due to active transcription (Gu et al., 2018; Tunnacliffe and Chubb, 2020), whereas others report rather a decrease in mobility (Mach et al., 2022). Furthermore, other studies report diverse effects of RNA polymerase II inhibition on chromatin motion (Germier et al., 2017; Ku et al., 2022; Shaban et al., 2018). It has been also shown that the removal of RNA polymerase II from chromatin relaxes chromatin and increases its mobility (Babokhov et al., 2020). Conversely, there is an established view that chromatin mobility at the sites of double-strand DNA breaks increases concomitant with their repair (Eaton and Zidovska, 2020; Hauer and Gasser, 2017; Hauer et al., 2017; Nagai et al., 2010). Analysis of fluorescently tagged histones using displacement correlation spectroscopy has shown that chromatin undergoes coherent micron-scale motion at the time scales of 5-10 seconds independently of the cell cycle stage in mammalian cells (Zidovska et al., 2013). This coherent motion extended beyond individual chromosomes, suggesting mechanical coupling between chromosomes. Furthermore, the correlated motion of chromatin was ATP-dependent and completely disappeared upon DNA damage induction (Eaton and Zidovska, 2020; Zidovska et al., 2013).

DNA replication is a highly conserved energy-dependent process occurring in S-phase of the cell cycle, when chromatin structures undergo extensive reorganization to facilitate DNA synthesis (Vincent et al., 2008). An early study in budding yeast (Heun et al., 2001) demonstrated that individual heterologous loci became constrained in S-phase when integrated close to early- and late-firing replication origins, but not at the telomeric or centromeric regions. However, changes in chromatin mobility in S-phase were not observed when analyzing it at the level of chromosome territories in mammalian cells (Walter et al., 2003). Recent work using a CRISPR-based DNA imaging system suggests that local chromatin motion is restricted upon S-phase entry and more markedly in mid-late S-phase (Ma et al., 2019).

Altogether, it is not clear whether and how chromatin mobility changes during DNA replication and a mechanism behind the changes in chromatin motion. Therefore, it is important to address how changes in structure and metabolism of chromatin affect its mobility. It is quite intriguing to postulate that the process of genome duplication in mammals, which is performed at the level of naked DNA and involves local chromatin decondensation and rearrangements at the complete hierarchy of domains (Baddeley et al., 2010; Chagin et al., 2019; Löb et al., 2016; Sadoni et al., 2004; Sporbert et al., 2002) is associated with changes in chromatin mobility. Furthermore, it is tempting to speculate that the modulation of LCD may play a regulatory role; for example, by helping to define the transcriptional profile of the nucleus, by provoking collisions between regulatory regions, promoter regions and transcription factories. These events could be halted or slowed down during the replication of the genome, avoiding collisions of the transcription with the replication machineries. An alternative but not mutually exclusive model is that changes in LCD result from the execution of nuclear processes such as transcription or replication. This is particularly appealing as DNA/RNA helicases and polymerases are, in essence, motor proteins that reel DNA through. To distinguish between these possibilities, alterations in LCD have to be characterized within the context of relevant nuclear processes and by labeling DNA directly and in an unbiased manner.

The process of genome replication has a particular and intrinsic connection between chromatin organization and the spatio-temporal progression of genome replication (reviewed in (Mamberti and Cardoso, 2020)). In that sense, firing of origins of replication by the activation of DNA helicase complexes followed by the loading of synthetic polymerase complexes tracks chromatin compaction and upon DNA duplication the focal chromatin organization at multiple hierarchical levels is preserved and can be detected over several cell generations (Cremer et al., 2020; Jackson and Pombo, 1998; Sadoni et al., 2004; Sparvoli et al., 1994). Importantly, genome replication is the only DNA metabolic process that encompasses the entire genome, thus ensuring the preservation of the genetic material upon cell division.

As most of the studies introduce artificial DNA sequences in genomic loci and use a large array of chromatin binding proteins to visualize the loci, chromatin dynamics may be altered in the subsequent process (Germier et al., 2017). Therefore, a more direct way to measure chromatin dynamics is to label and track the DNA directly (Schermelleh et al., 2001). A similar procedure has previously been used to mark chromosome territories and characterize their long-term rearrangements (Bornfleth et al., 1999; Pliss et al., 2009; Walter et al., 2003).

In this study, we investigated the mobility of chromatin in human cells, focusing on how changes in chromatin mobility are influenced by cell cycle progression and, in particular, DNA replication. To achieve this, we performed a detailed analysis of chromatin mobility in S-phase by combining locus-independent global labeling of DNA with reliable particle tracking. Measurement of the DNA content of the labeled structures allowed us to elucidate whether DNA replication affects chromatin mobility at the level of replication domains. Our results show that chromatin mobility generally decreases during S-phase and, in particular, at the proximity of the DNA polymerase complexes. Furthermore, we extended our study to dissect mechanisms behind the S-phase related changes in chromatin mobility and inhibited DNA synthesis using small molecule inhibitors. We showed that chromatin mobility is further decreased in S phase after inhibition of DNA synthesis. These results imply that loading of the polymerase complexes rather than the synthesis of DNA *per se* restraints DNA mobility.

## RESULTS AND DISCUSSION

### Genome-wide labeling of DNA and quantification of labeled chromatin domains

To evaluate LCD (local chromatin diffusion) relative to the cell cycle stage, we first developed an experimental system to monitor both replication and chromatin changes in living cells in real-time. We generated HeLa cell lines that stably express GFP-tagged proliferating cell nuclear antigen (PCNA) and single-stranded DNA binding protein (RPA) (Methods, Supplementary Table 1). We transfected fluorescent PCNA plasmid to label replication sites in human normal diploid fibroblasts (IMR90) (Nichols et al. 1977). PCNA is a core component of the DNA replication machinery and a marker for cell cycle progression (Figure 1A) (Chagin et al., 2016; Easwaran et al., 2005; Leonhardt et al., 2000; Moldovan et al., 2007; Prelich et al., 1987). To visualize the mobility of native chromatin, we took the advantage of the ongoing DNA replication. We delivered a pulse of the fluorescently labeled nucleotide Cy3-dUTP by electroporation into an asynchronously growing population of human HeLa GFP-PCNA tumor cells and human diploid IMR90 fibroblasts, which allowed us to study chromatin dynamics in a global genome-wide manner (Methods, Figure 1A). The nucleotide is incorporated into the nascent DNA of the cells in various periods of S-phase, effectively labeling the chromatin directly in an unbiased manner (Manders et al., 1999; Sadoni et al., 2004; Schermelleh et al., 2001).

**Figure 1:**
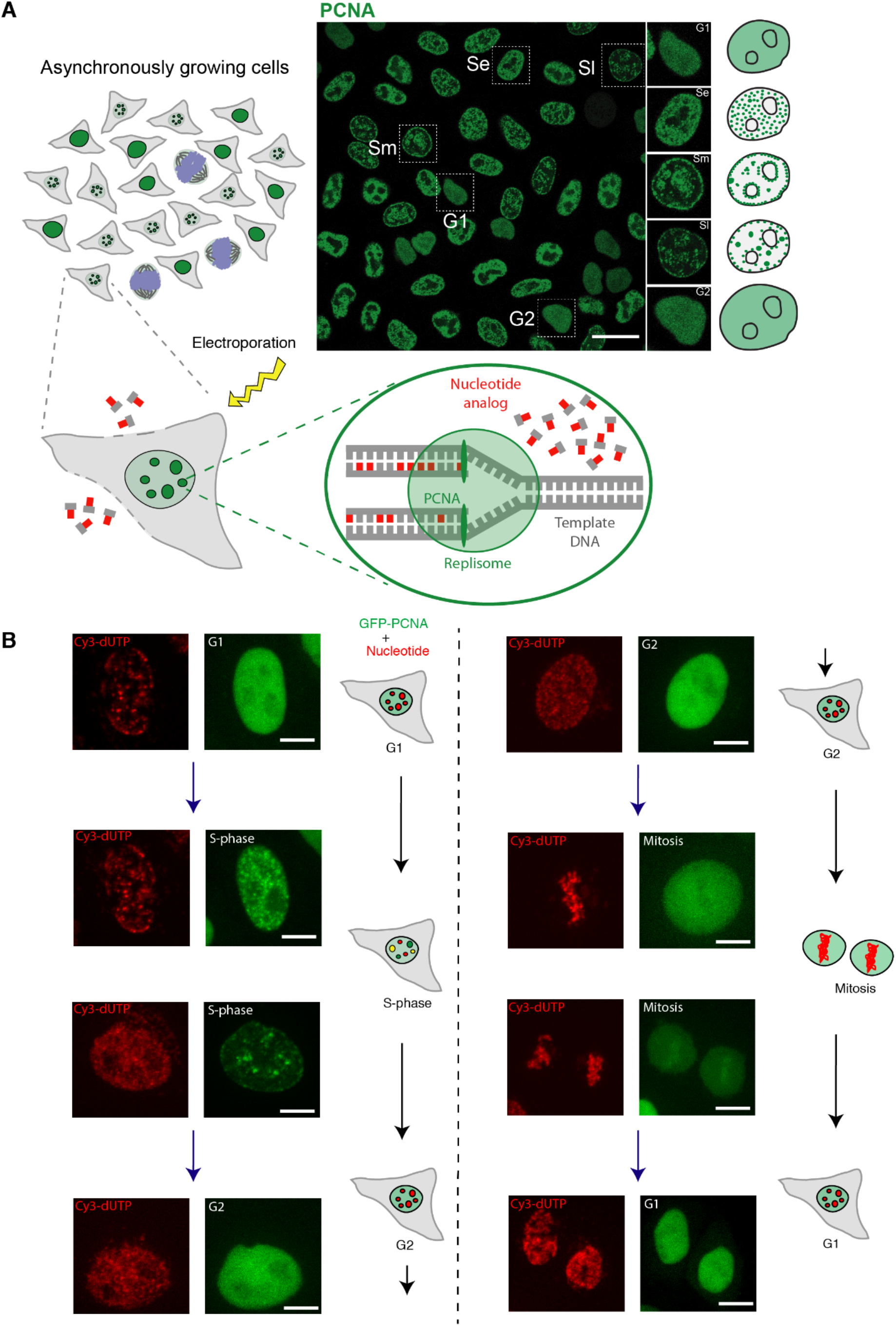
Incorporation of Cy3-dUTP in HeLa cell nuclei labels the whole genome randomly and with equal probability. **(A)** Schematic illustration of the labeling system for monitoring chromatin mobility and cell cycle progression. During S-phase, PCNA accumulates within the nucleus at sites of active DNA replication and exhibits a distinct puncta pattern. During G1 and G2, GFP-PCNA is diffusely distributed throughout the nucleus. Asynchronously growing populations of cells were exposed to electroporation to promote the uptake of Cy3-dUTP. In cells undergoing DNA replication, this fluorescent nucleotide is incorporated into nascent DNA strands at sites of active DNA replication, resulting in the direct fluorescent labeling of genomic segments. Based on the PCNA pattern, different cell cycle stages can be differentiated as shown in the image on the right (Se-Early S, Sm-Mid S, SL-Late S, G1/G2-Gap phases, Green - PCNA). **(B)** After Cy3-dUTP labeling (shown in red), cells were followed by time lapse microscopy to identify the cell cycle (sub)stages and their progression. The representative images of different cells using time lapse microscopy were shown to depict the patterns of PCNA(shown in green) in each substage and their change over time (colocalized signals in yellow). This was used to classify cells in G1, S and G2 phases of the cell cycle for motion analysis. Approximately 18 to 24 hours after nucleotide electroporation, Cy3-dUTP-labeled cells were imaged for motion analysis (see also movies 1-5).The contrast of the images was adjusted linearly for visualization purposes. Scale bar: 5 µm.

The Cy3-dUTP labeled chromatin structures were stable over the cell cycle progression and in subsequent cell cycles. Using time lapse microscopy, we followed the cells that incorporated nucleotides in the initial S phase stage over subsequent cell cycles. We used GFP-PCNA nuclear pattern to determine the cell cycle stages and sub-periods of S-phase (Methods, Microscopy). This allowed us to classify cells in different cell cycle stages and sub-periods of S-phase (G1, early S, mid S, late S, G2), which is illustrated in Figure 1B (see also movies 1-5). With this approach, DNA labeled during the pulse of Cy3-dUTP nucleotide corresponds to genomic regions replicated concomitantly during an S-phase sub-period. Since LCD measurements depend on the object size, it was important to evaluate the size of the labeled DNA domains. This allowed us to correlate the chromatin domain sizes and their diffusion rates. For this purpose, we measured the total DNA amount in a cell and the fraction of it that corresponded to the labeled domain (Methods, DNA quantification). First, we applied chemical fixation to cells labeled with Cy3-dUTP using formaldehyde. The total DNA was then labeled using the DNA dye DAPI. Next, we segmented the entire nucleus as well as the individual labeled chromatin foci within the same cell. The fraction of DAPI intensity within the segmented replication focus (I_RFi_) over the total DNA intensity within the cell (I_DNA total_) yields the amount of DNA present per labeled chromatin focus (Figure 2A, Supplementary Figure S1). Since nuclear DNA amount doubles continuously throughout the S-phase (Chagin et al., 2016; Leonhardt et al., 2000), (Figure 2B) it was important to scale the total DNA amount by a correction factor depending on S-phase sub-stage to measure the DNA amount per focus more accurately. The relative amount of DNA throughout the cell cycle stages and sub-stages of S-phase was calculated and plotted as histograms, with the mean of the histogram for each cell cycle (sub)stage constituting the cell cycle correction factor (Figure 2B). The fraction of DAPI intensities were corrected by multiplication with the genome size corresponding to the cell cycle stage. The genome size of HeLa Kyoto cells is GS = 9.682±0.002 Gbp (Chagin et al., 2016) and for IMR90 fibroblasts the genome size is 6.37 Gbp as measured earlier (Nichols et al. 1977). We plotted the DNA amount present in each replication focus in Figure 2C for HeLa on left and IMR90 on right. The highest frequency of average DNA amount per focus (mode + 1 bin) was about 300-600 kbp of DNA (Figure 2C). Altogether, with our labeling approach, we labeled DNA domains of sizes ranging from 0.5 Mbp to 10 Mbp, with the vast majority corresponding to 0.5 Mbp, which correspond well to multi-loop chromatin domains corresponding in size to topological associated domains (TADs) (reviewed in (Giorgetti and Heard, 2016).

**Figure 2:**
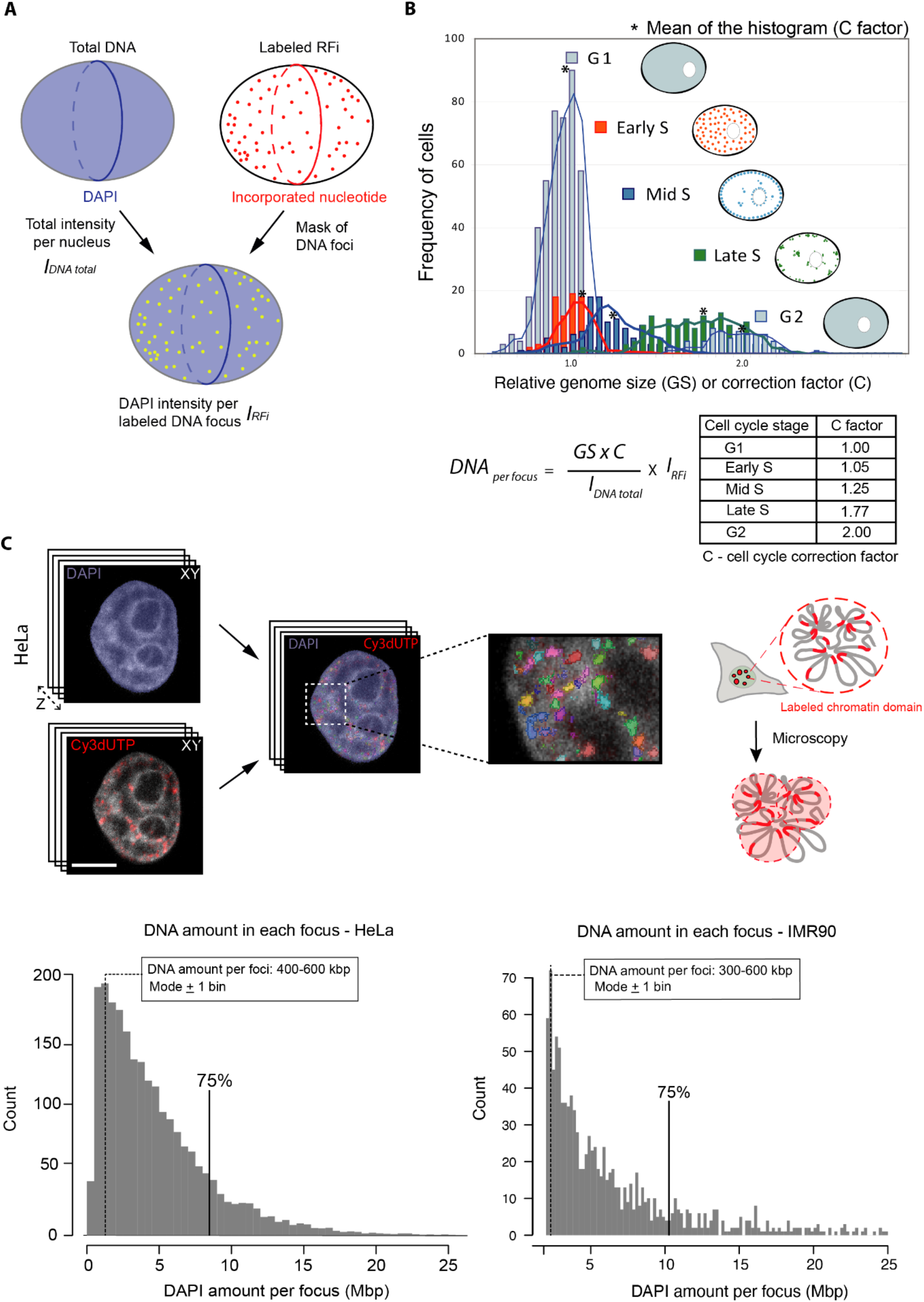
Principle of measuring DNA content per labeled DNA focus using confocal data. **(A)** To determine the amount of DNA per labeled DNA focus; we used the total DAPI signal (DNA amount) of the segmented whole nucleus (I_DNA TOTAL_). The DNA intensity per labeled DNA focus within the segmented foci is obtained (I_RFi_) by masking the replication foci and estimating the corresponding portion of DAPI signal. **(B)** Throughout the S-phase progression the amount of DNA increases twofold from early to late S-phase. The amount of DNA present in the nucleus at a particular cell cycle stage can be determined by measuring the DNA amount in the population of cells, while using the PCNA pattern to determine the cell cycle stage and S-phase sub-stage (see also Figure 1). The relative mean amount of DNA of each of the cell cycle (sub-)stages is used to calculate the cell cycle correction factor. The cell cycle correction factor (C/cell cycle stage) was estimated as: 1.0/G1; 1.05/early S-phase; 1.25/mid-S-phase; 1.77/late S-phase, 2/G2. The G1 genome size (GS) for HeLa cells is 9.7 Gbp (Chagin et al., 2016). The amount of DNA per labeled focus is the ratio of I**_RFi_** and I**_DNA TOTAL_** multiplied by C x GS. **(C)** The illustration on right depicts the imaging of labeled replication foci using confocal microscopy. DNA quantification of replication labeled foci in tumor HeLa and normal diploid IMR90 cells was done by imaging full z-stacks volume of chromatin labeled with Cy3-dUTP and DNA with DAPI and imaged using confocal spinning disk microscopy (Supplementary Figure S1, Supplementary Table 5). The histogram represents the DNA amount per focus for labeled S-phase cells (N= 30 cells) for HeLa and IMR90 cells. The mode ± 1 bin of the histogram represents the highest frequency of average size of replication domains labeled (300-600 kbp). Scale bar: 5 µm.

### Chromatin motion decreases in the S-phase of the cell cycle relative to the G1 and G2 phases

To determine how the global dynamics of chromatin changes during cell cycle progression, we used LCD measurements relative to the cell cycle stage. Live cell time-lapse image sequences of HeLa and IMR90 cells after labeling chromatin with Cy3-dUTP were obtained and motion analysis was performed to determine the type of motion (Figure 3A, Methods). Normal diffusion or Brownian motion is a linear diffusion model with ɑ = 1 and when ɑ > 1 it is termed super diffusion. First, the cells were annotated according to the different cell cycle stages (G1, S, G2) based on the PCNA subnuclear pattern (Methods). PCNA forms puncta or foci at the active replication sites during S-phase and this was used to classify cells in S-phase. We were able to distinguish between G1 and G2 cells, even though they exhibit a similar diffused PCNA subnuclear distribution, based on the information on the preceding cell cycle stage from the time lapse analysis performed after Cy3-dUTP labeling (Methods, Microscopy). Specifically, cells with diffusely distributed PCNA signal which had previously undergone mitosis were in G1 phase, whereas the ones with similar diffuse PCNA pattern that had previously undergone S-phase (punctated PCNA pattern) were classified as being in G2 phase (Figure 1B). The PCNA signal was also used to segment the nucleus, and the individual chromatin foci were detected within the segmented nuclei. Probabilistic tracking was performed to obtain individual chromatin trajectories (Figure 3B; Supplementary Figure S2). In case of IMR90 cells, affine image registration was performed using the method in (Celikay et al., 2022) to address the stronger cell movement compared to HeLa cells. This was followed by a Mean Square Displacement (MSD) analysis to determine the chromatin motion in different cells (Supplementary Figure S2). In fixed cells, labeled chromatin foci showed almost no motion, which was used as a control for the stability of the imaging system and the tracking protocol. We plotted the MSD over time (up to 20 s) for chromatin foci in cells from different cell cycle stages as well as for fixed cells (Figure 3C). As we focussed on chromatin mobility changes during S-phase, the G1, G2 cells were together in Figure 3C. The mean square displacement curves of G1, G2, S-phase (separated) are plotted in Supplementary Figure 2B.

**Figure 3:**
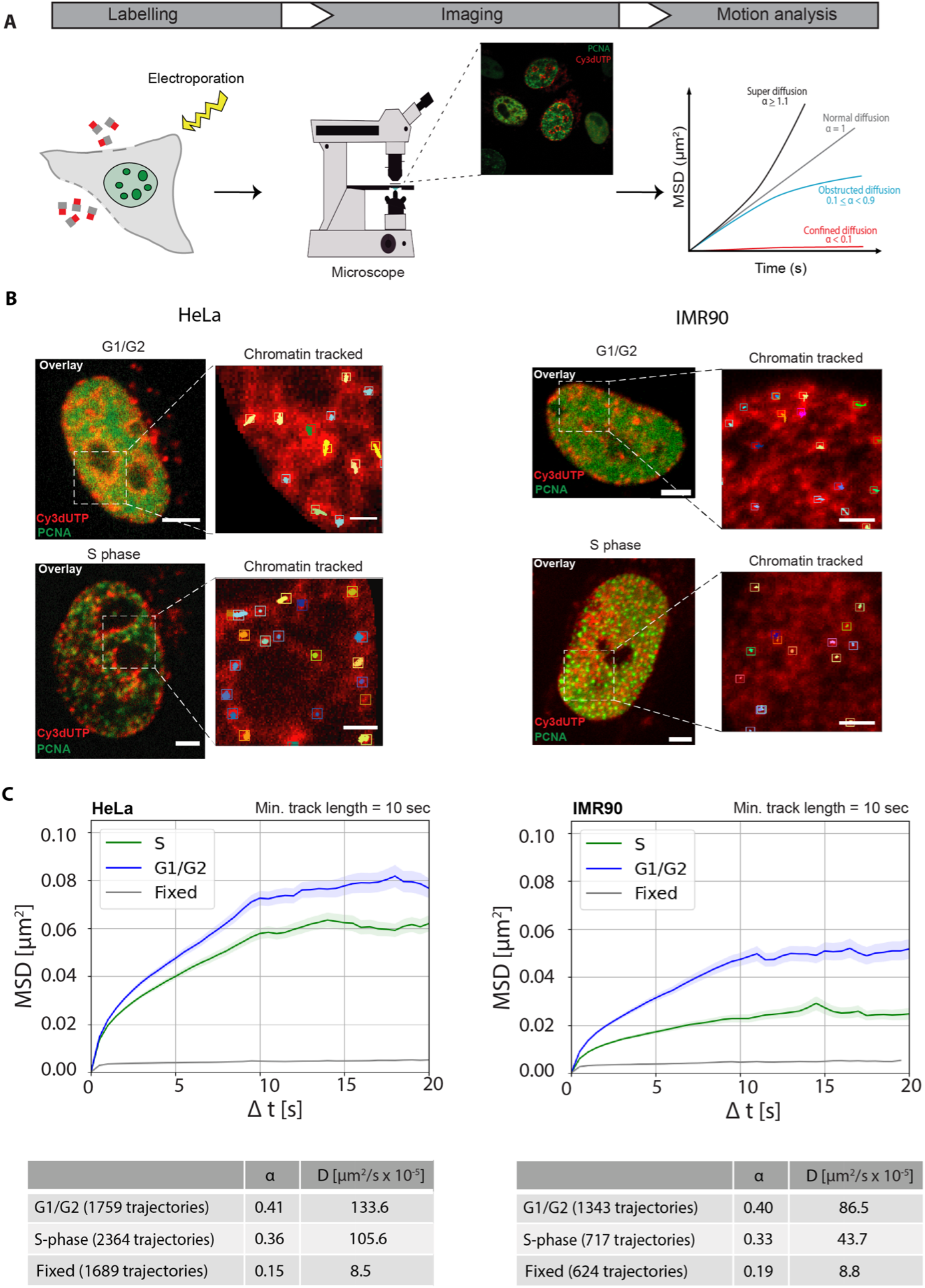
Single-particle motion analysis of labeled chromatin throughout the cell cycle. **(A)** Brief schematics of the main steps of motion analysis starting with chromatin labeling using fluorescently labeled nucleotides via electroporation, followed the next day (i.e., cell cycle) by confocal time-lapse imaging of the chromatin channel and performing motion analysis on computed tracks to determine the diffusion rates of chromatin (Methods). **(B)** Overlay images of HeLa Kyoto and IMR90 cells expressing GFP/miRFP-PCNA and labeled chromatin (Cy3dUTP) in different cell cycle stages (G1/G2 – diffused PCNA, S-phase – PCNA puncta). Cropped region (white box) showing the chromatin tracks of individual foci in both G1/G2 and S-phase cells. The aggregates of Cy3-dUTP that are found in cytoplasm are excluded from the analysis using a nuclear mask. See also movies 1-5. **(C)** Result of motion analysis of computed chromatin tracks for different cell cycle stages (Supplementary Figure S2) for HeLa and IMR90 cells. Mean Square Displacement (MSD, μm^2^) curves were plotted over time (s). Mean square displacement curves for G1/G2, S-phase, fixed cells with a minimum track length of 10 seconds and a total time of 20 seconds were plotted with error bars (SEM - standard error of the mean) representing the deviations between the MSD curves for an image sequence in transparent color around the curve. Scale bar: 5 µm. Insets scale bar: 1 µm.

We observed significantly constrained global chromatin motion in S-phase cells compared to non-replicating G1/G2 cells suggesting that chromatin was more constrained during DNA replication. This effect was stronger in IMR90 cells compared to HeLa Kyoto. The table shows the average diffusion rates (Figure 3D). For HeLa average diffusion rate of chromatin in G1/G2 was D = 133.6 μm^2^/s x 10^-5^, whereas the diffusion rates dropped to D = 105.6 μm^2^/s x 10^-5^ during S-phase (Figure 3C). For IMR90 average diffusion rate of chromatin in G1/G2 was D = 86.5 μm^2^/s x 10^-5^, whereas the diffusion rates dropped to D = 43.7 μm^2^/s x 10^-5^ during S-phase (Figure 3C). We computed the ɑ values in different stages, which define the type of diffusion motion. Chromatin exhibited anomalous subdiffusion or obstructed diffusion with 0.1 < ɑ < 0.9. Anomalous diffusion of cellular structures including chromatin with α values between 0.1 and 0.9 have been reported (Bronshtein et al., 2015; Ghosh and Webb, 1994; Mach et al., 2022; Oliveira et al., 2021; Simson et al., 1998; Smith et al., 1999).

In agreement with our results, it has been initially reported in yeast that some chromatin loci are constrained during S-phase (Heun et al., 2001). This study has been extended to the mammalian genome using the CRISPR targeted labeling of specific genomic loci to demonstrate that the S-phase mobility of the labeled chromosomal loci decreases in S-phase compared to G1/G2 (Ma et al., 2019). Another study reported that during DNA replication there were changes in chromatin mobility due to an unknown mechanism (Nozaki et al., 2017).

As we measured decrease in global chromatin motion during S-phase, which includes labeled chromatin which is replicating as well as non-replicating, we next focused the study on the microenvironment of active replication sites. This opened the question of whether the loading of the replisome on chromatin or its enzymatic activity during S-phase actively restricted chromatin motion. Hence, we analyzed in detail the spatial relationship of chromatin diffusion and DNA replication sites.

### Chromatin motion decreases in proximity to active DNA replication sites

DNA replication involves systematic and structured assembly of proteins directly or indirectly involved in DNA synthesis. DNA replication factors such as DNA polymerase clamp protein (PCNA), the DNA helicase complex that unwinds DNA, and the single-stranded DNA binding protein A (RPA) complex, which stabilizes and protects the single stranded DNA exposed upon helicase activity are illustrated in (Figure 4A). The DNA polymerase clamp PCNA, one of the most well studied replication proteins, was used to mark the active DNA replication sites. To test whether DNA replication factors restrict chromatin motion, we performed proximity analysis (Methods, Figure 4B). As before, we used Cy3-dUTP to label chromatin in the S-phase of the previous cell cycle. We then followed the cells through the cell cycle to select cells in which some of the sites of labeled chromatin were replicating in the S-phase of the next cell cycle at the time of observation. This allowed us to image the labeled chromatin marked in the previous cell cycle together with a live-cell marker (fluorescent PCNA) for the active replication sites in the next cell cycle (Figure 4B). Subsequently, we measured the mobility of chromatin from these S-phase cells at increasing center to center distances (CCD, R) from active replication sites (Figure 4C). For chromatin outside the CCD with replication sites in these S-phase cells, we observed the same diffusion rate as before for the chromatin foci in S-phase cells with no differentiation of whether chromatin was actively replicating or not (Figure 3). However, we observed that the chromatin in the proximity of replication sites (actively replicating) had more restricted motion when located up to 1 μm (center to center) distance to an active replisome, and this effect vanished at higher distances (Figure 4C, Supplementary Figure S3). These data indicate that the reduction of chromatin motion in S-phase is spatially correlated with DNA replication and suggest that DNA synthesis restricts chromatin motion. Hence, we next investigated whether loading of the DNA replication machinery restricts chromatin motion or alternatively DNA synthesis activity is responsible for it.

**Figure 4:**
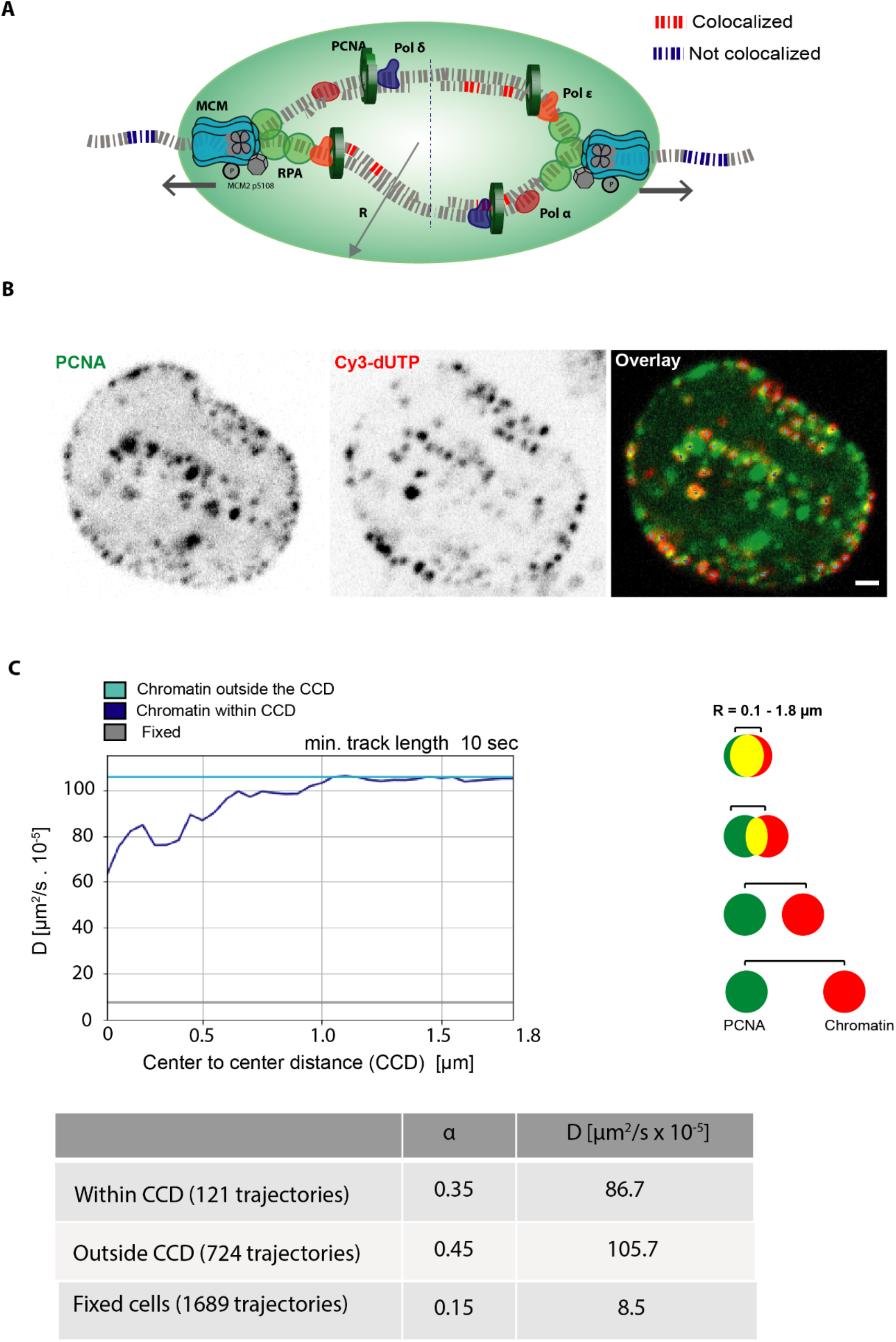
Analysis of chromatin mobility versus distance (proximity) to the DNA replication machinery. **(A)** Schematic illustration of replisome components (helicase, replication protein A, proliferating cell nuclear antigen) actively replicating chromatin. The geometric centers of the labeled chromatin foci and labeled replication sites were first defined. The chromatin within the defined center to center distance (CCD) to a PCNA labeled replication site is defined as chromatin that is within CCD and, otherwise, is defined as outside the CCD. **(B)** In order to obtain mobility information of labeled chromatin in the proximity of PCNA foci (active replication sites) one frame of PCNA channel was acquired followed by 50 frames of the chromatin channel with a frame rate of 0.5 seconds. The images show the spatial distribution of PCNA and chromatin foci (Cy3-dUTP). **(C)** The graph represents the average diffusion rates of the mean square displacement curves (MSD) of chromatin within the center to center distance (CCD) and chromatin outside the center to center distance (CCD) with increasing distance (R) measured between the centers of PCNA and chromatin foci (Supplementary Figure S3).The table below provides the detailed information on number of trajectories per individual sample along with average diffusion rates (µm^2^/s x 10^-5^) and anomalous α coefficient showing subdiffusion at 0.5 µm center to center distance (CCD). Scale bar: 1 µm.

### DNA synthesis inhibition leads to activation of DNA helicases and accumulation of single stranded DNA binding proteins and DNA polymerases

During DNA replication, replisome components are assembled at the origin of replication to form an active replisome (Casas-Delucchi and Cardoso, 2011; Yao and O’Donnell, 2010, 2016). To test whether the process of DNA synthesis itself is responsible for constraining chromatin, we analyzed chromatin motion after inducing replication stress. By treating cells with aphidicolin, DNA synthesis is slowed down or stopped altogether (Vesela et al., 2017). Aphidicolin is a tetracyclic antibiotic isolated from *Nigrospora sphaerica,* which interferes with DNA replication directly by inhibiting DNA polymerases α, ε, and δ (Bambara and Jessee, 1991; Baranovskiy et al., 2014; Byrnes, 1984; Cheng and Kuchta, 1993). Our hypothesis was that it is the loading of replisome components that affects the chromatin motion (LCD). Therefore, we focused on LCD measurements after inhibiting DNA synthesis directly with aphidicolin and characterized the effects on chromatin motion in order to understand the mechanism behind it.

First, we tested in detail the rate and level of inhibition of DNA synthesis with aphidicolin (150 μM) using thymidine analogs (in this case EdU), which get incorporated into newly synthesized DNA and can be detected using click-IT chemistry (Methods). We visualized GFP-PCNA and EdU in fixed cells and performed high-throughput image analysis to characterize the effect of aphidicolin on DNA synthesis inhibition at different timepoints (Methods, Supplementary Figure S4). We observed that DNA synthesis was inhibited minutes after aphidicolin treatment. Using high-content microscopy, we quantified the population of cells actively synthesizing DNA (EdU signal) upon stress and observed that in almost 99 % of the cell population, DNA replication was inhibited within half an hour of aphidicolin incubation (Methods, Figure 5A, Supplementary Figure S4, S5). Subsequent experiments were all performed with these conditions.

**Figure 5:**
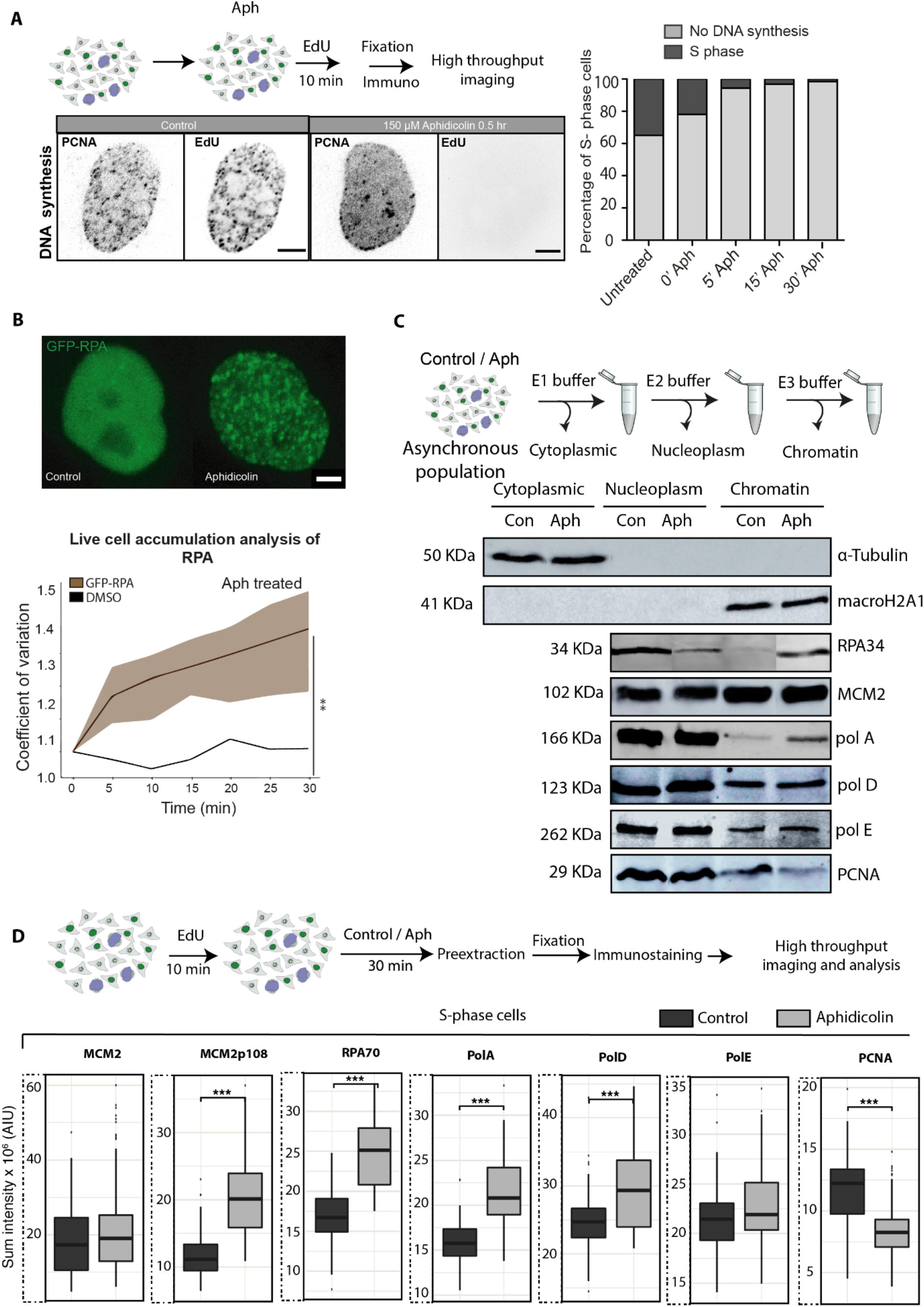
Dissecting the kinetics of replisome components after inhibition of DNA synthesis. **(A)** The use of thymidine nucleotide analogs like 5-ethynyl-2′-deoxyuridine (EdU), which is incorporated into replicating DNA, allows us to estimate the time needed for complete inhibition of DNA synthesis. The representative images show no incorporation of EdU in S-phase cells upon aphidicolin (Aph) treatment for 30’. The plots below the images depict the % of cells with no DNA synthesis as scored by the EdU signal and the corresponding % of cells still replicating DNA (Supplementary Figures S4, S5). **(B)** The line plot shows the live cell accumulation analysis showing the normalized average RPA70 accumulation at replication sites (coefficient of variation ± standard deviation in transparent color) of HeLa cells stably expressing GFP-RPA34 (Supplementary Figures S6, S7, S8). **(C)** Western blots of cytoplasm, nucleoplasm and chromatin fractions of asynchronous population of HeLa cells probed for different replication factors. The western blots shown are cropped from the same replicates for easier visualization without contrast adjustment and the full blots are shown and highlighted in Supplementary Figure S9. **(D)** HeLa cells were pulsed with EdU for 10 min to identify S-phase cells and pre-extracted to detect chromatin bound proteins and different replication factors were detected using immunofluorescence. High-throughput imaging and image analysis were performed (Supplementary Figures S10, S11). Box plots depict the accumulation of the replisome factors indicated at DNA replication sites. Same Y-axis scale plots are shown in Supplementary Figure S11B. The boxplot lower and upper hinges correspond to the first and third quartiles (the 25th and 75th percentiles), the upper whisker extends from the hinge to the largest value no further than 1.5 x IQR from the hinge (where IQR is the interquartile range, or distance between the first and third quartiles). The lower whisker extends from the hinge to the smallest value at most 1.5 x IQR of the hinge. The horizontal line represents the median value. The outliers plotted individually as separate dots outside of the whiskers.*** p < 0.001 by Wilcoxon rank-sum test, for aphidicolin-treated versus control sample. Scale bar: 5 µm.

Secondly, we made use of the above conditions in which DNA synthesis was inhibited, and analyzed the consequences of replication stress on the replisome components and their kinetics. For this purpose, we performed time-lapse microscopy of GFP-PCNA and GFP-RPA34 expressing cells. During active DNA synthesis, the DNA polymerase clamp and processivity factor PCNA is loaded onto the DNA as a trimeric ring and is tightly bound to the DNA (Figure 5A). During aphidicolin treatment though, PCNA dissociated from DNA as shown before (Görisch et al., 2008) (Rausch et al., 2021)(Figure 5A). Aphidicolin treatment does not stop helicase activity and the single stranded DNA binding protein RPA is loaded on the ssDNA after being unwound by the DNA helicase. The more the DNA double helix is unwound, the more RPA loads onto the ssDNA generated (Rausch et al., 2021). For this analysis, we generated a HeLa cell line stably expressing GFP-RPA34 (Supplementary Figure S6). We performed time-lapse microscopy on HeLa GFP-RPA34 cells every 5 minutes for 60 min for both aphidicolin treated and control DMSO treated cells (Supplementary Figure S7A). We observed that RPA accumulated over time on DNA at replication sites in aphidicolin-treated cells but not in the control cells (Supplementary Figures S7). RPA accumulation indicated that the DNA helicase complexes continued unwinding the DNA, which allowed for increasing amounts of RPA to bind and, at the same time, the DNA polymerases were not active displacing the RPA while synthesizing the second (complementary) DNA strand (Görisch et al., 2008). Therefore, we studied the kinetics of accumulation of RPA on chromatin upon DNA synthesis inhibition by quantifying the accumulation of GFP-RPA34 in live cells upon treatment with aphidicolin normalized to DMSO treated cells using the coefficient of variation (Cv), which indicates the amount of RPA protein accumulated over time (Methods, Figure 5B, Supplementary Figure S8). We observed clear accumulation of RPA over time relative to control, showing that the single-strand DNA binding protein accumulates on chromatin. Hence, this indicates that upon stress the DNA helicase remained active unwinding the DNA.

Next, we analyzed the distribution of the helicase subunit MCM2 and its phosphorylated (p108) form along with DNA polymerases α, ε, and δ (Supplementary Table 4) at the chromatin. It has been previously described that the phosphorylated form of MCM2 is the active form for DNA unwinding (Forsburg, 2004; Montagnoli et al., 2006). We predicted from the RPA accumulation that the helicase subunit was present at the replication sites and actively spooling the DNA through after the DNA synthesis inhibition. We first performed western blot analysis of different replication factors from asynchronous populations of HeLa cells after isolating the cytoplasm, nucleoplasm, and chromatin fractions (Methods). We tested the fractionation protocol by blotting the membranes with antibodies to α-tubulin for the cytoplasmic fraction and macro H2A1 histone for the chromatin fraction (Figure 5C). The same fractions were then incubated with antibodies for different replication factors. We observed significant dissociation of PCNA from chromatin and accumulation of RPA on chromatin upon aphidicolin treatment (Figure 5C) consistent with our fixed cell and live cell microscopy analysis. We found no significant changes in MCM2 helicase subunit levels on chromatin and higher levels of phosphorylation of MCM2 upon treatment with aphidicolin (Figure 5C). Lastly, we incubated the blots with antibodies recognizing the catalytic subunits of the DNA polymerases α, δ and ε complexes (Methods, Supplementary Table 4). The DNA polymerases showed a different behavior as compared to the DNA polymerase clamp protein, with DNA polymerase α being enriched on chromatin upon stress, with only minor to no changes being observed for the processive DNA polymerases δ and ε (Figure 5C, 5D). It is of note that both these processive DNA polymerases bind the polymerase clamp PCNA whereas the far less processive DNA polymerase α does not. The full length blots are shown in the Supplementary Figure S9.

We then performed an orthogonal analysis using high-throughput microscopy and image analysis. We labeled cells with EdU for 10 minutes to mark the S-phase cells and treated cells with DMSO/aphidicolin and subsequently performed pre-extraction to remove the unbound fraction of proteins and only detect the chromatin bound proteins. In this manner, we separately quantified accumulation only in S-phase cells and not in populations of cells including all cell cycle stages as in the previous western blot analysis (Figure 5C). The pre-extracted cells were fixed and immunostained for different replisome components (Figure 5D). The cells were imaged using a spinning disk confocal microscopy system (Supplementary Table 5) and image analysis was performed using the KNIME software with custom pipeline to quantify the accumulation/loss of replication factors on chromatin in S-phase cells (Methods, Supplementary Table 9, Supplementary Figures S10, S11). Using the EdU signal, the S-phase cells were selected for the quantitation of chromatin bound replisome components (Figure 5D). We found that PCNA dissociated from chromatin and RPA accumulated on chromatin upon stress in accordance with our previous analysis (Figure 5C). We found no changes in MCM2 helicase subunit but an increase in active MCM2p108 levels upon stress. This is consistent with no new loading of DNA helicases but de novo activation of already loaded helicase complexes (Ge et al., 2007; Ibarra et al., 2008). Finally, we observed significant accumulation of DNA polymerases α and δ on chromatin after aphidicolin treatment. PCNA does not associate with DNA polymerase α, which has a low processivity, but it associates with DNA polymerases ε and δ, which constitute the processive synthetic machinery responsible for most of the duplication of the genome. Hence, it was surprising that these two polymerases remain associated and even load de novo at non-synthetizing replication sites. The increase in DNA polymerase α could lead to the recruitment of alternative polymerase clamp 9-1-1 as reported before (Michael et al., 2000; Van et al., 2010; Yan and Michael, 2009a, 2009b). Several scenarios explaining the different levels of DNA polymerases α and δ upon stress are possible (Supplementary Figure S12): i) multiple polymerase complexes may load within the same Okazaki fragment, which is less likely in view of what is known on DNA replication (Supplementary Figure S12A); ii) multiple Okazaki fragments each with DNA polymerase α and δ within the same replication fork may form on the extended single stranded DNA unwound by the helicase complex (Supplementary Figure S12B); iii) additional replication origins may fire in the proximity of the stalled replication fork, which would explain the increase in both active phosphorylated helicase and DNA polymerases α and δ (Supplementary Figure S12C). Having established the conditions in which DNA synthesis but not DNA unwinding is blocked and concomitantly polymerases are accumulated, we then addressed the consequences for chromatin motion.

### Accumulation of replisome components but not processive DNA synthesis *per se* restricts chromatin motion

To elucidate the roles of the processive DNA synthesis and loading of synthetic machinery in chromatin motion decrease in S-phase, we imaged single cells for PCNA, RPA34, and Cy3-dUTP pre and post aphidicolin treatment (Figure 6A, see also movies 4,5). The PCNA and RPA patterns did not change in G1/G2, whereas in S-phase the PCNA was dissociated from chromatin and RPA was accumulated at the same previously replicating sites (Figure 6A, see also movies 6,7). Chromatin motion analysis was performed on DNA labeled with Cy3-dUTP for both G1/G2 cells and S-phase cells pre and post-treatment with aphidicolin. We observed that chromatin motion was unaffected in G1/G2, which fit with our prediction, as in G1/G2 there is no active DNA synthesis besides possible DNA repair processes on a limited genomic scale (Figure 6B). As hydroxyurea, another DNA synthesis inhibitor, significantly affected chromatin mobility outside of S-phase, we did not further pursue it. Surprisingly, aphidicolin treatment and inhibition of DNA synthesis led to additional decrease in chromatin motion (Figure 6B) and the chromatin became even more constrained than at the proximity of the active replication sites in S-phase cells (see Figure 4C). As quantified above, after aphidicolin treatment, the helicases were still loaded and actively spooling DNA through, whereas the DNA polymerases α and δ albeit not synthesizing DNA accumulated on chromatin at the sites of helicase/RPA accumulation (Figure 5).

**Figure 6:**
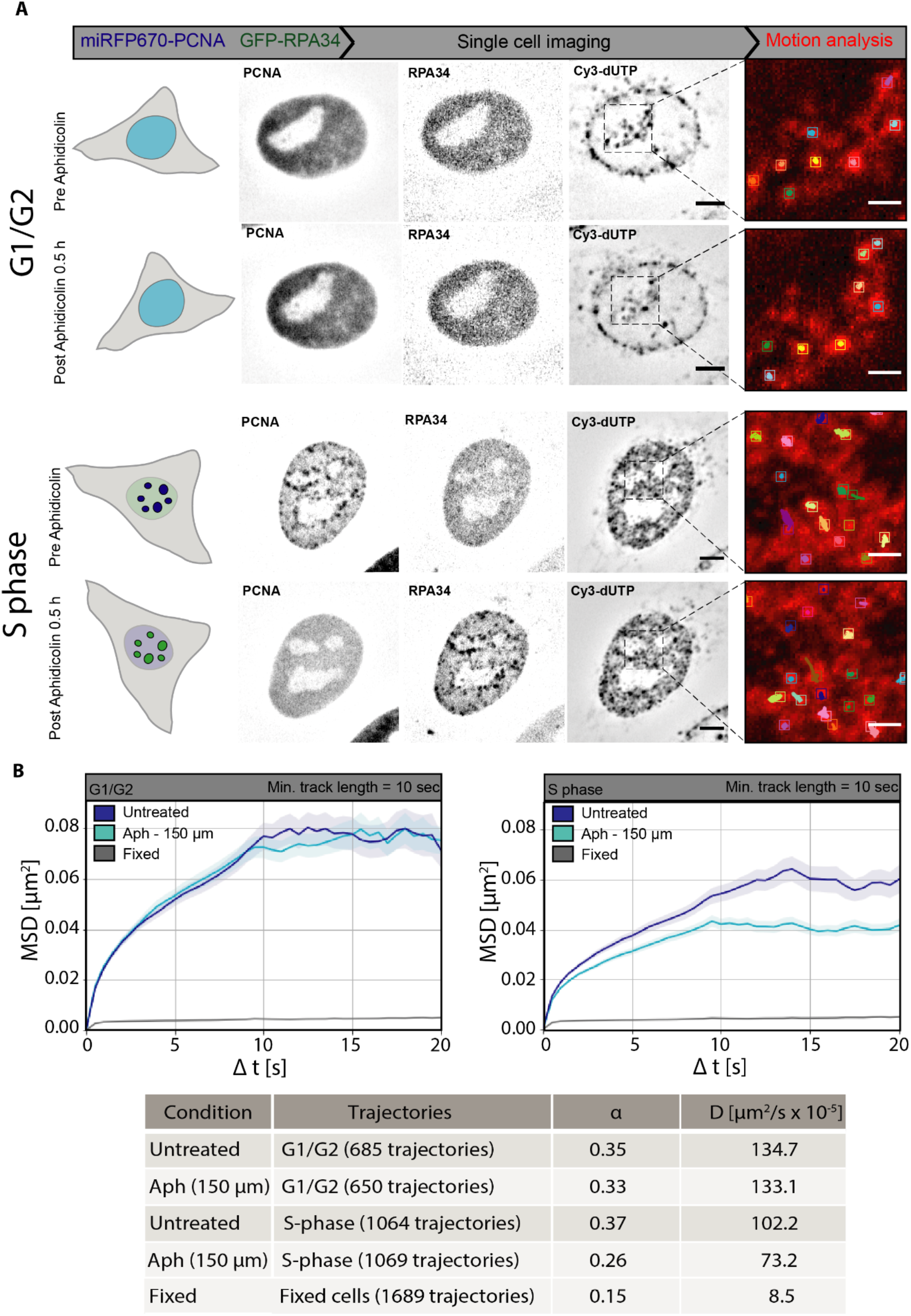
Inhibition of DNA synthesis by aphidicolin further restricts chromatin mobility in S phase but not in G1/G2 cells. **(A)** Representative images of HeLa GFP-RPA34 cells transfected with a construct coding for miRFP670-PCNA and Cy3-dUTP nucleotides for both G1/G2 and S-phase cells pre and post aphidicolin (Aph) treatment. The chromatin foci were imaged using the spinning disk microscope. The image sequences were used to perform motion analysis. The cropped region (black dashed lines) show the motion analysis of chromatin tracks before and after treatment of the same cells (see also movies 6,7). **(B)** Mean square displacement curves over time were plotted for all chromatin tracks for untreated and aphidicolin-treated (150 µM) cells in G1/G2 and S-phase. The error bars are represented in transparent color around the curve. The table below provides the detailed information on number of trajectories per individual sample along with average diffusion rates (µm^2^/s x 10^-5^) and anomalous α coefficient showing subdiffusion. The MSD’s were plotted with error bars (standard deviation) represented in transparent color around the curve. Scale bar: 5 µm. Insets scale bar: 1 µm.

In summary, we propose that the accumulation of the helicase and polymerase complexes on chromatin together with the continuous loading of the single stranded DNA binding protein (RPA) covering the ssDNA strands, stiffens the DNA polymer and restricts its diffusional motion. This study provides new insights on the kinetics of DNA replication proteins loading upon DNA replication stress and elucidates the transient and localized immobilization of chromatin during DNA replication.

## Materials and Methods

### Cells

All cells used were tested and negative for mycoplasma. Human cervical cancer cell line HeLa Kyoto (Erfle et al., 2007) and human normal diploid fibroblasts from lung tissue IMR90 (Nichols et al. 1977) were used in the study. Previously published HeLa Kyoto cells expressing GFP-PCNA (Chagin et al., 2016) fusion protein, were used to monitor cell cycle progression. HeLa Kyoto GFP-RPA34 were generated using the Flp-In recombination system based on the Flp site-specific recombinase (Cat.No.: K6010-01, Invitrogen, Waltham, Massachusetts, USA). The HeLa Kyoto FRTLacZ cells containing a genomically integrated FRT site described earlier (Chagin et al., 2016) were cotransfected with pFRT-B-GRPA34 (Supplementary Table 2) (encoding GFP-RPA34) and pOG44 Flp-recombinase using Neon transfection (Cat.No.: MPK5000, Invitrogen, Waltham, Massachusetts, USA). Four hours after transfection the cell culture medium was exchanged and cells were grown for 48 h and selected with 2.5 mg ml^-1^ blasticidin (Cat.No.: R210-01, Invitrogen, Waltham, Massachusetts, USA). A stable monoclonal line was isolated using blasticidin selection. All cells were maintained in Dulbecco’s modified Eagle medium (DMEM) high glucose (Cat.No.: D6429, Sigma-Aldrich Chemie GmbH, Steinheim, Germany) supplemented with 10% fetal calf serum, 1x glutamine (Cat.No.: G7513, Sigma-Aldrich, St Louis, MO, USA) and 1 µM gentamicin (Cat.No.: G1397, Sigma-Aldrich, St Louis, MO, USA) in a humidified atmosphere with 5% CO2 at 37 °C. Additional experiments confirmed that the transgenic gene product co-localized with the endogenous protein (not shown) and was present at sites of active replication (Supplementary Figure S6). The culture medium was changed every day and cells were split every two days. Cell line characteristics are summarized in Supplementary Table 1.

To block DNA replication, cells were treated with aphidicolin (Aph) (Cat.No.: A0781-1MG, Sigma-Aldrich, St Louis, MO, USA) at final concentration of 150 µM (Supplementary Table 3). Cells were subsequently examined for 30 minutes (aphidicolin) following drug exposure. To confirm that DNA synthesis was inhibited, cells were labeled with 10 μM nucleoside analog 5-ethynyl-2’-deoxyuridine (Cat.No.: 7845.1, ClickIt-EdU cell proliferation assay, Carl Roth, Karlsruhe, Germany) (Supplementary Table 3) in media for 10 minutes to evaluate the extent of replication in control and treated cells (Supplementary Figure S4).

For synchronization of HeLa cells, the cells were seeded on tissue culture dishes at high confluency. Once the cells were confluent, the cells were placed on a shaker for 5 minutes. The detached mitotic cells were collected from the supernatant, and seeded on coverslips. Once the cells were in G1, they were fixed and stained with DAPI for quantification (Supplementary Figure S2B).

### Live cell imaging and replication labeling

For live-cell microscopy, cells were transfected using a Neon transfection system (Cat.No.: MPK5000, Invitrogen, Waltham, Massachusetts, USA). Briefly, the asynchronous population of cells were washed with 1x PBS/EDTA, trypsinized, and collected in a 15 ml tube. The cells were pelleted at 300 x g for 5 minutes. The media was removed and cells were resuspended in 100 µl resuspension buffer R and transferred to a 1.5 ml microcentrifuge tube. Either 15 µg of plasmid DNA or/and 0.5 ul (25 nM) Cy3-dUTP (Cat.No.:ENZ-42501, Enzo Life Sciences, Lörrach, Germany) was added to the cell mixture (Supplementary Tables 2, 3). The Neon^TM^ tip was immersed into the cell mixture and the mixture pipetted taking care to avoid bubbles. The tip was immersed in electrolytic buffer E2 and cells were electroporated (HeLa [voltage - 1005 V, width - 35, pulses - 2], IMR90 [1100 V, width - 30, pulses - 1]). The electroporated mixture was transferred to Ibidi μ-dish chambers (Cat.No.: 80826, Ibidi, Gräfelfing, Germany). Additionally, IMR90 cells were transfected with miRFP670-PCNA plasmid (Supplementary Table 2) to mark the DNA replication sites. After transfection, cells were allowed to attach overnight and were imaged the next day. All imaging was performed at 37 °C with a humidified atmosphere of 5% CO_2_ using an Olympus environmental chamber (spinning disk microscope, Supplementary Table 5).

### Immunofluorescence

For immunofluorescence, cells were fixed with 3.7 % formaldehyde/1x phosphate-buffered saline (PBS) (Cat.No.: F8775, Sigma-Aldrich Chemie GmbH, Steinheim, Germany) for 15 minutes and permeabilized with 0.7 % Triton-X100 in 1x PBS for 20 minutes. All washing steps were performed with PBS-T (1x PBS/0.075 % Tween- 20). For detection of PCNA, cells were further incubated for 5 minutes in ice-cold methanol for antigen retrieval. Blocking (1% bovine serum albumin in 1x PBS) was performed for 30 minutes at room temperature. EdU was detected using the Click-IT assay as described by the manufacturer (1:1000 6-FAM azide or 1:2000 5/6-Sulforhodamine azide; Cat.No.: 7806 and 7776 respectively, Carl Roth, Karlsruhe, Germany). Primary and secondary antibodies were diluted in the blocking buffer and incubated for 1 hour at room temperature with subsequent 3×10 minutes of PBS-T washing. DNA was counterstained with DAPI (4′,6-diamidino-2-phenylindole, 10 μg/ml, Cat.No.: D27802, Sigma-Aldrich Chemie GmbH, Steinheim, Germany) for 10 minutes, and samples were mounted in Vectashield (Cat.No.: VEC-H-1000,Vector Laboratories, Inc., Burlingame,CA, USA). Antibody characteristics are summarized in Supplementary Table 4.

### Western blot and chromatin fractionation

Cells for western blot were washed with 5 ml ice-cold 1x PBS once and 2 ml of ice-cold 1x PBS was added and cells were scraped using a cell scraper. Cells were then centrifuged in a 15 ml tube at 500 x g for 5 minutes. Cells were lysed for total cell lysates for 1 hr at 4 *°*C using the IP lysis buffer with (150 mM NaCl (Cat.No.: 0601.2, Carl Roth, Karlsruhe, Germany), 200 mM TrisCl pH 8 (Cat.No.: A1086.500, Diagonal, Münster, Germany), 5 mM EDTA (Cat.No.: 8040.2, Carl Roth, Karlsruhe, Germany), 0.5 % NP-40 (Cat.No.: 74385, Sigma-Aldrich Chemie GmbH, Steinheim, Germany)) and protease and phosphatase inhibitors (PMSF (Cat.No.: 6367.1, Carl Roth, Karlsruhe, Germany), PepA (Cat.No.: 2936.2, Carl Roth, Karlsruhe, Germany), NaF (Cat.No.: 67414-1ML-F, Sigma-Aldrich Chemie GmbH, Steinheim, Germany), Na_3_VO_4_ (Cat.No.: S6508-10G, Sigma-Aldrich Chemie GmbH, Steinheim, Germany)). Protein fractionation of control and treated samples was performed as described in (Gillotin, 2018). Briefly, equal number of cells were washed with buffer E1 (cytoplasmic fraction) and centrifuged at 1200 x g for 2 min and collected into a new tube. The step was repeated 2 times to remove excess cytoplasmic fraction. The pellet was then washed with buffer E2 (nucleoplasm fraction) and collected into a new tube. The chromatin fraction was isolated with buffer E3 and 1:1000 benzonase for 20 minutes at 25 *°*C.

All lysates was then centrifuged at 13,000 rpm for 20 minutes at 4 *°*C. The supernatant was collected into a new 1.5 ml tube and protein concentration was measured using the bovine serum albumin protein standard assay (Cat.No.: 23208, Thermo Fisher Scientific, Waltham, Massachusetts, USA) according to the manufacturer’s protocol. 10 % SDS-PAGE gel was prepared and 50 μg of protein lysate was loaded along with the protein standard ladder (Cat.No.: P7719S, New England Biolabs, Ipswich, Massachusetts, United States), and electrophoresis was performed for 1.5 hr in ice-cold 1x Laemmli electrophoresis running buffer. Then, the protein was transferred to the 0.2 μm nitrocellulose membrane using a semi-dry transfer system (#1703940, Trans-Blot® SD Semi-Dry Transfer Cell, Bio-Rad, Hercules, CA, USA) for 55 min at 25 V using 1x transfer buffer (Pierce Western Blot Transfer Buffer 10x, Thermo Fisher Scientific, Waltham, Massachusetts, USA). After the transfer, the blotting membrane was incubated in a blocking buffer (5 % low-fat milk in 1x PBS) for 30 minutes. The primary antibodies (Supplementary Table 4) were diluted in blocking buffer to 5 % milk and incubated at 4 *°*C overnight. The next day the membrane was washed 3 times with 1x PBS-T (0.075 %) 10 minutes each. The membrane was then incubated with secondary antibodies (Supplementary Table 4) for 1 hr at room temperature. The membrane was washed again with 1x PBS-T (0.075 %) 3 times 10 minutes each and incubated with 1:1 ECL chemiluminescence solution (Clarity Western ECL, #170-5061, Bio-Rad Laboratories, Hercules, CA, USA). Signal was detected using an Amersham AI600 imager (Supplementary Table 5).

### Microscopy

Live cell imaging for chromatin mobility measurements were performed using the PerkinElmer UltraVIEW VoX system with a 60x/1.45 numerical aperture plan-apochromatic oil immersion objective. Cy3 and GFP were excited sequentially using 543 nm and 488 nm solid-state diode laser lines to minimize crosstalk. The standard protocol for examining chromatin mobility in Cy3-dUTP labeled nuclei was as follows: first, a reference image comprising the miRFP670/GFP-PCNA, Cy3-dUTP, and the phase-contrast signal was collected from a single focal plane corresponding to the middle of the nucleus. This image demarcated the nuclear boundary, provided cell cycle information, and, in the case of S-phase cells, allowed us to correlate the positions of Cy3-dUTP foci with sites of DNA replication. Second, while maintaining the same focal plane, a time series (30-60 seconds) at a frame rate of 500 ms was captured. To maximize the temporal resolution, the time series consisted solely of the Cy3-dUTP channel and a PCNA reference frame at the beginning to obtain information on the cell cycle stage.

Multiple point time-lapse microscopy was performed using the multi-time option available in the spinning disk Volocity 6.3 software to image the chromatin (Cy3-dUTP) of the same cells pre and post treatment of aphidicolin. To minimize photo-toxicity over the course of the experiment, transmitted light contrast imaging was used to focus the cells. Live cell imaging was performed by following cells through the cell cycle and G1 and G2 stages were classified based on the previous cell cycle stage.

For the inhibition experiments (aphidicolin) different cells/points were chosen using the multipoint function of the Perkin Elmer spinning disk, and image sequences before the treatment were acquired. The reference image consisted of GFP-RPA34, Cy3-dUTP, and miRFP670-PCNA using 488 nm, 561 nm, and 640 nm solid-state diode lasers, respectively. After acquiring the reference images the media containing the small molecule inhibitor was added to cells on the microscope for the required time and after treatment image sequences were acquired for analysis of chromatin motion.

High-throughput imaging was performed using the 40x/0.95 numerical aperture air objective of the PerkinElmer Operetta system. We used different filters (excitation/emission: 360/400, 460/490, 560/580) to image DAPI, EdU, and different replication proteins.(Figure 5, Supplementary Figure S5, Supplementary Table 5).

### Quantification of DNA synthesis inhibition

The high-throughput images were used to quantify the percentage of cells with inhibition of DNA synthesis upon aphidicolin treatment. A minimum of 100 fields with around 2000-5000 cells were acquired in all channels. The images were then analyzed using the PerkinElmer Harmony software. The steps in brief (Supplementary Figure S4, S5) include segmentation of nuclei using cell types specific parameters like the diameter, splitting coefficient, and intensity threshold. The segmentation was then validated by visually checking it in randomly selected regions. Once the nuclei were segmented, cells touching the border were omitted. The intensity values with mean, median, standard deviation, and the sum of the intensities were obtained for individual cells. The datasheets were then imported to R and plots were generated. EdU signal was used to identify the population of cells actively replicating upon Aph treatment (Figure 5A). The background intensity for EdU staining was determined using a negative control which was not treated with EdU but stained. The cells showing a mean intensity greater than the background intensity were separated into an EdU positive population and plotted.

### DNA quantification of labeled chromatin

DNA quantification of the labeled foci was done by automated image analysis. Image sequences with labeled chromatin were acquired on a Ultra-View VoX spinning disk microscope, using a 60x objective (Figure 2, Supplementary Figure S1). For segmentation of replication foci, we used the protocol originally described in (Chagin et al., 2016, 2015). The channels comprising DAPI replication foci signals were imported into the software Perkin Elmer Volocity 6.3 and converted into volumes. The pixel dimensions of the images were set to the specifications for the spinning disk (x/y: 0.066 μm and z: 0.3 μm). The following processing steps were applied: Find objects (“nucleus”) using the DAPI channel, method “Intensity” (set manually to the optimal value), use fill holes in object/dilate/erode until the object optimally fits the nucleus, exclude objects by size < 500 μm^3^. Find objects using the label channel, method “Intensity” (lower limit: 1, upper limit: 65535), separate touching objects, exclude “foci’’ not touching “nucleus”. Using the detected foci, the DNA content of foci was determined via the sum of intensities in the DAPI channel and the genome size of the cell type (Figure 2, Supplementary Figure S1).

### Automated tracking of chromatin structures in time-lapse videos

The motility of fluorescently labeled chromatin structures in live-cell fluorescence microscopy images was quantified within manually segmented single nuclei. The background image intensity was adjusted for each image sequence to the computed mean intensity value over all time points within a manually selected region of interest (ROI) of the background. Automatic tracking of multiple fluorescently labeled chromatin structures was performed using a probabilistic particle tracking approach, which is based on Bayesian filtering and multi-sensor data fusion (Ritter et al., 2021). This approach combines Kalman filtering with particle filtering and integrates multiple measurements by separate sensor models and sequential multi-sensor data fusion. Detection-based and prediction-based measurements are obtained by elliptical sampling (Godinez and Rohr, 2015), and the separate sensor models allow taking into account different uncertainties. In addition, motion information based on displacements from past time points is exploited and integrated in the cost function for correspondence finding. Chromatin structures are detected by the spot-enhancing filter (SEF) (Sage et al., 2005) which consists of a Laplacian-of-Gaussian (LoG) filter followed by thresholding the filtered image and determination of local maxima. The threshold is automatically determined by the mean of the absolute values of the filtered image plus a factor times the standard deviation. We used the same threshold factor for all images of an image sequence (Supplementary Figure S2).

### Chromatin motility analysis

Based on the computed trajectories, the motility of chromatin structures was analyzed and the motion type was determined for different cell cycle stages along with active replication sites, and inhibition of DNA synthesis with aphidicolin. We performed a mean square displacement (MSD) analysis (Saxton, 1997) and computed the MSD as a function of the time interval Δt for each trajectory (Supplementary Figure S2). The MSD curves for all trajectories with a minimum time duration of 10 s (corresponding to 20 time steps) under one condition were averaged. We considered only the trajectories with a time duration larger than the minimum time duration which improved the accuracy of the motility analysis. We fitted the anomalous diffusion model to the calculated MSD values to obtain the anomalous diffusion coefficient α. The motion was classified into confined diffusion, obstructed diffusion, and normal diffusion (Bacher et al., 2004). To determine the diffusion coefficient D [μm²s^-1^], the diffusion model was fitted to the MSD values. In case of IMR90 cells, affine image registration was performed using the method in (Celikay et al., 2022) to address the stronger cell movement compared to HeLa cells.

### Center to center distance / Proximity analysis

Automatic proximity analysis of chromatin and PCNA was performed using the computed trajectories of chromatin structures and detected sites of active DNA synthesis represented by fluorescently labeled PCNA. Only trajectories of chromatin structures present at the first time point of an image sequence and with a minimum time duration of 10 s (corresponding to 20 time steps) were considered. PCNA foci were automatically detected in the fluorescence microscopy images by the SEF filter (Supplementary Figure S2). For each PCNA image, a single nucleus was manually segmented and the background intensity was adjusted to the computed mean intensity value within a manually selected ROI of the background. Proximity was determined for the first time point of the trajectory of a chromatin structure and detected PCNA foci using a graph-based *k*-d-tree approach (Bentley, 1975). Due to the *k*-d-tree structure, this approach allows efficient computation of the nearest neighbor query based on the Euclidean distance between foci in the chromatin and PCNA channel. If a chromatin structure at the first time point of the image sequence has a nearest PCNA neighbor within a maximum distance, the trajectory of a chromatin structure is considered within center to center distance (CCD). Otherwise, the trajectory is considered outside the center to center distance (CCD) (Supplementary Figure S3).

### Accumulation analysis

To analyze the focal RPA accumulation upon DMSO/aphidicolin treatment, cell nuclei were segmented using the Volocity software (Version 6.3, Perkin Elmer). The GFP-RPA34 signal was segmented before and after treatment of the same cell in the live experiments and plotted over time after DMSO and drug treatment. The GFP-RPA intensities were measured and the coefficient of variation cV = **σ**/µ, with **σ** = standard deviation and µ = mean) was calculated for all time points (Supplementary Figures S7, S8). All values were normalized to the DMSO treatment *c*V = *c*V(*tp*x)/*c*V(*tp*0) with *tp*x: any given time point imaged, *tp*0: pretreatment time point).and plotted using RStudio (Supplementary Table 9).

### High throughput image analysis of replisome components

The images from the Nikon crest Ti2 system were analyzed with the custom made image analysis pipeline in KNIME Analytics Platform. The image analysis pipeline was constructed as follows (Supplementary Figures S10, S11). Briefly, the channels were separated. The DAPI channel was used for the nuclei segmentation. Nuclei were segmented based on manually chosen intensity threshold, the Watershed Transform was applied next to separate the close-positioned nuclei. The segmented nuclei were converted into a mask with each nucleus DAPI intensity and texture features recorded. The nuclei population was further thresholded by nucleus area and circularity to eliminate segmentation artifacts. The EdU and replication protein channels were subjected to foci segmentation based on a wavelet transform algorithm. The algorithm parameters were selected individually for each type of the replication protein and maintained the same between the control and treated samples. The nuclear mask and EdU foci/replication protein foci masks were overlaid to filter only the foci inside the nuclear areas. The EdU foci/replication protein foci intensity parameters (total focus intensity, mean focus intensity), area and foci number per nucleus were exported as XLSX files for further analysis. The data was analyzed in R Studio (https://posit.co/download/rstudio-desktop/). First, the S phase cell population was identified by the number of EdU foci per nucleus. The EdU foci number threshold was set as 50 for the cells in control samples, and 55 for aphidicolin-treated samples among all datasets. The nuclei in S phase were next analyzed for their replication protein accumulation. The total levels of the replication proteins was plotted as boxplots, *** p < 0.001 by Wilcoxon rank-sum test, for aphidicolin-treated vs. control sample.

## Acknowledgments

We thank Anne Lehmkuhl and Diana Imblan for their excellent technical support. We are thankful to Alexander Rapp, Hector Romero, and Gaudenz Danuser for their valuable suggestions. We also thank Argyris Papantonis (Georg-August-Universität Göttingen, Germany) for providing the IMR90 cells.

## Data Availability Statement

All data are available from the OMERO open microscopy environment public repository http://cc-omero.bio.tutudarmstadt.de/webclient/userdata/?experimenter=-1 (https://doi.org/10.48328/tudatalib-873). All renewable biological materials will be made available upon request from the corresponding author M. Cristina Cardoso.

## Author contributions

M.K.P., C.R., V.O.C. and M.C.C. designed the study and wrote the manuscript. K.R., and H.L., discussed the project and contributed to the manuscript writing. M.K.P., C.R., V.O.C., J.M., K.C., J.H.S., D.L., K.K., P.P. and A.K.S contributed to experiments, data analysis and figure preparation. All authors reviewed the manuscript.

## SUPPLEMENTARY TABLES AND FIGURES

**Supplementary table 1:**
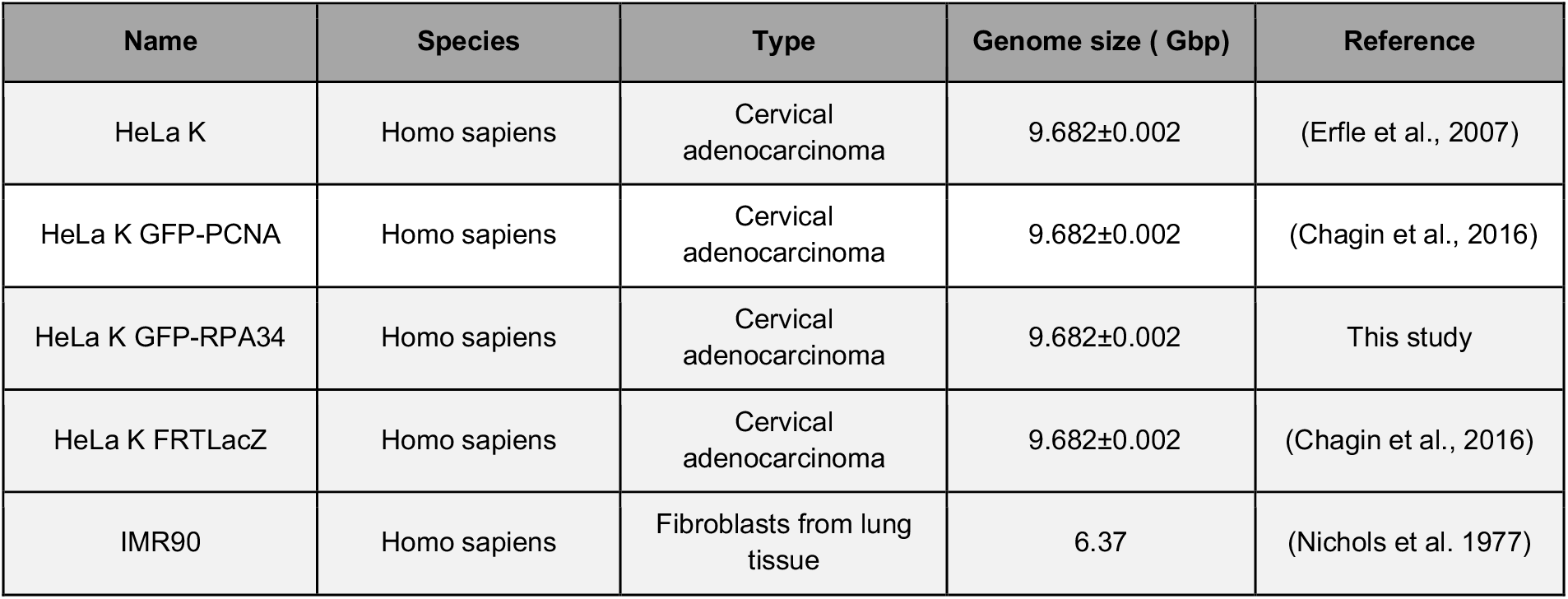
Cell line characteristics

**Supplementary table 2:** Plasmid characteristics

| Name | pc number | Fluorophore | Protein of Interest | Promoter | References |
| --- | --- | --- | --- | --- | --- |
| pmiRFP670-PCNA | 3385 | miRFP670 | Human PCNA | CMV | (Rausch et al., 2021) |
| pFRT-B-GRPA | 1232 | GFP | Human RPA34 | EF1α | This study |
| pFRT-B-GPCNA | 1274 | GFP | Human PCNA | EF1α | Chagin et al., 2016) |

\* pc: plasmid collection.

**Supplementary table 3:**
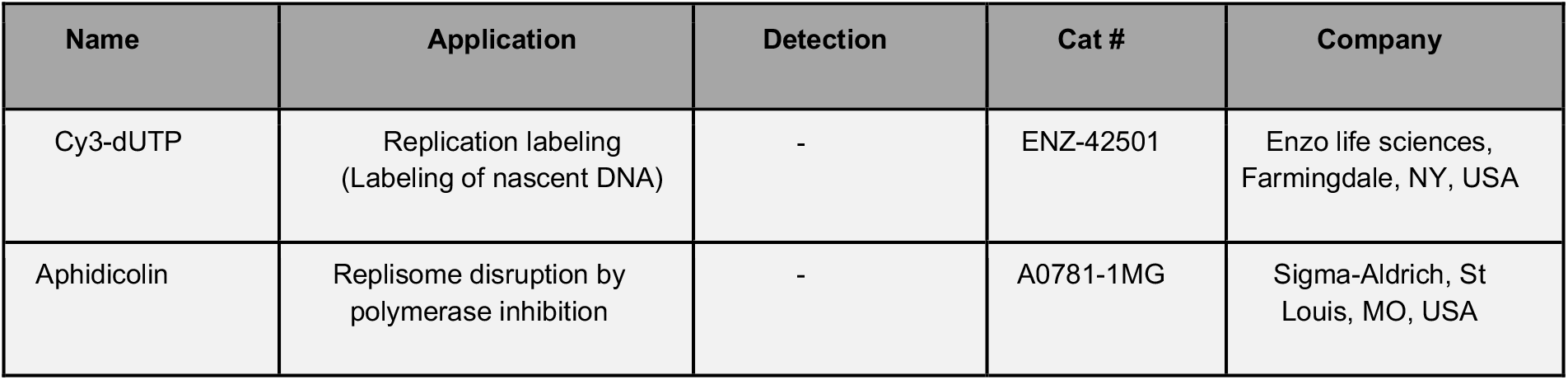
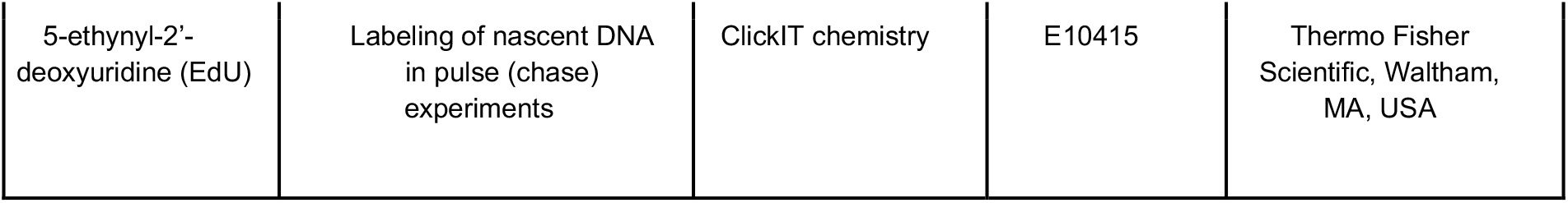
Nucleotide and chemical characteristics

| Name | Application | Detection | Cat # | Company |
| --- | --- | --- | --- | --- |
| Cy3-dUTP | Replication labeling<br>(Labeling of nascent DNA) | - | ENZ-42501 | Enzo life sciences,<br>Farmingdale, NY, USA |
| Aphidicolin | Replisome disruption by<br>polymerase inhibition | - | A0781-1MG | Sigma-Aldrich, St<br>Louis, MO, USA |
| 5-ethynyl-2'-deoxyuridine (EdU) | Labeling of nascent DNA in pulse (chase) experiments | ClickIT chemistry | E10415 | Thermo Fisher Scientific, Waltham, MA, USA |

**Supplementary table 4:** Primary and secondary antibody characteristics

| Reactivity | Host | Clonality | Dilution | Application | Cat / Clone<sup>#</sup> | Company / References |
| --- | --- | --- | --- | --- | --- | --- |
| anti RPA34 | Mouse | Monoclonal | 1:2 | IF, WB | 9H8H4<sup>#</sup> | Gift from Mark Kenny/J. Hurwitz (Kenny et al., 1990) |
| anti RPA70A | Mouse | Monoclonal | 1:2 | IF, WB | 7G9E3<sup>#</sup> | Gift from Mark Kenny/J. Hurwitz (Kenny et al., 1990) |
| Anti MCM2 | Rabbit | Monoclonal | 1:5000 IF, 1:10000 WB | IF, WB | ab 108935/ EPR4120 | Abcam, Cambridge, United Kingdom |
| anti MCM2pS108 | Rabbit | Monoclonal | 1:1000 | IF, WB | 3267-1 | Epitomics, Burlingame, CA, United States |
| anti pol Alpha | Mouse | Monoclonal | Undiluted | IF, WB | SJK-287-38<sup>#</sup> | ATCC (Tanaka et al., 1982) |
| anti pol Delta | Mouse | Monoclonal | 1:500 | IF, WB | 610972 | BD biosciences, New Jersey, USA (Li et al., 2016) |
| anti pol Epsilon | Rabbit | Polyclonal | 1:500 | IF, WB | GTX132100 | GeneTex, Irvine, California, United States |
| anti PCNA | Mouse | Monoclonal | 1:100 | IF*, WB | M0879 / PC10<sup>#</sup> | Dako, Hamburg, Germany (Waseem and Lane, 1990) |
| anti Histone H3 | Rat | Monoclonal | 1:250 | WB | 61647/ 1C8B2<sup>#</sup> | Active Motif, California, USA |
| anti MacroH2A1 | Rabbit | Polyclonal | 1:1000 | WB | 07-219 | Active Motif, California, USA |
| anti GFP | Rat | Monoclonal | 1:1000 | WB | 3H9# | Chromotek 3H9,<br>Planegg-Martinsried,<br>Germany |
| Anti tubulin alpha | Mouse | Monoclonal | 1:5000 | WB | clone DM1A#/<br>T9026 | Sigma,<br>Missouri,<br>United States |
| anti-mouse<br>IgG Cy3 | Donkey | Polyclonal | 1:800 | IF<br>(fluorescent<br>secondary) | 715-165-151 | The Jackson<br>Laboratory, Bar<br>Harbor, ME, USA |
| anti-rabbit<br>IgG Cy3 | Donkey | Polyclonal | 1:800 | IF<br>(fluorescent<br>secondary) | 711-165-152 | The Jackson<br>Laboratory, Bar<br>Harbor, ME, USA |
| anti-mouse<br>IgG Cy5 | Donkey | Polyclonal | 1:800 | F (fluorescent<br>secondary) | 715-175-150 | The Jackson<br>Laboratory, Bar<br>Harbor, ME, USA |
| anti-rabbit<br>IgG Cy5 | Donkey | Polyclonal | 1:800 | F (fluorescent<br>secondary) | 711-175-152 | The Jackson<br>Laboratory, Bar<br>Harbor, ME, USA |
| anti-mouse<br>IgG AF488 | Goat | Polyclonal | 1:800 | IF<br>(fluorescent<br>secondary) | 2120125 | Invitrogen,<br>Waltham,<br>Massachusetts, USA |
| anti-rabbit<br>IgG AF488 | Donkey | Polyclonal | 1:800 | IF<br>(fluorescent<br>secondary) | A11034 | Invitrogen,<br>Waltham,<br>Massachusetts, USA |
| anti rat IgG HRP | Goat | Polyclonal | 1:5000 | WB<br>(HRP<br>secondary) | 112-035-068 | The Jackson<br>Laboratory, Bar<br>Harbor, ME, USA |
| anti mouse IgG HRP | Sheep | Polyclonal | 1:5000 | WB<br>(HRP<br>secondary) | NA931 | Amersham<br>pharmacia,<br>Amersham, United<br>Kingdom |

\*Methanol treatment required #Clone number

**Supplementary table 5:** Imaging systems characteristics

| Microscope/<br>Company | Lasers/lamps | Filters (ex. &<br>em. [nm])* | Objectives/<br>lenses | Detection<br>system | Incubation<br>system | Application |
| --- | --- | --- | --- | --- | --- | --- |
| Ultra-View<br>VoX<br>spinning disk<br>microscope/<br>PerkinElmer<br>Life<br>Sciences,<br>UK | solid state<br>diode lasers<br>(405 nm,<br>488 nm,<br>561 nm,<br>640 nm) | 405/488/56<br>8/640**<br>405: 415–475<br>488: 505–549<br>561: 580–650<br>640: 664–754 | oil immersion<br>60x Plan-Apochromat<br>(NA 1.45) | cooled 14-bit<br>Hamamatsu®<br>C9100-50<br>EMCCD | closed live-cell<br>microscopy<br>chamber<br>(ACU control,<br>Olympus) for<br>time-lapse<br>microscopy | time-lapse<br>microscopy<br>& confocal z-<br>stack<br>imaging |
| Widefield<br>microscope<br>Axiovert<br>200 /Zeiss,<br>Germany | HBO100<br>mercury<br>lamp | 488: 473-491 &<br>506-534<br>561: 550-580 &<br>590-650<br>640: 590-650<br>& 663-738 | oil immersion<br>63x Plan-Apochromat<br>(NA 1.4) | 12-bit<br>AxioCam<br>mRM | - | Multi channel<br>wide-field<br>imaging |
| Leica SP5 II<br>confocal<br>microscope<br>/Wetzelar,<br>Germany | 405 nm diode 488<br>nm Argon, 561<br>nm DPSS, 633<br>nm HeNe | AOBS beam<br>splitter | HCX PL APO<br>63x / 1.4-0.6 oil<br>lambda blue &<br>HCX PL APO<br>100x (NA 1.44)<br>oil Corr CS | 2 HyD Hybrid<br>Detectors | - | confocal z-<br>stack<br>imaging |
| Amersham Al600<br>imager | Chemiluminesce<br>nce, UV<br>transillumination | - | large aperture<br>FUJINON™<br>f/0.85 43 mm | 16-bit Peltier<br>cooled Fujifilm<br>Super CCD | - | Western<br>blots and<br>DNA agarose<br>gels |
| Operetta high<br>throughput<br>imaging/<br>PerkinElmer Life<br>Sciences, UK | Xenon fiber-optic<br>light source, 300<br>W, 360 – 640 nm<br>continuous<br>spectrum LED<br>light source for<br>transmission<br>mode | ex:360/400,<br>460/490,<br>560/580<br>em: 410/480,<br>500/550,<br>560/630 | 20x or 40x air<br>(0.45 NA and<br>0.95 NA) long<br>WD*** | 14 bit<br>Jenoptik<br>firecamj203<br>Sony Chip<br>ICX285<br>cooled 20°C<br>below<br>environment | - | high<br>throughput,<br>high content<br>imaging and<br>image<br>analysis |
| Nikon TiE2 inverted with crest spinning disk unit/ Nikon, Japan | SPECTRA X light engine<br>395/25 nm with 295 mW<br>440/20 nm with 256 mW<br>470/24 nm with 196 mW<br>510/25 nm with 62 mW<br>540/30 nm with 231 mW<br>550/15 nm with 260 mW<br>575/25 nm with 310 mW | LED-DA/FI/TR/Cy5-4X-B<br>Quadbandpasse x:390/18, 475/35, 535/50<br>em:460/60, 530/43, 580LP | 40x air (0.95 NA) & 250 µm WD*** | Cooled Nikon Qi2 camera and 16.25 megapixel sCMOS sensor. readout noise is: 2.2. electron | - | high throughput, high content imaging and image analysis |

\* ex.: excitation & em.: emission, \*\* dichroic specification, \*\*\* WD: working distance.

**Supplementary table 6:** Data description for DNA quantification

| Name | Cell stage | DAPI SUM | Correction factor (C) | Corrected genome size (GSxC) Gbp |
| --- | --- | --- | --- | --- |
| Cell1_crop_DAPI_Cy3dUTP_HeLa | SE | 6.55E+08 | 1.05 | 10.185 |
| Cell2_crop_DAPI_Cy3dUTP_HeLa | SE | 8.49E+08 | 1.05 | 10.185 |
| Cell3_crop_DAPI_Cy3dUTP_HeLa | SE | 9.68E+08 | 1.05 | 10.185 |
| Cell4_crop_DAPI_Cy3dUTP_HeLa | SM | 1.14E+09 | 1.25 | 12.125 |
| Cell5_crop_DAPI_Cy3dUTP_HeLa | SL | 1.15E+09 | 1.77 | 17.169 |
| Cell6_crop_DAPI_Cy3dUTP_HeLa | SE | 8.29E+08 | 1.05 | 10.185 |
| Cell7_crop_DAPI_Cy3dUTP_HeLa | SE | 9.49E+08 | 1.05 | 10.185 |
| Cell8_crop_DAPI_Cy3dUTP_HeLa | SM | 1.01E+09 | 1.25 | 12.125 |
| Cell9_crop_DAPI_Cy3dUTP_HeLa | SM | 1.17E+09 | 1.25 | 12.125 |
| Cell10_crop_DAPI_Cy3dUTP_HeLa | SM | 1.00E+09 | 1.25 | 12.125 |
| Cell11_crop_DAPI_Cy3dUTP_HeLa | SL | 1.34E+09 | 1.77 | 17.169 |
| Cell12_crop_DAPI_Cy3dUTP_HeLa | SM | 1.06E+09 | 1.25 | 12.125 |
| Cell13_crop_DAPI_Cy3dUTP_HeLa | SL | 1.49E+09 | 1.77 | 17.169 |
| Cell14_crop_DAPI_Cy3dUTP_HeLa | SE | 8.71E+08 | 1.05 | 10.185 |
| Cell15_crop_DAPI_Cy3dUTP_HeLa | SL | 1.18E+09 | 1.77 | 17.169 |
| Cell16_crop_DAPI_Cy3dUTP_HeLa | SL | 1.48E+09 | 1.77 | 17.169 |
| Cell17_crop_DAPI_Cy3dUTP_HeLa | SL | 1.09E+09 | 1.77 | 17.169 |
| Cell18_crop_DAPI_Cy3dUTP_HeLa | SL | 1.07E+09 | 1.77 | 17.169 |
| Cell19_crop_DAPI_Cy3dUTP_HeLa | SL | 1.17E+09 | 1.77 | 17.169 |
| Cell20_crop_DAPI_Cy3dUTP_HeLa | SM | 1.09E+09 | 1.25 | 12.125 |
| Cell21_crop_DAPI_Cy3dUTP_HeLa | SL | 1.02E+09 | 1.77 | 17.169 |
| Cell22_crop_DAPI_Cy3dUTP_HeLa | SL | 1.02E+09 | 1.77 | 17.169 |
| Cell23_crop_DAPI_Cy3dUTP_HeLa | SE | 9.33E+08 | 1.05 | 10.185 |
| Cell24_crop_DAPI_Cy3dUTP_HeLa | SE | 8.90E+08 | 1.05 | 10.185 |
| Cell25_crop_DAPI_Cy3dUTP_HeLa | SE | 7.67E+08 | 1.05 | 10.185 |
| Cell26_crop_DAPI_Cy3dUTP_HeLa | SE | 8.39E+08 | 1.05 | 10.185 |
| Cell27_crop_DAPI_Cy3dUTP_HeLa | SM | 1.04E+09 | 1.25 | 12.125 |
| Cell28_crop_DAPI_Cy3dUTP_HeLa | SM | 1.01E+09 | 1.25 | 12.125 |
| Cell29_crop_DAPI_Cy3dUTP_HeLa | SM | 1.05E+09 | 1.25 | 12.125 |
| Cell30_crop_DAPI_Cy3dUTP_HeLa | SM | 1.02E+09 | 1.25 | 12.125 |
| Cell1_DAPI_Cy3dUTP_IMR90 | - | 1.08E+09 | - | 6.37 |
| Cell2_DAPI_Cy3dUTP_IMR90 | - | 5.04E+08 | - | 6.37 |
| Cell3_DAPI_Cy3dUTP_IMR90 | - | 1.09E+09 | - | 6.37 |
| Cell4_DAPI_Cy3dUTP_IMR90 | - | 8.72E+08 | - | 6.37 |
| Cell5_DAPI_Cy3dUTP_IMR90 | - | 1.08E+09 | - | 6.37 |
| Cell6_DAPI_Cy3dUTP_IMR90 | - | 5.39E+08 | - | 6.37 |
| Cell7_DAPI_Cy3dUTP_IMR90 | - | 6.33E+08 | - | 6.37 |
| Cell8_DAPI_Cy3dUTP_IMR90 | - | 5.52E+08 | - | 6.37 |
| Cell9_DAPI_Cy3dUTP_IMR90 | - | 6.20E+08 | - | 6.37 |
| Cell10_DAPI_Cy3dUTP_IMR90 | - | 6.68E+08 | - | 6.37 |
| Cell13_DAPI_Cy3dUTP_IMR90 | - | 7.43E+08 | - | 6.37 |
| Cell14_DAPI_Cy3dUTP_IMR90 | - | 6.53E+08 | - | 6.37 |
| Cell15_DAPI_Cy3dUTP_IMR90 | - | 7.25E+08 | - | 6.37 |
| Cell16_DAPI_Cy3dUTP_IMR90 | - | 4.41E+08 | - | 6.37 |
| Cell17_DAPI_Cy3dUTP_IMR90 | - | 1.29E+09 | - | 6.37 |
| Cell18_DAPI_Cy3dUTP_IMR90 | - | 7.51E+08 | - | 6.37 |

**Supplementary table 7:** Data description for 2D confocal fixed images

| Name | Cell stage | Time (s) | Channels* | Pixel size (nm) | frame rate (ms) |
| --- | --- | --- | --- | --- | --- |
| Cell1_HeLa_fixedcells.tif | SE | 40 | 3 | 120 | 500 |
| Cell2_HeLa_fixedcells.tif | SE | 40 | 3 | 120 | 500 |
| Cell3_HeLa_fixedcells.tif | SE | 40 | 3 | 120 | 500 |
| Cell4_HeLa_fixedcells.tif | SM | 40 | 3 | 120 | 500 |
| Cell5_HeLa_fixedcells.tif | SM | 40 | 3 | 120 | 500 |
| Cell6_HeLa_fixedcells.tif | SM | 40 | 3 | 120 | 500 |
| Cell7_HeLa_fixedcells.tif | SM | 40 | 3 | 120 | 500 |
| Cell8_HeLa_fixedcells.tif | SL | 40 | 3 | 120 | 500 |
| Cell9_HeLa_fixedcells.tif | SL | 40 | 3 | 120 | 500 |
| Cell10_HeLa_fixedcells.tif | SL | 40 | 3 | 120 | 500 |
| Cell11_HeLa_fixedcells.tif | SE | 40 | 3 | 120 | 500 |
| Cell12_HeLa_fixedcells.tif | SE | 40 | 3 | 120 | 500 |
| Cell13_HeLa_fixedcells.tif | SE | 40 | 3 | 120 | 500 |
| Cell14_HeLa_fixedcells.tif | SL | 40 | 3 | 120 | 500 |
| Cell15_HeLa_fixedcells.tif | SE | 40 | 3 | 120 | 500 |
| Cell16_HeLa_fixedcells.tif | SE | 40 | 3 | 120 | 500 |
| Cell17_HeLa_fixedcells.tif | SL | 40 | 3 | 120 | 500 |
| Cell18_HeLa_fixedcells.tif | G2 | 40 | 3 | 120 | 500 |
| Cell19_HeLa_fixedcells.tif | G1 | 40 | 3 | 120 | 500 |
| Cell20_HeLa_fixedcells.tif | SM | 40 | 3 | 120 | 500 |
| Cell21_HeLa_fixedcells.tif | SE | 40 | 3 | 120 | 500 |
| Cell22_HeLa_fixedcells.tif | SE | 40 | 3 | 120 | 500 |
| Cell23_HeLa_fixedcells.tif | SL | 40 | 3 | 120 | 500 |
| Cell24_HeLa_fixedcells.tif | SM | 40 | 3 | 120 | 500 |
| Cell25_HeLa_fixedcells.tif | SM | 40 | 3 | 120 | 500 |
| Cell26_HeLa_fixedcells.tif | SE | 40 | 3 | 120 | 500 |
| Cell27_HeLa_fixedcells.tif | SE | 40 | 3 | 120 | 500 |
| Cell28_HeLa_fixedcells.tif | SE | 40 | 3 | 120 | 500 |
| Cell29_HeLa_fixedcells.tif | SM | 40 | 3 | 120 | 500 |
| Cell30_HeLa_fixedcells.tif | SE | 40 | 3 | 120 | 500 |
| Cell31_HeLa_fixedcells.tif | SL | 40 | 3 | 120 | 500 |
| Cell32_HeLa_fixedcells.tif | SE | 40 | 3 | 120 | 500 |
| Cell33_HeLa_fixedcells.tif | SE | 40 | 3 | 120 | 500 |
| Cell34_HeLa_fixedcells.tif | SE | 40 | 3 | 120 | 500 |
| Cell35_HeLa_fixedcells.tif | SE | 40 | 3 | 120 | 500 |
| Cell1_IMR90_fixed.tif | - | 40 | 3 | 120 | 500 |
| Cell2_IMR90_fixed.tif | - | 40 | 3 | 120 | 500 |
| Cell3_IMR90_fixed.tif | - | 40 | 3 | 120 | 500 |
| Cell4_IMR90_fixed.tif | - | 40 | 3 | 120 | 500 |
| Cell5_IMR90_fixed.tif | - | 40 | 3 | 120 | 500 |
| Cell6_IMR90_fixed.tif | - | 40 | 3 | 120 | 500 |
| Cell7_IMR90_fixed.tif | - | 40 | 3 | 120 | 500 |
| Cell8_IMR90_fixed.tif | - | 40 | 3 | 120 | 500 |
| Cell9_IMR90_fixed.tif | - | 40 | 3 | 120 | 500 |
| Cell10_IMR90_fixed.tif | - | 40 | 3 | 120 | 500 |
| Cell11_IMR90_fixed.tif | - | 40 | 3 | 120 | 500 |
| Cell12_IMR90_fixed.tif | - | 40 | 3 | 120 | 500 |
| Cell13_IMR90_fixed.tif | - | 40 | 3 | 120 | 500 |
| Cell14_IMR90_fixed.tif | - | 40 | 3 | 120 | 500 |
| Cell15_IMR90_fixed.tif | - | 40 | 3 | 120 | 500 |
| Cell16_IMR90_fixed.tif | - | 40 | 3 | 120 | 500 |
| Cell17_IMR90_fixed.tif | - | 40 | 3 | 120 | 500 |
| Cell18_IMR90_fixed.tif | - | 40 | 3 | 120 | 500 |

**Supplementary table 8:** Data description for 2D confocal live images

| Name | Cell stage | Time (s) | Exposure time (ms) | Channels* | Frame rate (ms) |
| --- | --- | --- | --- | --- | --- |
| n001_G_Aph_4_AT_c01.tif | G1 | 40 | 300 | 1 | 500 |
| n001_G_Aph_4_AT_c02.tif | G1 | - | 500 | 3 | - |
| n001_G_Aph_4_BT_c01.tif | G1 | 40 | 300 | 1 | 500 |
| n001_G_Aph_4_BT_c02.tif | G1 | - | 500 | 3 | - |
| n002_G_Aph_4_AT_c01.tif | G2 | 40 | 300 | 1 | 500 |
| n002_G_Aph_4_AT_c02.tif | G2 | - | 500 | 3 | - |
| n002_G_Aph_4_BT_c01.tif | G2 | 40 | 300 | 1 | 500 |
| n002_G_Aph_4_BT_c02.tif | G2 | - | 500 | 3 | - |
| n003_G_Aph_5_AT_c01.tif | G2 | 40 | 300 | 1 | 500 |
| n003_G_Aph_5_AT_c02.tif | G2 | - | 500 | 3 | - |
| n003_G_Aph_5_BT_c01.tif | G2 | 40 | 300 | 1 | 500 |
| n003_G_Aph_5_BT_c02.tif | G2 | - | 500 | 3 | - |
| n004_G_Aph_5_AT_c01.tif | G2 | 40 | 300 | 1 | 500 |
| n004_G_Aph_5_AT_c02.tif | G2 | - | 500 | 3 | - |
| n004_G_Aph_5_BT_c01.tif | G2 | 40 | 300 | 1 | 500 |
| n004_G_Aph_5_BT_c02.tif | G2 | - | 500 | 3 | - |
| n005_G_Aph_5_AT_c01.tif | G1 | 40 | 300 | 1 | 500 |
| n005_G_Aph_5_AT_c02.tif | G1 | - | 500 | 3 | - |
| n005_G_Aph_5_BT_c01.tif | G1 | 40 | 300 | 1 | 500 |
| n005_G_Aph_5_BT_c02.tif | G1 | - | 500 | 3 | - |
| n006_G_Aph_6_AT_c01.tif | G2 | 40 | 300 | 1 | 500 |
| n006_G_Aph_6_AT_c02.tif | G2 | - | 500 | 3 | - |
| n006_G_Aph_6_BT_c01.tif | G2 | 40 | 300 | 1 | 500 |
| n006_G_Aph_6_BT_c02.tif | G2 | - | 500 | 3 | - |
| n007_G_Aph_10_AT_c01.tif | G1 | 40 | 300 | 1 | 500 |
| n007_G_Aph_10_AT_c02.tif | G1 | - | 500 | 3 | - |
| n007_G_Aph_10_BT_c01.tif | G1 | 40 | 300 | 1 | 500 |
| n007_G_Aph_10_BT_c02.tif | G1 | - | 500 | 3 | - |
| n008_G_Aph_10_AT_c01.tif | G1 | 40 | 300 | 1 | 500 |
| n008_G_Aph_10_AT_c02.tif | G1 | - | 500 | 3 | - |
| n008_G_Aph_10_BT_c01.tif | G1 | 40 | 300 | 1 | 500 |
| n008_G_Aph_10_BT_c02.tif | G1 | - | 500 | 3 | - |
| n009_G_Aph_10_AT_c01.tif | G1 | 40 | 300 | 1 | 500 |
| n009_G_Aph_10_AT_c02.tif | G1 | - | 500 | 3 | - |
| n009_G_Aph_10_BT_c01.tif | G1 | 40 | 300 | 1 | 500 |
| n009_G_Aph_10_BT_c02.tif | G1 | - | 500 | 3 | - |
| n0012_G_Aph_11_AT_c01.tif | G2 | 40 | 300 | 1 | 500 |
| n0012_G_Aph_11_AT_c02.tif | G2 | - | 500 | 3 | - |
| n0012_G_Aph_11_BT_c01.tif | G2 | 40 | 300 | 1 | 500 |
| n0012_G_Aph_11_BT_c02.tif | G2 | - | 500 | 3 | - |
| n0013_G_Aph_11_AT_c01.tif | G2 | 40 | 300 | 1 | 500 |
| n0013_G_Aph_11_AT_c02.tif | G2 | - | 500 | 3 | - |
| n0013_G_Aph_11_BT_c01.tif | G2 | 40 | 300 | 1 | 500 |
| n0013_G_Aph_11_BT_c02.tif | G2 | - | 500 | 3 | - |
| n0015_G_Aph_15_AT_c01.tif | G1 | 40 | 300 | 1 | 500 |
| n0015_G_Aph_15_AT_c02.tif | G1 | - | 500 | 3 | - |
| n0015_G_Aph_15_BT_c01.tif | G1 | 40 | 300 | 1 | 500 |
| n0015_G_Aph_15_BT_c02.tif | G1 | - | 500 | 3 | - |
| n0017_G_Aph_16_AT_c01.tif | G1 | 40 | 300 | 1 | 500 |
| n0017_G_Aph_16_AT_c02.tif | G1 | - | 500 | 3 | - |
| n0017_G_Aph_16_BT_c01.tif | G1 | 40 | 300 | 1 | 500 |
| n0017_G_Aph_16_BT_c02.tif | G1 | - | 500 | 3 | - |
| n0018_G_Aph_18_AT_c01.tif | G1 | 40 | 300 | 1 | 500 |
| n0018_G_Aph_18_AT_c02.tif | G1 | - | 500 | 3 | - |
| n0018_G_Aph_18_BT_c01.tif | G1 | 40 | 300 | 1 | 500 |
| n0018_G_Aph_18_BT_c02.tif | G1 | - | 500 | 3 | - |
| n0020_G_Aph_21_AT_c01.tif | G2 | 40 | 300 | 1 | 500 |
| n0020_G_Aph_21_AT_c02.tif | G2 | - | 500 | 3 | - |
| n0020_G_Aph_21_BT_c01.tif | G2 | 40 | 300 | 1 | 500 |
| n0020_G_Aph_21_BT_c02.tif | G2 | - | 500 | 3 | - |
| n001_S_Aph_1_AT_c01.tif | S | 40 | 300 | 1 | 500 |
| n001_S_Aph_1_AT_c02.tif | S | - | 500 | 3 | - |
| n001_S_Aph_1_BT_c01.tif | S | 40 | 300 | 1 | 500 |
| n001_S_Aph_1_BT_c02.tif | S | - | 500 | 3 | - |
| n002_S_Aph_1_AT_c01.tif | S | 40 | 300 | 1 | 500 |
| n002_S_Aph_1_AT_c02.tif | S | - | 500 | 3 | - |
| n002_S_Aph_1_BT_c01.tif | S | 40 | 300 | 1 | 500 |
| n002_S_Aph_1_BT_c02.tif | S | - | 500 | 3 | - |
| n004_S_Aph_1_AT_c01.tif | S | 40 | 300 | 1 | 500 |
| n004_S_Aph_1_AT_c02.tif | S | - | 500 | 3 | - |
| n004_S_Aph_1_BT_c01.tif | S | 40 | 300 | 1 | 500 |
| n004_S_Aph_1_BT_c02.tif | S | - | 500 | 3 | - |
| n006_S_Aph_2_AT_c01.tif | S | 40 | 300 | 1 | 500 |
| n006_S_Aph_2_AT_c02.tif | S | - | 500 | 3 | - |
| n006_S_Aph_2_BT_c01.tif | S | 40 | 300 | 1 | 500 |
| n006_S_Aph_2_BT_c02.tif | S | - | 500 | 3 | - |
| n008_S_Aph_4_AT_c01.tif | S | 40 | 300 | 1 | 500 |
| n008_S_Aph_4_AT_c02.tif | S | - | 500 | 3 | - |
| n008_S_Aph_4_BT_c01.tif | S | 40 | 300 | 1 | 500 |
| n008_S_Aph_4_BT_c02.tif | S | - | 500 | 3 | - |
| n009_S_Aph_5_AT_c01.tif | S | 40 | 300 | 1 | 500 |
| n009_S_Aph_5_AT_c02.tif | S | - | 500 | 3 | - |
| n009_S_Aph_5_BT_c01.tif | S | 40 | 300 | 1 | 500 |
| n009_S_Aph_5_BT_c02.tif | S | - | 500 | 3 | - |
| n0012_S_Aph_6_AT_c01.tif | S | 40 | 300 | 1 | 500 |
| n0012_S_Aph_6_AT_c02.tif | S | - | 500 | 3 | - |
| n0012_S_Aph_6_BT_c01.tif | S | 40 | 300 | 1 | 500 |
| n0012_S_Aph_6_BT_c02.tif | S | - | 500 | 3 | - |
| n0013_S_Aph_7_AT_c01.tif | S | 40 | 300 | 1 | 500 |
| n0013_S_Aph_7_AT_c02.tif | S | - | 500 | 3 | - |
| n0013_S_Aph_7_BT_c01.tif | S | 40 | 300 | 1 | 500 |
| n0013_S_Aph_7_BT_c02.tif | S | - | 500 | 3 | - |
| n0014_S_Aph_7_AT_c01.tif | S | 40 | 300 | 1 | 500 |
| n0014_S_Aph_7_AT_c02.tif | S | - | 500 | 3 | - |
| n0014_S_Aph_7_BT_c01.tif | S | 40 | 300 | 1 | 500 |
| n0014_S_Aph_7_BT_c02.tif | S | - | 500 | 3 | - |
| n0015_S_Aph_8_AT_c01.tif | S | 40 | 300 | 1 | 500 |
| n0015_S_Aph_8_AT_c02.tif | S | - | 500 | 3 | - |
| n0015_S_Aph_8_BT_c01.tif | S | 40 | 300 | 1 | 500 |
| n0015_S_Aph_8_BT_c02.tif | S | - | 500 | 3 | - |
| n0016_S_Aph_8_AT_c01.tif | S | 40 | 300 | 1 | 500 |
| n0016_S_Aph_8_AT_c02.tif | S | - | 500 | 3 | - |
| n0016_S_Aph_8_BT_c01.tif | S | - | 500 | 3 | 500 |
| n0016_S_Aph_8_BT_c02.tif | S | 40 | 300 | 1 | - |
| n0017_S_Aph_8_AT_c01.tif | S | - | 500 | 3 | 500 |
| n0017_S_Aph_8_AT_c02.tif | S | 40 | 300 | 1 | - |
| n0017_S_Aph_8_BT_c01.tif | S | - | 500 | 3 | 500 |
| n0017_S_Aph_8_BT_c02.tif | S | 40 | 300 | 1 | - |
| n0018_S_Aph_8_AT_c01.tif | S | - | 500 | 3 | 500 |
| n0018_S_Aph_8_AT_c02.tif | S | 40 | 300 | 1 | - |
| n0018_S_Aph_8_BT_c01.tif | S | - | 500 | 3 | 500 |
| n0018_S_Aph_8_BT_c02.tif | S | 40 | 300 | 1 | - |
| n0019_S_Aph_9_AT_c01.tif | S | - | 500 | 3 | 500 |
| n0019_S_Aph_9_AT_c02.tif | S | 40 | 300 | 1 | - |
| n0019_S_Aph_9_BT_c01.tif | S | - | 500 | 3 | 500 |
| n0019_S_Aph_9_BT_c02.tif | S | - | 500 | 3 | - |
| n0020_S_Aph_9_AT_c01.tif | S | 40 | 300 | 1 | 500 |
| n0020_S_Aph_9_AT_c02.tif | S | - | 500 | 3 | - |
| n0020_S_Aph_9_BT_c01.tif | S | 40 | 300 | 1 | 500 |
| n0020_S_Aph_9_BT_c02.tif | S | - | 500 | 3 | - |
| n0021_S_Aph_1_0_AT_c01.tif | S | 40 | 300 | 1 | 500 |
| n0021_S_Aph_1_0_AT_c02.tif | S | - | 500 | 3 | - |
| n0021_S_Aph_1_0_BT_c01.tif | S | 40 | 300 | 1 | 500 |
| n0021_S_Aph_1_0_BT_c02.tif | S | - | 500 | 3 | - |
| n0022_S_Aph_1_2_AT_c01.tif | S | 40 | 300 | 1 | 500 |
| n0022_S_Aph_1_2_AT_c02.tif | S | - | 500 | 3 | - |
| n0022_S_Aph_1_2_BT_c01.tif | S | 40 | 300 | 1 | 500 |
| n0022_S_Aph_1_2_BT_c02.tif | S | - | 500 | 3 | - |
| n0023_S_Aph_1_3_AT_c01.tif | S | 40 | 300 | 1 | 500 |
| n0023_S_Aph_1_3_AT_c02.tif | S | - | 500 | 3 | - |
| n0023_S_Aph_1_3_BT_c01.tif | S | 40 | 300 | 1 | 500 |
| n0023_S_Aph_1_3_BT_c02.tif | S | - | 500 | 3 | - |
| n0024_S_Aph_1_4_AT_c01.tif | S | 40 | 300 | 1 | 500 |
| n0024_S_Aph_1_4_AT_c02.tif | S | - | 500 | 3 | - |
| n0024_S_Aph_1_4_BT_c01.tif | S | 40 | 300 | 1 | 500 |
| n0024_S_Aph_1_4_BT_c02.tif | S | - | 500 | 3 | - |
| n0025_S_Aph_1_7_AT_c01.tif | S | 40 | 300 | 1 | 500 |
| n0025_S_Aph_1_7_AT_c02.tif | S | - | 500 | 3 | - |
| n0025_S_Aph_1_7_BT_c01.tif | S | 40 | 300 | 1 | 500 |
| n0025_S_Aph_1_7_BT_c02.tif | S | - | 500 | 3 | - |
| n0026_S_Aph_1_7_AT_c01.tif | S | 40 | 300 | 1 | 500 |
| n0026_S_Aph_1_7_AT_c02.tif | S | - | 500 | 3 | - |
| n0026_S_Aph_1_7_BT_c01.tif | S | 40 | 300 | 1 | 500 |
| n0026_S_Aph_1_7_BT_c02.tif | S | - | 500 | 3 | - |
| n0027_S_Aph_21_AT_c01.tif | S | 40 | 300 | 1 | 500 |
| n0027_S_Aph_21_AT_c02.tif | S | - | 500 | 3 | - |
| n0027_S_Aph_21_BT_c01.tif | S | 40 | 300 | 1 | 500 |
| n0027_S_Aph_21_BT_c02.tif | S | - | 500 | 3 | - |
| n0028_S_Aph_21_AT_c01.tif | S | 40 | 300 | 1 | 500 |
| n0028_S_Aph_21_AT_c02.tif | S | - | 500 | 3 | - |
| n0028_S_Aph_21_BT_c01.tif | S | 40 | 300 | 1 | 500 |
| n0028_S_Aph_21_BT_c02.tif | S | - | 500 | 3 | - |
| n0001_coloc_c02.tif | S | 40 | 300 | 2 | 500 |
| n0002_coloc_c02.tif | S | 40 | 300 | 2 | 500 |
| n0003_coloc_c02.tif | S | 40 | 300 | 2 | 500 |
| n0004_coloc_c02.tif | S | 40 | 300 | 2 | 500 |
| n0005_coloc_c02.tif | S | 40 | 300 | 2 | 500 |
| n0006_coloc_c02.tif | S | 40 | 300 | 2 | 500 |
| n0007_coloc_c02.tif | S | 40 | 300 | 2 | 500 |
| n0008_coloc_c02.tif | S | 40 | 300 | 2 | 500 |
| n0009_coloc_c02.tif | S | 40 | 300 | 2 | 500 |
| n0010_coloc_c02.tif | S | 40 | 300 | 2 | 500 |
| n0011_coloc_c02.tif | S | 40 | 300 | 2 | 500 |
| n0012_coloc_c02.tif | S | 40 | 300 | 2 | 500 |
| n0013_coloc_c02.tif | S | 40 | 300 | 2 | 500 |
| n0014_coloc_c02.tif | S | 40 | 300 | 2 | 500 |
| n0015_coloc_c02.tif | S | 40 | 300 | 2 | 500 |
| n0016_coloc_c02.tif | S | 40 | 300 | 2 | 500 |
| n0017_coloc_c02.tif | S | 40 | 300 | 2 | 500 |
| n0018_coloc_c02.tif | S | 40 | 300 | 2 | 500 |
| n0019_coloc_c02.tif | S | 40 | 300 | 2 | 500 |
| n0020_coloc_c02.tif | S | 40 | 300 | 2 | 500 |
| n0021_coloc_c02.tif | S | 40 | 300 | 2 | 500 |
| n0022_coloc_c02.tif | S | 40 | 300 | 2 | 500 |
| n0023_coloc_c02.tif | S | 40 | 300 | 2 | 500 |
| n0024_coloc_c02.tif | S | 40 | 300 | 2 | 500 |
| n0025_coloc_c02.tif | S | 40 | 300 | 2 | 500 |
| n0026_coloc_c02.tif | S | 40 | 300 | 2 | 500 |
| n0027_coloc_c02.tif | S | 40 | 300 | 2 | 500 |
| n0028_coloc_c02.tif | S | 40 | 300 | 2 | 500 |
| n0029_coloc_c02.tif | S | 40 | 300 | 2 | 500 |
| n0030_coloc_c02.tif | S | 40 | 300 | 2 | 500 |
| n0031_coloc_c02.tif | S | 40 | 300 | 2 | 500 |
| n0032_coloc_c02.tif | S | 40 | 300 | 2 | 500 |
| n0033_coloc_c02.tif | S | 40 | 300 | 2 | 500 |
| n001_G1_IMR90_PCNA_Cy3<br>dUTP_60x.tif | G1 | 40 | 300 | 2 | 500 |
| n002_G1_IMR90_PCNA_Cy3<br>dUTP_60x.tif | G1 | 40 | 300 | 2 | 500 |
| n003_G1_IMR90_PCNA_Cy3<br>dUTP_60x.tif | G1 | 40 | 300 | 2 | 500 |
| n004_G1_IMR90_PCNA_Cy3<br>dUTP_60x.tif | G1 | 40 | 300 | 2 | 500 |
| n005_G1_IMR90_PCNA_Cy3<br>dUTP_60x.tif | G1 | 40 | 300 | 2 | 500 |
| n006_G1_IMR90_PCNA_Cy3<br>dUTP_60x.tif | G1 | 40 | 300 | 2 | 500 |
| n007_G1_IMR90_PCNA_Cy3<br>dUTP_60x.tif | G1 | 40 | 300 | 2 | 500 |
| n008_G1_IMR90_PCNA_Cy3<br>dUTP_60x.tif | G1 | 40 | 300 | 2 | 500 |
| n009_G1_IMR90_PCNA_Cy3<br>dUTP_60x.tif | G1 | 40 | 300 | 2 | 500 |
| n010_G1_IMR90_PCNA_Cy3<br>dUTP_60x.tif | G1 | 40 | 300 | 2 | 500 |
| n011_G1_IMR90_PCNA_Cy3<br>dUTP_60x.tif | G1 | 40 | 300 | 2 | 500 |
| n012_G1_IMR90_PCNA_Cy3<br>dUTP_60x.tif | G1 | 40 | 300 | 2 | 500 |
| n013_G1_IMR90_PCNA_Cy3<br>dUTP_60x.tif | G1 | 40 | 300 | 2 | 500 |
| n014_G1_IMR90_PCNA_Cy3<br>dUTP_60x.tif | G1 | 40 | 300 | 2 | 500 |
| n001_G2_IMR90_PCNA_Cy3<br>dUTP_60x.tif | G2 | 40 | 300 | 2 | 500 |
| n002_G2_IMR90_PCNA_Cy3<br>dUTP_60x.tif | G2 | 40 | 300 | 2 | 500 |
| n003_G2_IMR90_PCNA_Cy3<br>dUTP_60x.tif | G2 | 40 | 300 | 2 | 500 |
| n004_G2_IMR90_PCNA_Cy3<br>dUTP_60x.tif | G2 | 40 | 300 | 2 | 500 |
| n005_G2_IMR90_PCNA_Cy3<br>dUTP_60x.tif | G2 | 40 | 300 | 2 | 500 |
| n006_G2_IMR90_PCNA_Cy3<br>dUTP_60x.tif | G2 | 40 | 300 | 2 | 500 |
| n007_G2_IMR90_PCNA_Cy3<br>dUTP_60x.tif | G2 | 40 | 300 | 2 | 500 |
| n008_G2_IMR90_PCNA_Cy3<br>dUTP_60x.tif | G2 | 40 | 300 | 2 | 500 |
| n009_G2_IMR90_PCNA_Cy3<br>dUTP_60x.tif | G2 | 40 | 300 | 2 | 500 |
| n010_G2_IMR90_PCNA_Cy3<br>dUTP_60x.tif | G2 | 40 | 300 | 2 | 500 |
| n011_G2_IMR90_PCNA_Cy3<br>dUTP_60x.tif | G2 | 40 | 300 | 2 | 500 |
| n012_G2_IMR90_PCNA_Cy3<br>dUTP_60x.tif | G2 | 40 | 300 | 2 | 500 |
| n013_G2_IMR90_PCNA_Cy3<br>dUTP_60x.tif | G2 | 40 | 300 | 2 | 500 |
| n014_G2_IMR90_PCNA_Cy3<br>dUTP_60x.tif | G2 | 40 | 300 | 2 | 500 |
| n015_G2_IMR90_PCNA_Cy3<br>dUTP_60x.tif | G2 | 40 | 300 | 2 | 500 |
| n001_S_IMR90_PCNA_Cy3d<br>UTP_60x.tif | S | 40 | 300 | 2 | 500 |
| n002_S_IMR90_PCNA_Cy3d<br>UTP_60x.tif | S | 40 | 300 | 2 | 500 |
| n003_S_IMR90_PCNA_Cy3d<br>UTP_60x.tif | S | 40 | 300 | 2 | 500 |
| n004_S_IMR90_PCNA_Cy3d<br>UTP_60x.tif | S | 40 | 300 | 2 | 500 |
| n005_S_IMR90_PCNA_Cy3d<br>UTP_60x.tif | S | 40 | 300 | 2 | 500 |
| n006_S_IMR90_PCNA_Cy3d<br>UTP_60x.tif | S | 40 | 300 | 2 | 500 |
| n007_S_IMR90_PCNA_Cy3d<br>UTP_60x.tif | S | 40 | 300 | 2 | 500 |
| n008_S_IMR90_PCNA_Cy3d<br>UTP_60x.tif | S | 40 | 300 | 2 | 500 |
| n009_S_IMR90_PCNA_Cy3d<br>UTP_60x.tif | S | 40 | 300 | 2 | 500 |
| n010_S_IMR90_PCNA_Cy3d<br>UTP_60x.tif | S | 40 | 300 | 2 | 500 |
| n011_S_IMR90_PCNA_Cy3d<br>UTP_60x.tif | S | 40 | 300 | 2 | 500 |
| n012_S_IMR90_PCNA_Cy3d<br>UTP_60x.tif | S | 40 | 300 | 2 | 500 |
| n0013_S_IMR90_PCNA_Cy3<br>dUTP_60x.tif | S | 40 | 300 | 2 | 500 |
| n0014_S_IMR90_PCNA_Cy3<br>dUTP_60x.tif | S | 40 | 300 | 2 | 500 |

**Supplementary table 9:** Software and macros

| Name | Version | Website | Company/University | Application |
| --- | --- | --- | --- | --- |
| Volocity | 6.3 | - | PerkinElmer, USA | Acquiring live cell time lapses |
| ImageJ | 1.53c | https://imagej.nih.gov/ij/ | Wayne Rasband,<br>National Institutes of Health, USA | Image processing and image analysis |
| RStudio | 1.1.447-1.2.5033 | https://rstudio.com/ | RStudio | Statistical analysis and plotting |
| Harmony | 3.5.1 | https://www.perkinelmer.com/product/harmony-4-8-office-hh17000001 | PerkinElmer, USA | High content microscopy imaging and analysis |
| KNIME Analytics Platform | 3.5.2 | https://www.knime.com/knime-analytics-platform | KNIME AG,<br>Switzerland | High content microscopy image processing and analysis |
| Adobe Illustrator CS6 | 16 | https://www.adobe.com/ | Adobe, USA | Graphical sketch and figures arrangement |

**Figure S1:**
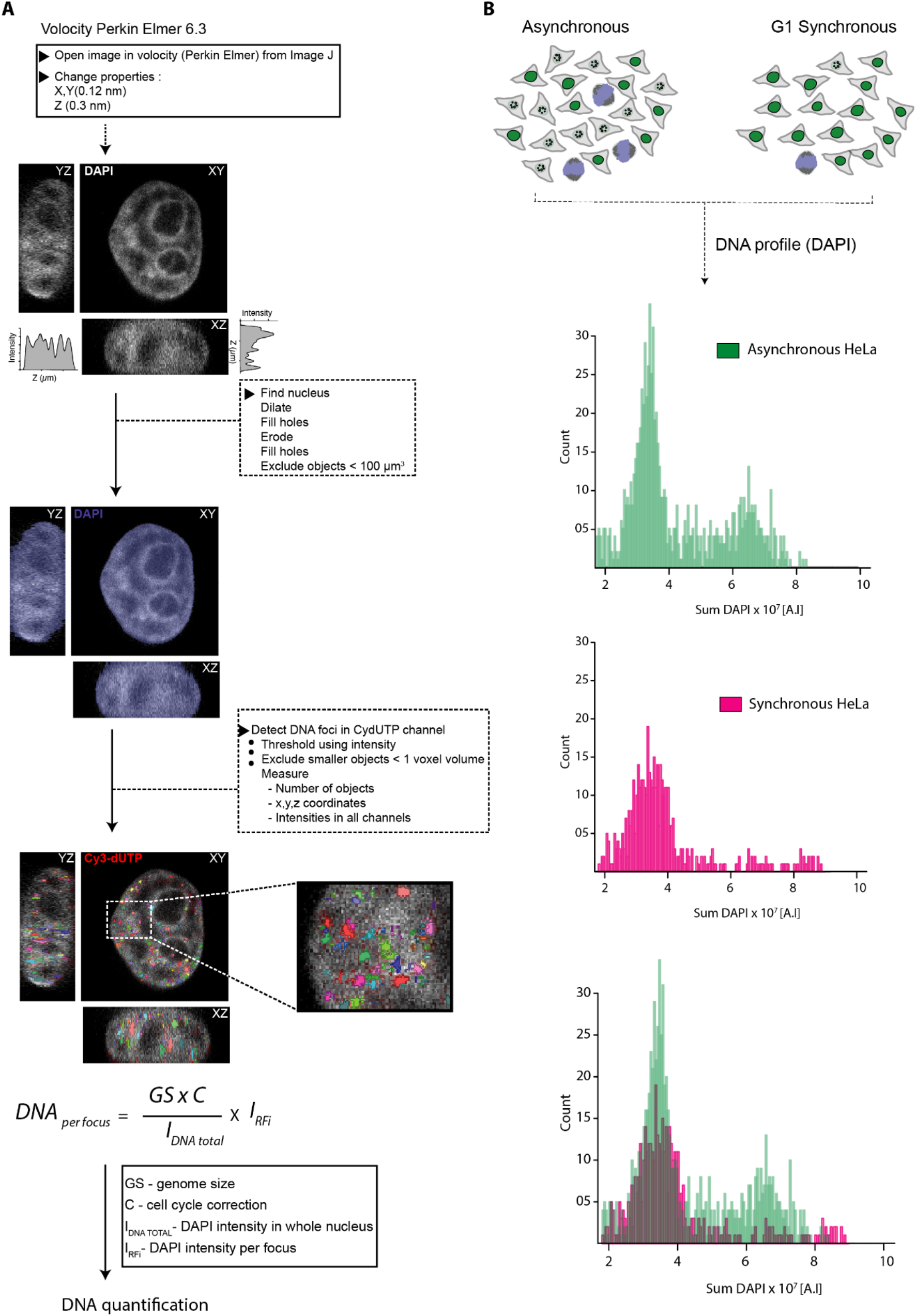
Pipeline and controls for DNA quantification of labeled replication foci using confocal data. **(A)** All image sequences were acquired using spinning disk microscopy (60x objective, 1.45 numerical aperture). For analysis, the (16 bit) images were imported from ImageJ into PerkinElmer Volocity 6.3 software. Using the DAPI channel, nuclear segmentation (object recognition) was performed to obtain a mask for the whole nuclear volume and determine the total DNA intensity (mask highlighted with blue). The labeled chromatin foci (Cy3-dUTP) were detected using the intensity and segmented in 3D (highlighted in color palette). The sum DAPI intensity within the segmented foci (I_RFi_) were also obtained. The ratio of DNA amount per focus and total DNA in the nucleus corrected for cell cycle stage gives the DNA amount per focus. For quantification see Figure 2. The intensity values of the total DAPI intensity within each segmented replication foci (I_RFi_) are plotted as a histogram. **(B)** HeLa Kyoto cells, either synchronously (magenta) (Methods) or asynchronously growing (green), were seeded on coverslips. The sum DAPI (DNA profile) of cells was plotted individual and in an overlap histogram (brown).

**Figure S2:**
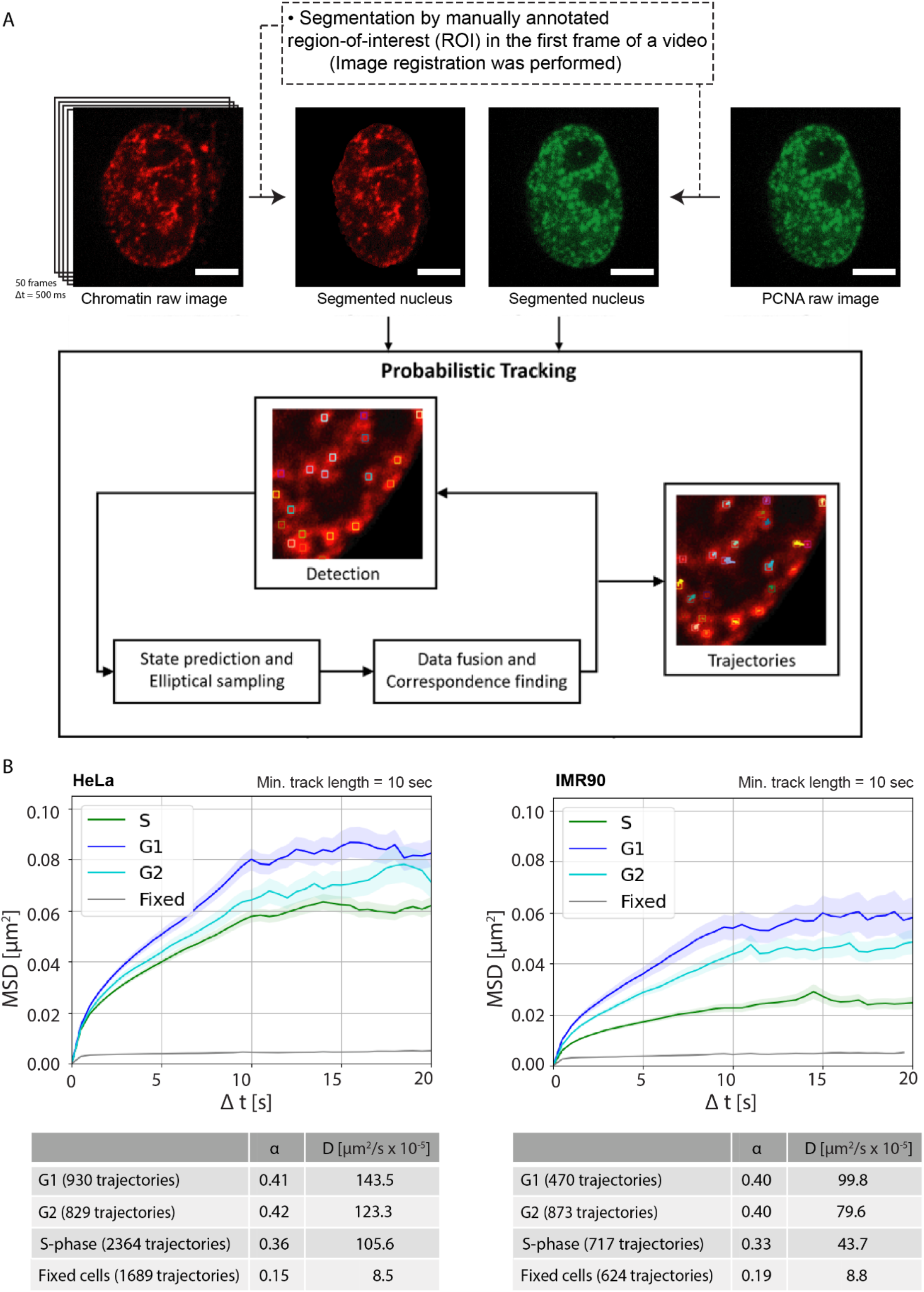
Workflow of chromatin motion and proximity analysis for confocal data. **(A)** Raw images of chromatin and PCNA were first segmented to remove the background noise outside the nucleus. In case of IMR90 cells, affine image registration was performed to address the stronger cell movement compared to HeLa cells (Methods). The probabilistic tracking method was then performed which includes detection of chromatin particles and tracking over the time to obtain trajectories. The tracks were then used to plot MSD curves or perform proximity analysis with PCNA. (**B)** Result of motion analysis of computed chromatin tracks for G1, G2, S-phase cell cycle stages (Figure 3) for HeLa and IMR90 cells. Mean Square Displacement (MSD, μm2) curves were plotted over time (s). Mean square displacement curves for G1/G2, S-phase, fixed cells with a minimum track length of 10 seconds and a total time of 20 seconds were plotted with error bars (SEM-standard error of the mean) representing the deviations between the MSD curves for an image sequence in transparent color around the curve. Scale bar: 10 µm.

**Figure S3:**
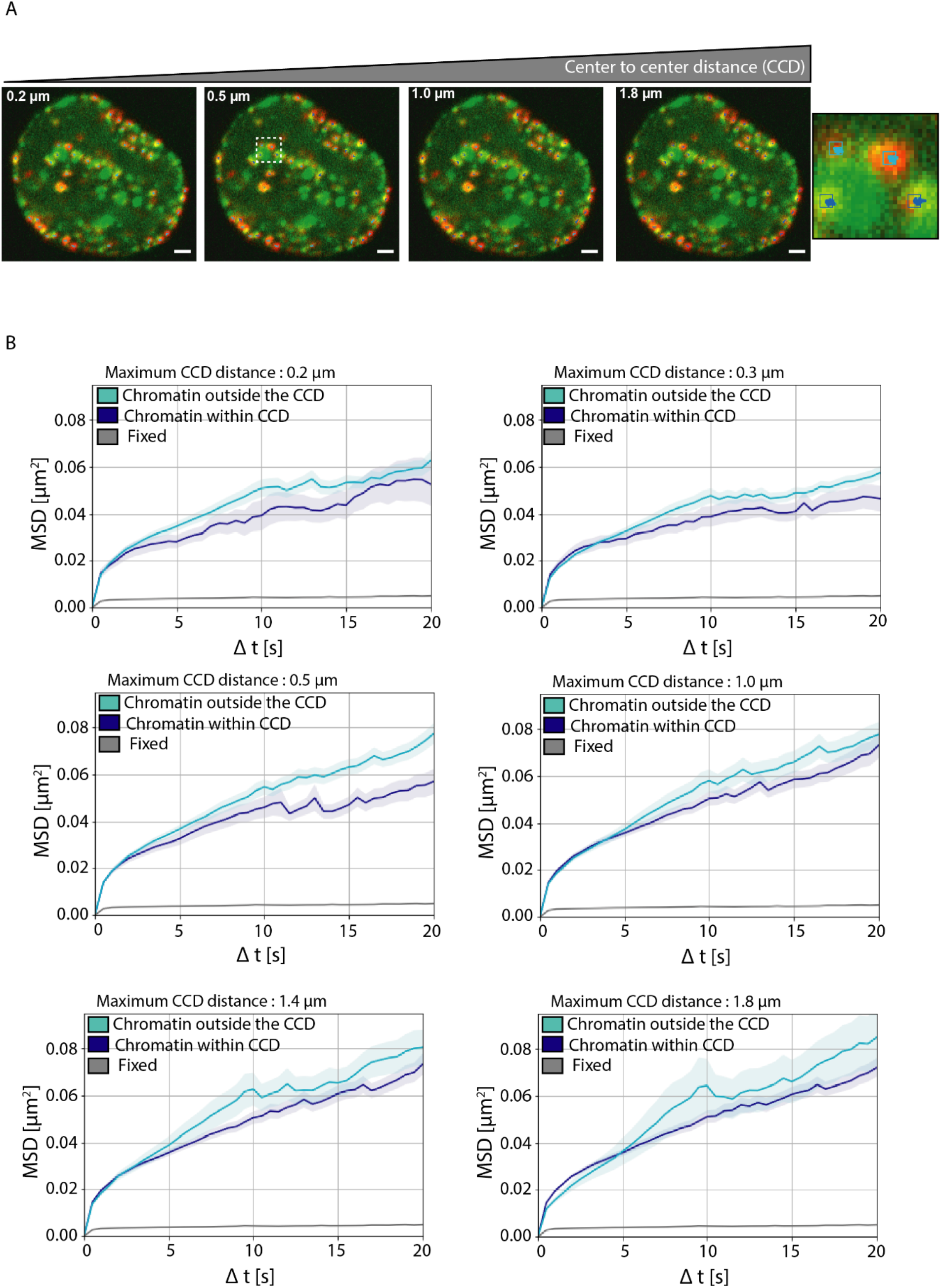
Mean square displacement analysis of labeled chromatin in the proximity of active replication sites at varying center to center distances. **(A)** Overlay images of GFP-PCNA (active replication sites, green) and labeled chromatin (red). Chromatin outside the center to center distance (CCD) is highlighted with cyan dots and chromatin within center to center distance (CCD) is highlighted with dark blue dots within the magnified inset (demarcated by a white box). **(B)** Mean square displacement (MSD) curves of chromatin outside the center to center distance (CCD) and chromatin within center to center distance (CCD) along with fixed cell control with increasing distance from 0.2 μm to 1.8 μm. This allows us to analyze the effect of replication factors on chromatin mobility at increasing distances.The MSD curves were plotted with error bars (standard deviation) represented in transparent color around the curve. Scale bar: 1 µm.

**Figure S4:**
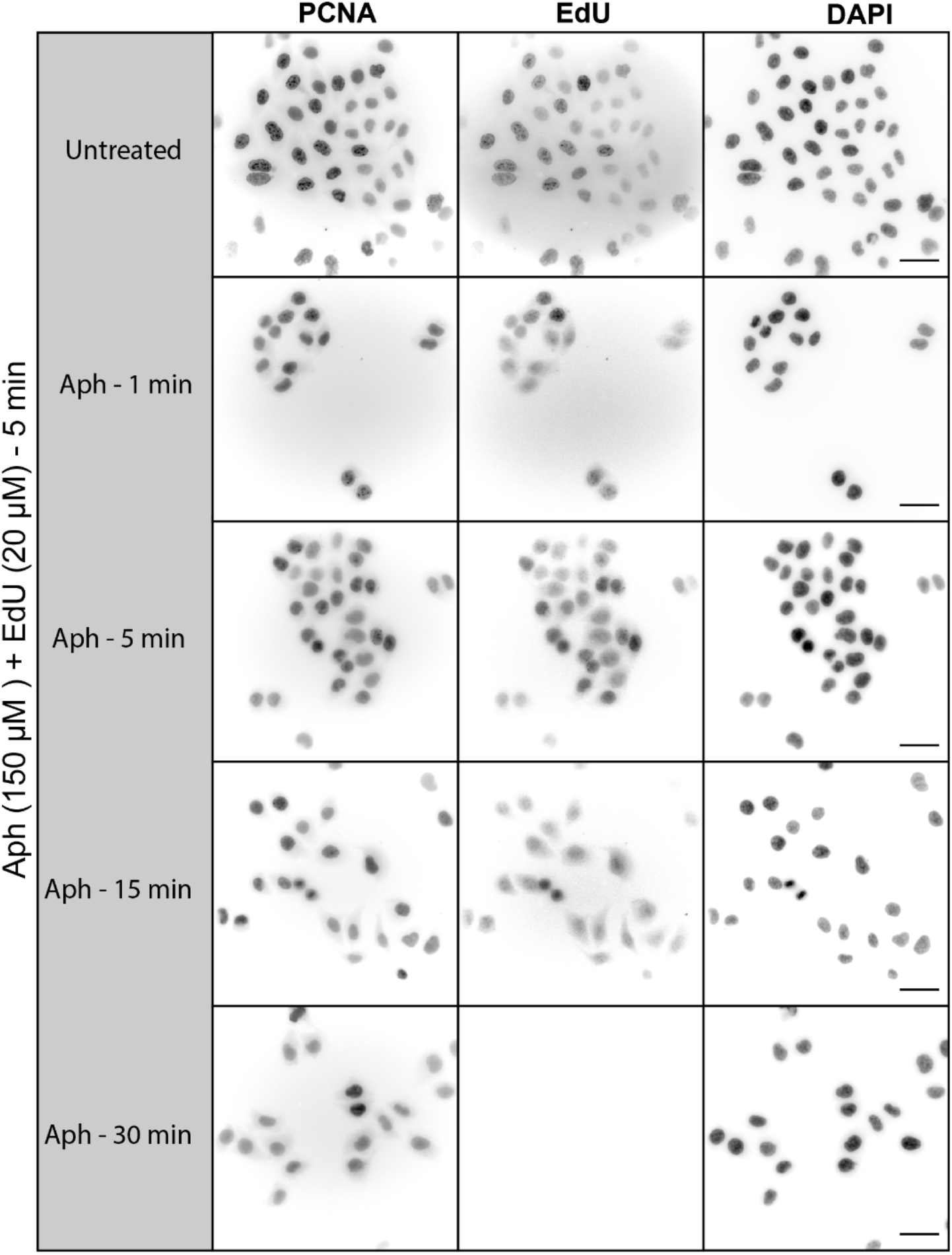
Time course analysis of inhibition of DNA synthesis upon aphidicolin treatment. **(A)** HeLa cells stably expressing GFP-PCNA were treated with aphidicolin (150 µm). Time course of treatments depicting the distribution of PCNA as well as the incorporation of the thymidine analog EdU is shown. EdU was used to detect DNA synthesis. For quantification see Figure 5A. Scale bar: 100 µm.

**Figure S5:**
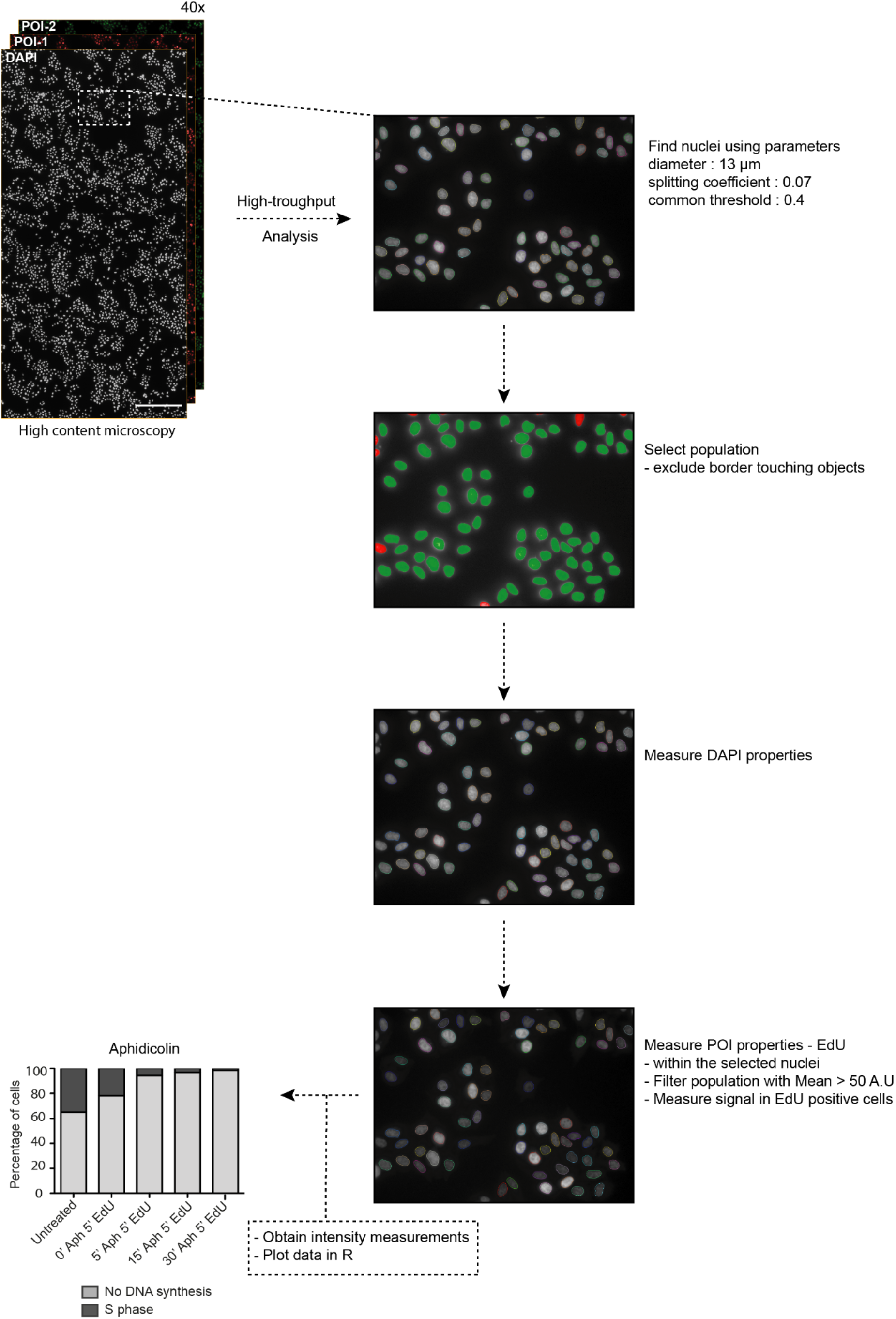
Schematic overview of the high throughput analysis pipeline of DNA synthesis inhibition by aphidicolin. Images were obtained using the 40x objective (numerical aperture 0.95) in the DAPI and EdU channels. Once the images were acquired they were analyzed using the Harmony software from PerkinElmer. Briefly, the DAPI channel was used to segment nuclei using properties including diameter, splitting coefficient and threshold. Once the nuclei were segmented, the population cells were then selected for analysis by removing the objects either touching the border or fused. In the population of cells selected, the properties including mean, median, standard deviation and sum intensities were measured in the DAPI and EdU channels. The data measurements were then imported to R and visualized for the EdU positive population. Scale bar: 500 μm.

**Figure S6:**
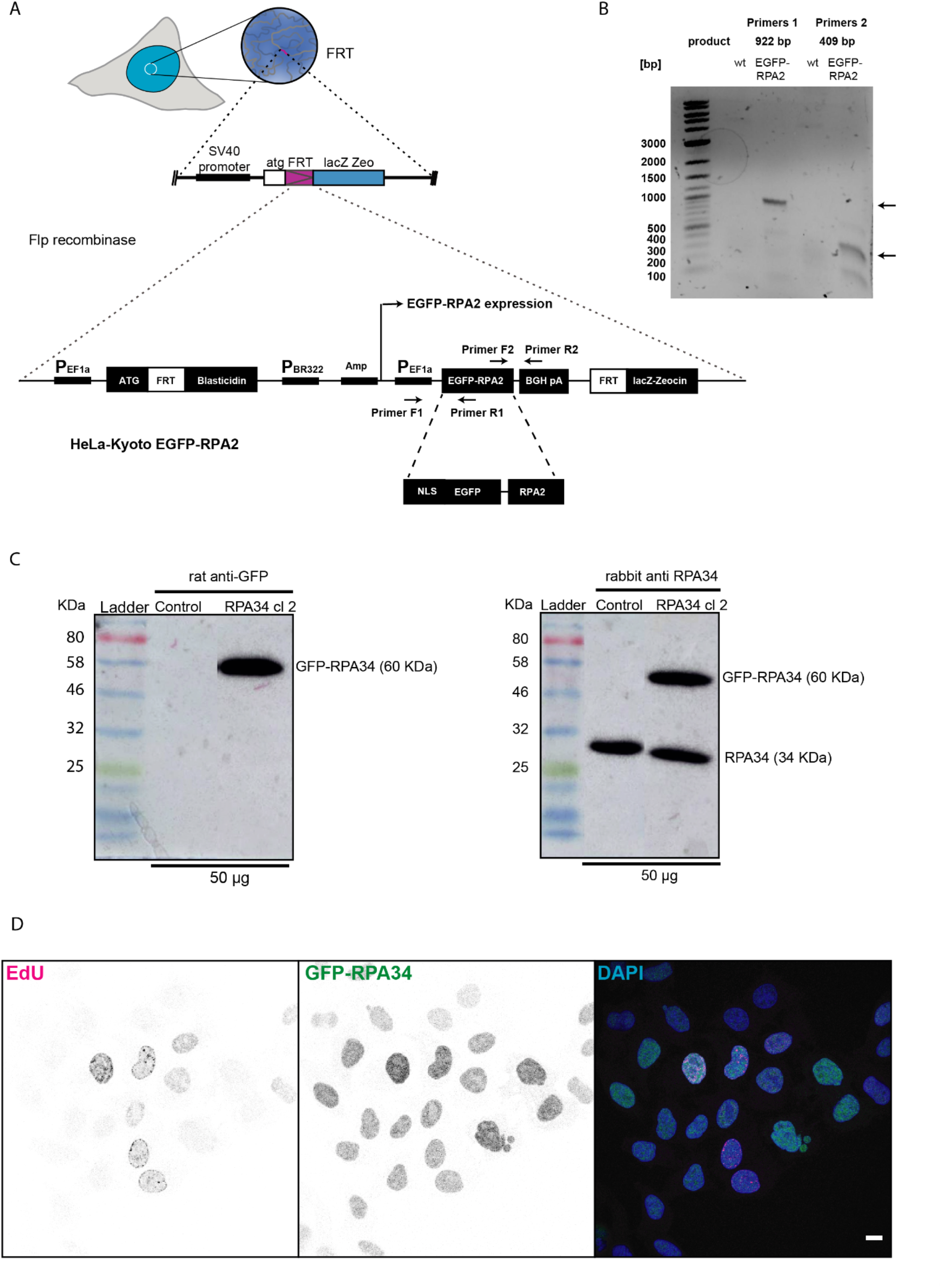
HeLa GFP-RPA cell line characterization after generation using the Flp-recombinase system. **(A)** Schematic representation of the insertion of fluorescently tagged RPA34 (aka RPA2) encoding plasmid DNA using the chromosomally inserted FRT site in the previously generated HeLa Kyoto FRTLacZ (Methods). Positive clones were selected using 1 μg/ml blasticidin in media. **(B)** PCR was performed on genomic DNA isolated from wild type and GFP-RPA34 clones using different primer pairs as indicated. **(C)** Cells from the positive clones were collected to perform a western blot using the anti-GFP antibody to test the level of GFP-RPA34 and an anti-RPA34 antibody was used to detect and compare both endogenous and fusion protein levels in control versus the positive clone. **(D)** The cells were seeded on glass coverslips, fixed and imaged to validate the protein localization relative to active DNA replication sites labeled EdU as well as DNA labeled with DAPI. Scale bar: 5 µm.

**Figure S7:**
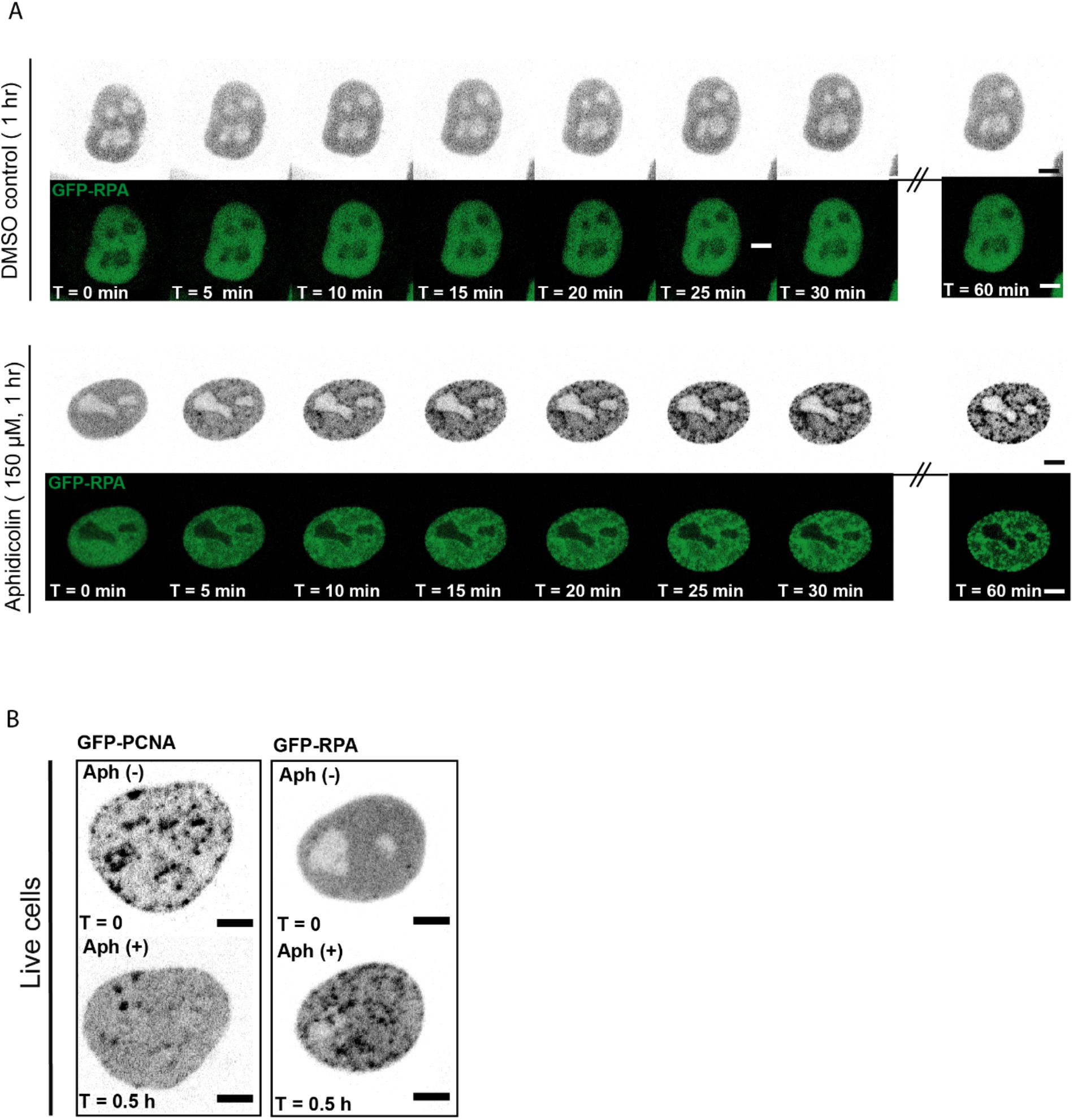
Live-cell time-lapse microscopy of HeLa GFP-RPA34 cells to determine RPA accumulation at replication sites upon aphidicolin treatment. **(A)** Confocal time-lapse microscopy of HeLa cells stably expressing GFP-RPA34 treated with DMSO (control), and aphidicolin. Movies were acquired at a frame rate of 5 minutes and at every time a Z stack was acquired. A single Z-plane at different timepoints was shown as a representative image for all three conditions. The time-lapse images were imported to PerkinElmer Volocity 6.3 for analysis and the accumulation of GFP-RPA34 at replication foci was measured over time. **(B)** Representative images of live HeLa cells stably expressing GFP-PCNA or GFP-RPA34 are shown in gray scale before and after aphidicolin treatment. Scale bar: 5 µm.

**Figure S8:**
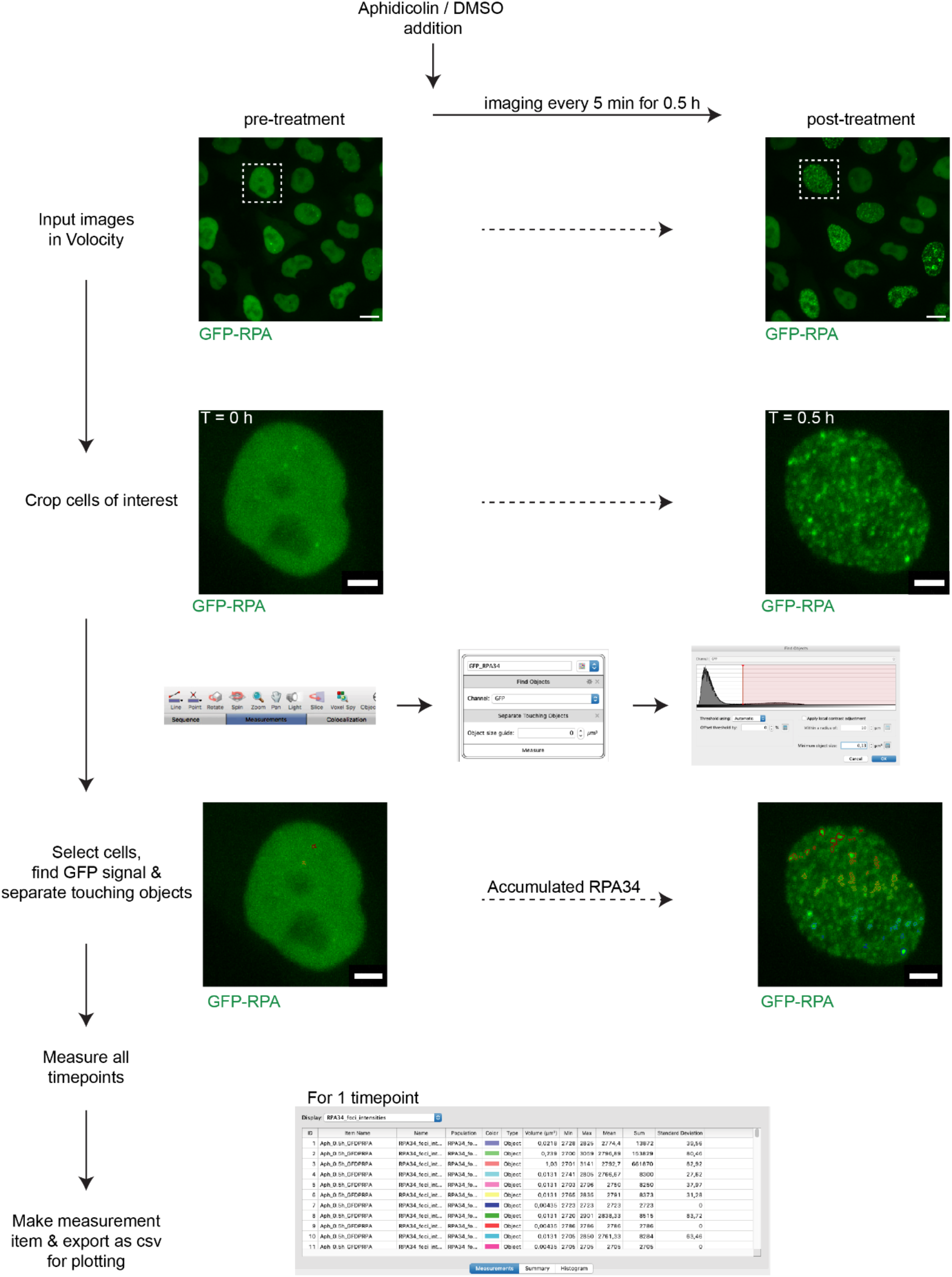
Pipeline for the analysis of RPA enrichment at replication sites in living cells. The enrichment or accumulation analysis was performed using PerkinElmer Volocity 6.3 software. Briefly cells of interest (white inserts) were cropped and Z-stacks along with different timepoints were imported together. The GFP-RPA34 intensities at different timepoints for aphidicolin (every 5 min / 0.5 h) were measured and the coefficient of variation c_V_ was calculated (Methods, Accumulation analysis). The intensity values were exported as CSV files for further analysis. The plots were generated in R studio. Scale bar: 5 μm. Cropped cells scale bar

**Figure S9:**
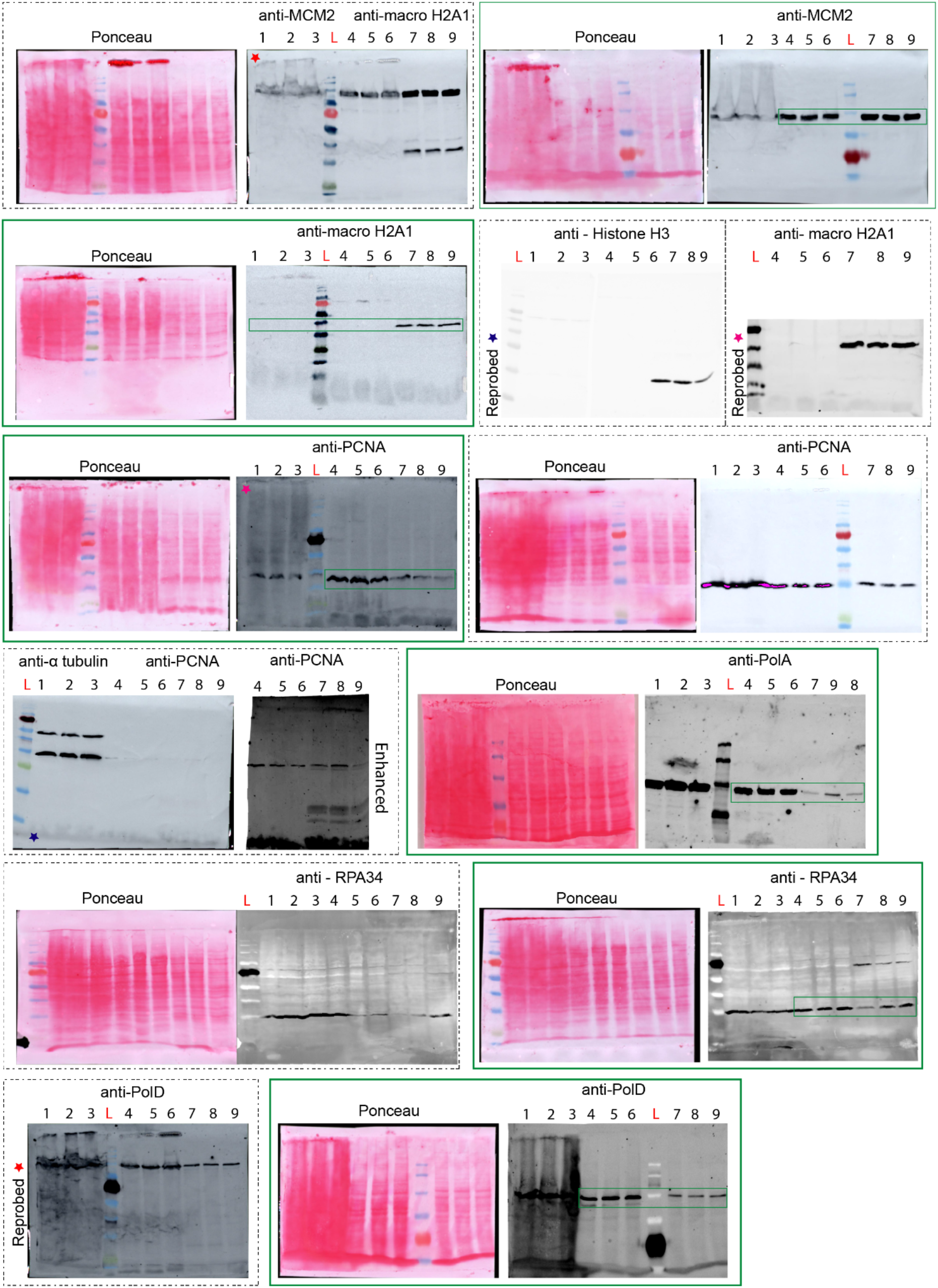
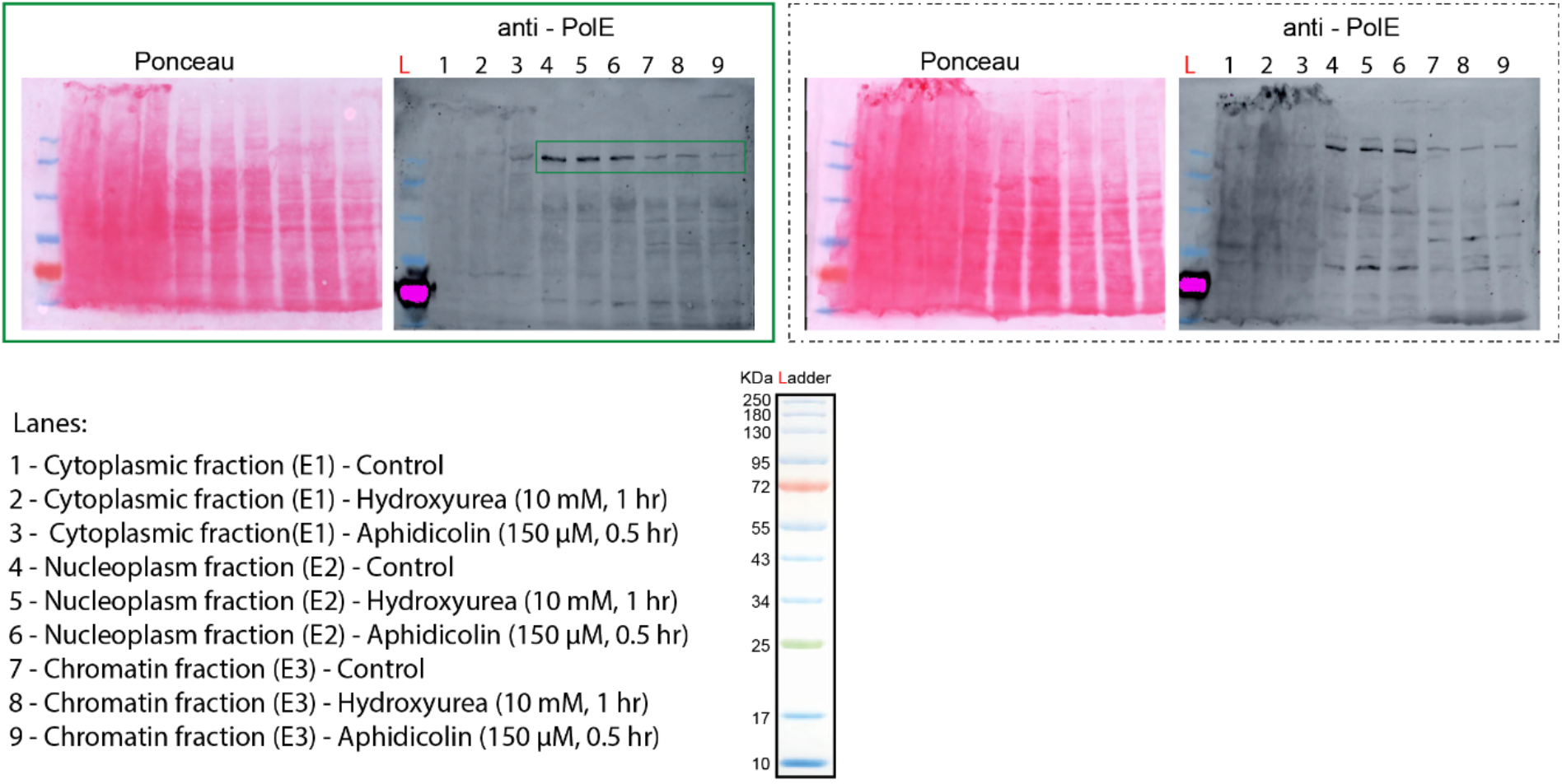
Full length western blots probed for different replication factors with cytoplasm, nucleoplasm and chromatin fractions. The blots that are reprobed with a different antibody are highlighted with color stars. The blots highlighted with green inserts are the full length and single contrast blots that are cropped for easier visualization as the loading order of samples and ladder is different and are summarized in Figure 5C.

**Figure S10:**
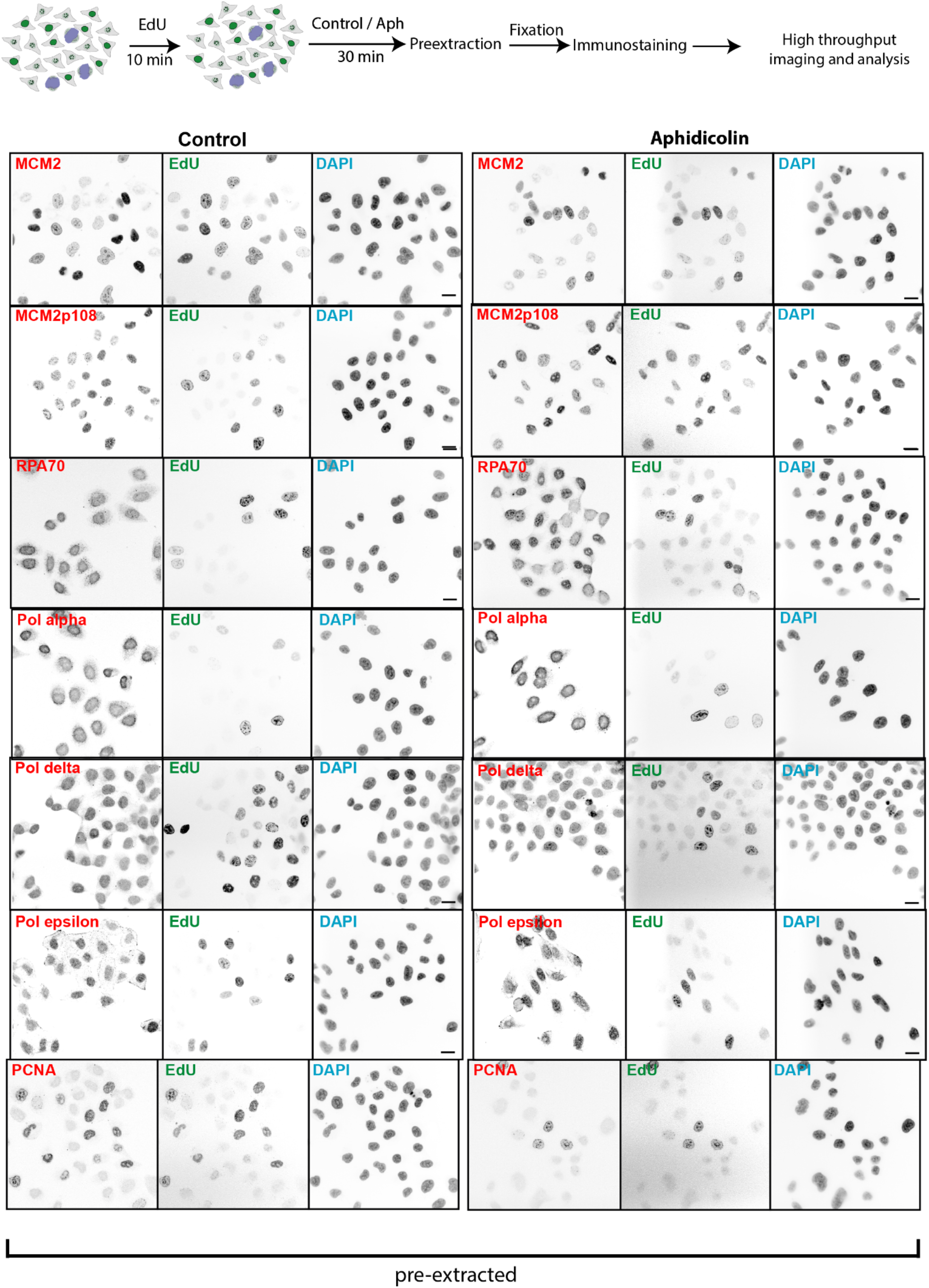
Analysis of replisome component enrichment at replication sites upon replication stress. Spatial localization and accumulation analysis of DNA replication factors such as DNA helicase subunit MCM2 and active MCM2 phosphorylated at S108, polymerase clamp protein (PCNA), single stranded binding protein (RPA) and DNA polymerases (α, δ, ε) catalytic subunits using immunostainings. The cells in S-phase were identified using an EdU pulse before the treatment. Representative low magnification images of protein localization in control and aphidicolin treated HeLa cells. For quantification see Figure 5D. Scale bar: 10 µm.

**Figure S11:**
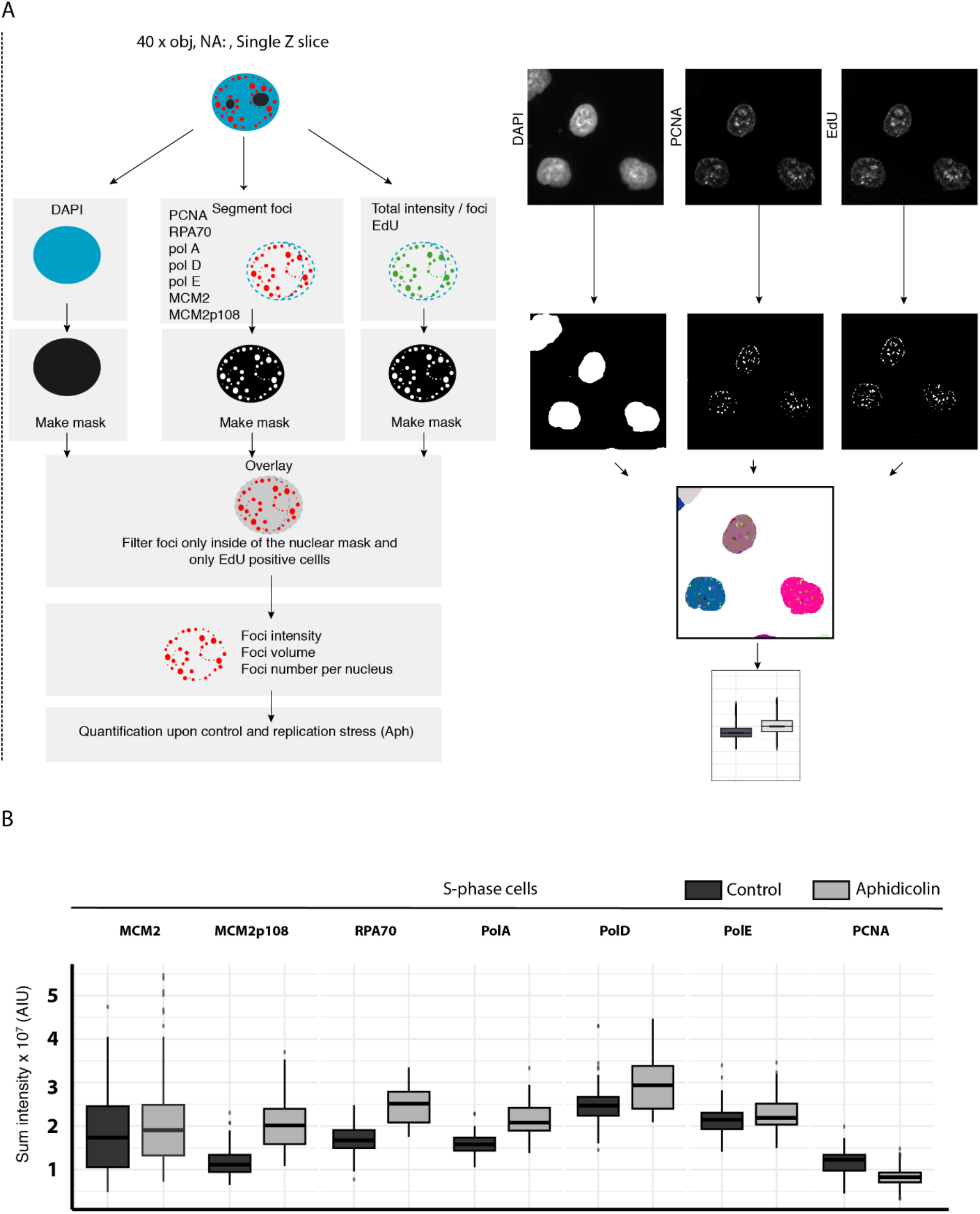
Pipeline for the analysis of replisome components on chromatin upon stress. **(A)** The accumulation analysis was performed. The channels were separated, the DAPI channel was used for creation of nuclear mask, the EdU and replication proteins channels were used for replication foci segmentation. The nuclear mask was overlaid with the replication foci mask, so that only foci inside of the nuclear area were selected. The replication foci parameters were measured as follows: focus intensity parameters, focus area, foci number per nucleus (Methods, High throughput image analysis of replisome component). The replication foci values per nucleus together with the DAPI nuclear intensity were exported for further analysis. The plots were generated in R studio. **(B)** Box plots from Figure 5D are shown here at the same Y axis scale.

**Figure S12:**
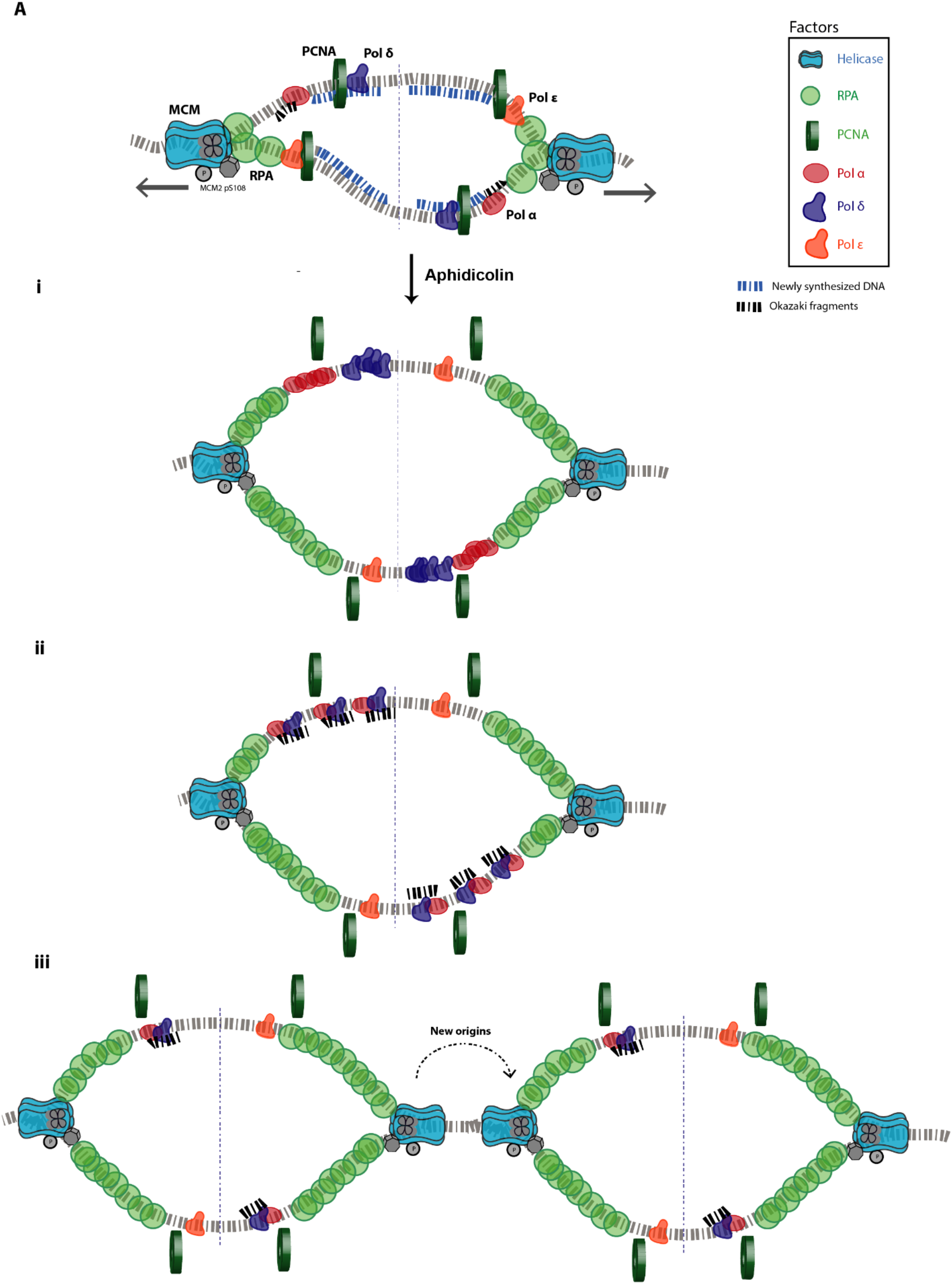
Model describing the different scenarios of replisome response to stress.

## Movie captions

Movie 1: Time lapse microscopy of HeLa K cells in G1 phase expressing fluorescent PCNA (green) and labeled chromatin (red). Scale bar: 5 µm.

Movie 2: Time lapse microscopy of HeLa K cells in G2 phase expressing fluorescent PCNA (green) and labeled chromatin (red). Scale bar: 5 µm.

Movie 3: Time lapse microscopy of HeLa K cells in S phase expressing fluorescent PCNA (green) and labeled chromatin (red). Scale bar: 5 µm.

Movie 4: Time lapse microscopy of IMR90 cells in G1 phase expressing fluorescent PCNA (green) and labeled chromatin (red). Scale bar: 5 µm.

Movie 5: Time lapse microscopy of IMR90 cells in S phase expressing fluorescent PCNA (green) and labeled chromatin (red). Scale bar: 5 µm.

Movie 6: Time lapse microscopy of HeLa K cells pre and post aphidicolin treatment in G1/G2 phase expressing fluorescent PCNA and RPA and labeled chromatin (red). Scale bar: 5 µm.

Movie 7: Time lapse microscopy of HeLa K cells pre and post aphidicolin treatment in S phase expressing fluorescent PCNA and RPA and labeled chromatin (red). Scale bar: 5 µm.

## References

1. Babokhov M, Hibino K, Itoh Y, Maeshima K. 2020. Local chromatin motion and transcription. J Mol Biol 432:694–700. doi:10.1016/j.jmb.2019.10.018

2. Bacher CP, Reichenzeller M, Athale C, Herrmann H, Eils R. 2004. 4-D single particle tracking of synthetic and proteinaceous microspheres reveals preferential movement of nuclear particles along chromatin - poor tracks. BMC Cell Biol 5:45. doi:10.1186/1471-2121-5-45

3. Baddeley D, Chagin VO, Schermelleh L, Martin S, Pombo A, Carlton PM, Gahl A, Domaing P, Birk U, Leonhardt H, Cremer C, Cardoso MC. 2010. Measurement of replication structures at the nanometer scale using super-resolution light microscopy. Nucleic Acids Res 38:e8. doi:10.1093/nar/gkp901

4. Bambara RA, Jessee CB. 1991. Properties of DNA polymerases delta and epsilon, and their roles in eukaryotic DNA replication. Biochim Biophys Acta 1088:11–24. doi:10.1016/0167-4781(91)90147-e

5. Baranovskiy AG, Babayeva ND, Suwa Y, Gu J, Pavlov YI, Tahirov TH. 2014. Structural basis for inhibition of DNA replication by aphidicolin. Nucleic Acids Res 42:14013–14021. doi:10.1093/nar/gku1209

6. Bentley JL. 1975. Multidimensional binary search trees used for associative searching. Commun ACM 18:509–517. doi:10.1145/361002.361007

7. Bornfleth H, Edelmann P, Zink D, Cremer T, Cremer C. 1999. Quantitative motion analysis of subchromosomal foci in living cells using four-dimensional microscopy. Biophys J 77:2871–2886. doi:10.1016/S0006-3495(99)77119-5

8. Bronshtein I, Kanter I, Kepten E, Lindner M, Berezin S, Shav-Tal Y, Garini Y. 2016. Exploring chromatin organization mechanisms through its dynamic properties. Nucleus 7:27–33. doi:10.1080/19491034.2016.1139272

9. Bronshtein I, Kepten E, Kanter I, Berezin S, Lindner M, Redwood AB, Mai S, Gonzalo S, Foisner R, Shav-Tal Y, Garini Y. 2015. Loss of lamin A function increases chromatin dynamics in the nuclear interior. Nat Commun 6:8044. doi:10.1038/ncomms9044

10. Byrnes JJ. 1984. Structural and functional properties of DNA polymerase delta from rabbit bone marrow. Mol Cell Biochem 62:13–24. doi:10.1007/BF00230073

11. Casas-Delucchi CS, Cardoso MC. 2011. Epigenetic control of DNA replication dynamics in mammals. Nucleus 2:370–382. doi:10.4161/nucl.2.5.17861

12. Chagin VO, Casas-Delucchi CS, Reinhart M, Schermelleh L, Markaki Y, Maiser A, Bolius JJ, Bensimon A, Fillies M, Domaing P, Rozanov YM, Leonhardt H, Cardoso MC. 2016. 4D Visualization of replication foci in mammalian cells corresponding to individual replicons. Nat Commun 7:11231. doi:10.1038/ncomms11231

13. Chagin VO, Reinhart B, Becker A, Mortusewicz O, Jost KL, Rapp A, Leonhardt H, Cardoso MC. 2019. Processive DNA synthesis is associated with localized decompaction of constitutive heterochromatin at the sites of DNA replication and repair. Nucleus 10:231–253. doi:10.1080/19491034.2019.1688932

14. Chagin VO, Reinhart M, Cardoso MC. 2015. High-resolution analysis of Mammalian DNA replication units. Methods Mol Biol 1300:43–65. doi:10.1007/978-1-4939-2596-4_3

15. Cheng CH, Kuchta RD. 1993. DNA polymerase epsilon: aphidicolin inhibition and the relationship between polymerase and exonuclease activity. Biochemistry 32:8568–8574. doi:10.1021/bi00084a025

16. Chubb JR, Boyle S, Perry P, Bickmore WA. 2002. Chromatin motion is constrained by association with nuclear compartments in human cells. Curr Biol 12:439–445. doi:10.1016/s0960-9822(02)00695-4

17. Conte M, Fiorillo L, Bianco S, Chiariello AM, Esposito A, Nicodemi M. 2020. Polymer physics indicates chromatin folding variability across single-cells results from state degeneracy in phase separation. Nat Commun 11:3289. doi:10.1038/s41467-020-17141-4

18. Cremer M, Brandstetter K, Maiser A, Rao SSP, Schmid VJ, Guirao-Ortiz M, Mitra N, Mamberti S, Klein KN, Gilbert DM, Leonhardt H, Cardoso MC, Aiden EL, Harz H, Cremer T. 2020. Cohesin depleted cells rebuild functional nuclear compartments after endomitosis. Nat Commun 11:6146. doi:10.1038/s41467-020-19876-6

19. Easwaran HP, Leonhardt H, Cardoso MC. 2005. Cell cycle markers for live cell analyses. Cell Cycle 4:453–455. doi:10.4161/cc.4.3.1525

20. Eaton JA, Zidovska A. 2020. Structural and dynamical signatures of local DNA damage in live cells. Biophys J 118:2168–2180. doi:10.1016/j.bpj.2019.10.042

21. Erfle H, Neumann B, Liebel U, Rogers P, Held M, Walter T, Ellenberg J, Pepperkok R. 2007. Reverse transfection on cell arrays for high content screening microscopy. Nat Protoc 2:392–399. doi:10.1038/nprot.2006.483

22. Esposito A, Annunziatella C, Bianco S, Chiariello AM, Fiorillo L, Nicodemi M. 2019. Models of polymer physics for the architecture of the cell nucleus. Wiley Interdiscip Rev Syst Biol Med 11:e1444. doi:10.1002/wsbm.1444

23. Esposito A, Bianco S, Fiorillo L, Conte M, Abraham A, Musella F, Nicodemi M, Prisco A, Chiariello AM. 2021. Polymer models are a versatile tool to study chromatin 3D organization. Biochem Soc Trans 49:1675–1684. doi:10.1042/BST20201004

24. Forsburg SL. 2004. Eukaryotic MCM proteins: beyond replication initiation. Microbiol Mol Biol Rev 68:109–131. doi:10.1128/MMBR.68.1.109-131.2004

25. Gasser SM. 2002. Visualizing chromatin dynamics in interphase nuclei. Science 296:1412–1416. doi:10.1126/science.1067703

26. Germier T, Kocanova S, Walther N, Bancaud A, Shaban HA, Sellou H, Politi AZ, Ellenberg J, Gallardo F, Bystricky K. 2017. Real-Time Imaging of a Single Gene Reveals Transcription-Initiated Local Confinement. Biophys J 113:1383–1394. doi:10.1016/j.bpj.2017.08.014

27. Ge XQ, Jackson DA, Blow JJ. 2007. Dormant origins licensed by excess Mcm2-7 are required for human cells to survive replicative stress. Genes Dev 21:3331–3341. doi:10.1101/gad.457807

28. Ghosh RN, Webb WW. 1994. Automated detection and tracking of individual and clustered cell surface low density lipoprotein receptor molecules. Biophys J 66:1301–1318. doi:10.1016/S0006-3495(94)80939-7

29. Gillotin S. 2018. Isolation of Chromatin-bound Proteins from Subcellular Fractions for Biochemical Analysis. Bio Protoc 8:e3035. doi:10.21769/BioProtoc.3035

30. Giorgetti L, Heard E. 2016. Closing the loop: 3C versus DNA FISH. Genome Biol 17:215. doi:10.1186/s13059-016-1081-2

31. Godinez WJ, Rohr K. 2015. Tracking multiple particles in fluorescence time-lapse microscopy images via probabilistic data association. IEEE Trans Med Imaging 34:415–432. doi:10.1109/TMI.2014.2359541

32. Görisch SM, Sporbert A, Stear JH, Grunewald I, Nowak D, Warbrick E, Leonhardt H, Cardoso MC. 2008. Uncoupling the replication machinery: replication fork progression in the absence of processive DNA synthesis. Cell Cycle 7:1983–1990. doi:10.4161/cc.7.13.6094

33. Gu B, Swigut T, Spencley A, Bauer MR, Chung M, Meyer T, Wysocka J. 2018. Transcription-coupled changes in nuclear mobility of mammalian cis-regulatory elements. Science 359:1050–1055. doi:10.1126/science.aao3136

34. Hauer MH, Gasser SM. 2017. Chromatin and nucleosome dynamics in DNA damage and repair. Genes Dev 31:2204–2221. doi:10.1101/gad.307702.117

35. Hauer MH, Seeber A, Singh V, Thierry R, Sack R, Amitai A, Kryzhanovska M, Eglinger J, Holcman D, Owen-Hughes T, Gasser SM. 2017. Histone degradation in response to DNA damage enhances chromatin dynamics and recombination rates. Nat Struct Mol Biol 24:99–107. doi:10.1038/nsmb.3347

36. Heun P, Laroche T, Shimada K, Furrer P, Gasser SM. 2001. Chromosome dynamics in the yeast interphase nucleus. Science 294:2181–2186. doi:10.1126/science.1065366

37. Hnisz D, Shrinivas K, Young RA, Chakraborty AK, Sharp PA. 2017. A phase separation model for transcriptional control. Cell 169:13–23. doi:10.1016/j.cell.2017.02.007

38. Ibarra A, Schwob E, Méndez J. 2008. Excess MCM proteins protect human cells from replicative stress by licensing backup origins of replication. Proc Natl Acad Sci USA 105:8956–8961. doi:10.1073/pnas.0803978105

39. Jackson DA, Pombo A. 1998. Replicon clusters are stable units of chromosome structure: evidence that nuclear organization contributes to the efficient activation and propagation of S phase in human cells. J Cell Biol 140:1285–1295. doi:10.1083/jcb.140.6.1285

40. Kenny MK, Schlegel U, Furneaux H, Hurwitz J. 1990. The role of human single-stranded DNA binding protein and its individual subunits in simian virus 40 DNA replication. J Biol Chem 265:7693–7700. doi:10.1016/S0021-9258(19)39170-7

41. Kilic S, Lezaja A, Gatti M, Bianco E, Michelena J, Imhof R, Altmeyer M. 2019. Phase separation of 53BP1 determines liquid-like behavior of DNA repair compartments. EMBO J 38:e101379. doi:10.15252/embj.2018101379

42. Ku H, Park G, Goo J, Lee J, Park TL, Shim H, Kim JH, Cho W-K, Jeong C. 2022. Effects of Transcription-Dependent Physical Perturbations on the Chromosome Dynamics in Living Cells. Front Cell Dev Biol 10:822026. doi:10.3389/fcell.2022.822026

43. Laghmach R, Di Pierro M, Potoyan D. 2021. A liquid state perspective on dynamics of chromatin compartments. Front Mol Biosci 8:781981. doi:10.3389/fmolb.2021.781981

44. Leonhardt H, Rahn HP, Weinzierl P, Sporbert A, Cremer T, Zink D, Cardoso MC. 2000. Dynamics of DNA replication factories in living cells. J Cell Biol 149:271–280. doi:10.1083/jcb.149.2.271

45. Levi V, Ruan Q, Plutz M, Belmont AS, Gratton E. 2005. Chromatin dynamics in interphase cells revealed by tracking in a two-photon excitation microscope. Biophys J 89:4275–4285. doi:10.1529/biophysj.105.066670

46. Li CM, Miao Y, Lingeman RG, Hickey RJ, Malkas LH. 2016. Partial purification of a megadalton DNA replication complex by free flow electrophoresis. PLoS ONE 11:e0169259. doi:10.1371/journal.pone.0169259

47. Löb D, Lengert N, Chagin VO, Reinhart M, Casas-Delucchi CS, Cardoso MC, Drossel B. 2016. 3D replicon distributions arise from stochastic initiation and domino-like DNA replication progression. Nat Commun 7:11207. doi:10.1038/ncomms11207

48. Mamberti S, Cardoso MC. 2020. Are the processes of DNA replication and DNA repair reading a common structural chromatin unit? Nucleus 11:66–82. doi:10.1080/19491034.2020.1744415

49. Manders EM, Kimura H, Cook PR. 1999. Direct imaging of DNA in living cells reveals the dynamics of chromosome formation. J Cell Biol 144:813–821. doi:10.1083/jcb.144.5.813

50. Ma H, Tu L-C, Chung Y-C, Naseri A, Grunwald D, Zhang S, Pederson T. 2019. Cell cycle- and genomic distance-dependent dynamics of a discrete chromosomal region. J Cell Biol 218:1467–1477. doi:10.1083/jcb.201807162

51. Mach P, Kos PI, Zhan Y, Cramard J, Gaudin S, Tünnermann J, Marchi E, Eglinger J, Zuin J, Kryzhanovska M, Smallwood S, Gelman L, Roth G, Nora EP, Tiana G, Giorgetti L. 2022. Cohesin and CTCF control the dynamics of chromosome folding. Nat Genet 54:1907–1918. doi:10.1038/s41588-022-01232-7

52. Marshall WF, Straight A, Marko JF, Swedlow J, Dernburg A, Belmont A, Murray AW, Agard DA, Sedat JW. 1997. Interphase chromosomes undergo constrained diffusional motion in living cells. Curr Biol 7:930–939. doi:10.1016/s0960-9822(06)00412-x

53. Michael WM, Ott R, Fanning E, Newport J. 2000. Activation of the DNA replication checkpoint through RNA synthesis by primase. Science 289:2133–2137. doi:10.1126/science.289.5487.2133

54. Moldovan G-L, Pfander B, Jentsch S. 2007. PCNA, the maestro of the replication fork. Cell 129:665– 679. doi:10.1016/j.cell.2007.05.003

55. Montagnoli A, Valsasina B, Brotherton D, Troiani S, Rainoldi S, Tenca P, Molinari A, Santocanale C. 2006. Identification of Mcm2 phosphorylation sites by S-phase-regulating kinases. J Biol Chem 281:10281–10290. doi:10.1074/jbc.M512921200

56. Nagai S, Heun P, Gasser SM. 2010. Roles for nuclear organization in the maintenance of genome stability. Epigenomics 2:289–305. doi:10.2217/epi.09.49

57. Nozaki T, Imai R, Tanbo M, Nagashima R, Tamura S, Tani T, Joti Y, Tomita M, Hibino K, Kanemaki MT, Wendt KS, Okada Y, Nagai T, Maeshima K. 2017. Dynamic Organization of Chromatin Domains Revealed by Super-Resolution Live-Cell Imaging. Mol Cell 67:282–293.e7. doi:10.1016/j.molcel.2017.06.018

58. Oliveira GM, Oravecz A, Kobi D, Maroquenne M, Bystricky K, Sexton T, Molina N. 2021. Precise measurements of chromatin diffusion dynamics by modeling using Gaussian processes. Nat Commun 12:6184. doi:10.1038/s41467-021-26466-7

59. Pliss A, Malyavantham K, Bhattacharya S, Zeitz M, Berezney R. 2009. Chromatin dynamics is correlated with replication timing. Chromosoma 118:459–470. doi:10.1007/s00412-009-0208-6

60. Prelich G, Kostura M, Marshak DR, Mathews MB, Stillman B. 1987. The cell-cycle regulated proliferating cell nuclear antigen is required for SV40 DNA replication in vitro. Nature 326:471–475. doi:10.1038/326471a0

61. Rausch C, Zhang P, Casas-Delucchi CS, Daiß JL, Engel C, Coster G, Hastert FD, Weber P, Cardoso MC. 2021. Cytosine base modifications regulate DNA duplex stability and metabolism. Nucleic Acids Res 49:12870–12894. doi:10.1093/nar/gkab509

62. Ritter C, Wollmann T, Lee JY, Imle A, Müller B, Fackler OT, Bartenschlager R, Rohr K. 2021. Data fusion and smoothing for probabilistic tracking of viral structures in fluorescence microscopy images. Med Image Anal 73:102168. doi:10.1016/j.media.2021.102168

63. Sadoni N, Cardoso MC, Stelzer EHK, Leonhardt H, Zink D. 2004. Stable chromosomal units determine the spatial and temporal organization of DNA replication. J Cell Sci 117:5353–5365. doi:10.1242/jcs.01412

64. Sage D, Neumann FR, Hediger F, Gasser SM, Unser M. 2005. Automatic tracking of individual fluorescence particles: application to the study of chromosome dynamics. IEEE Trans Image Process 14:1372–1383. doi:10.1109/TIP.2005.852787

65. Saxton MJ. 1997. Single-particle tracking: the distribution of diffusion coefficients. Biophys J 72:1744– 1753. doi:10.1016/S0006-3495(97)78820-9

66. Schermelleh L, Solovei I, Zink D, Cremer T. 2001. Two-color fluorescence labeling of early and mid-to-late replicating chromatin in living cells. Chromosome Res 9:77–80. doi:10.1023/a:1026799818566

67. Scipioni L, Lanzanó L, Diaspro A, Gratton E. 2018. Comprehensive correlation analysis for super-resolution dynamic fingerprinting of cellular compartments using the Zeiss Airyscan detector. Nat Commun 9:5120. doi:10.1038/s41467-018-07513-2

68. Shaban HA, Barth R, Bystricky K. 2018. Formation of correlated chromatin domains at nanoscale dynamic resolution during transcription. Nucleic Acids Res 46:e77. doi:10.1093/nar/gky269

69. Shin Y, Brangwynne CP. 2017. Liquid phase condensation in cell physiology and disease. Science 357. doi:10.1126/science.aaf4382

70. Shukron O, Seeber A, Amitai A, Holcman D. 2019. Advances Using Single-Particle Trajectories to Reconstruct Chromatin Organization and Dynamics. Trends Genet 35:685–705. doi:10.1016/j.tig.2019.06.007

71. Simson R, Yang B, Moore SE, Doherty P, Walsh FS, Jacobson KA. 1998. Structural mosaicism on the submicron scale in the plasma membrane. Biophys J 74:297–308. doi:10.1016/S0006-3495(98)77787-2

72. Smith PR, Morrison IE, Wilson KM, Fernández N, Cherry RJ. 1999. Anomalous diffusion of major histocompatibility complex class I molecules on HeLa cells determined by single particle tracking. Biophys J 76:3331–3344. doi:10.1016/S0006-3495(99)77486-2

73. Sparvoli E, Levi M, Rossi E. 1994. Replicon clusters may form structurally stable complexes of chromatin and chromosomes. J Cell Sci 107 (Pt 11):3097–3103. doi:10.1242/jcs.107.11.3097

74. Spegg V, Altmeyer M. 2021. Biomolecular condensates at sites of DNA damage: More than just a phase. DNA Repair (Amst) 106:103179. doi:10.1016/j.dnarep.2021.103179

75. Sporbert A, Gahl A, Ankerhold R, Leonhardt H, Cardoso MC. 2002. DNA polymerase clamp shows little turnover at established replication sites but sequential de novo assembly at adjacent origin clusters. Mol Cell 10:1355–1365. doi:10.1016/s1097-2765(02)00729-3

76. Tanaka S, Hu SZ, Wang TS, Korn D. 1982. Preparation and preliminary characterization of monoclonal antibodies against human DNA polymerase alpha. J Biol Chem 257:8386–8390.

77. Tunnacliffe E, Chubb JR. 2020. What is a transcriptional burst? Trends Genet 36:288–297. doi:10.1016/j.tig.2020.01.003

78. Uchino S, Ito Y, Sato Y, Handa T, Ohkawa Y, Tokunaga M, Kimura H. 2022. Live imaging of transcription sites using an elongating RNA polymerase II-specific probe. J Cell Biol 221. doi:10.1083/jcb.202104134

79. Van C, Yan S, Michael WM, Waga S, Cimprich KA. 2010. Continued primer synthesis at stalled replication forks contributes to checkpoint activation. J Cell Biol 189:233–246. doi:10.1083/jcb.200909105

80. Vesela E, Chroma K, Turi Z, Mistrik M. 2017. Common Chemical Inductors of Replication Stress: Focus on Cell-Based Studies. Biomolecules 7. doi:10.3390/biom7010019

81. Vincent JA, Kwong TJ, Tsukiyama T. 2008. ATP-dependent chromatin remodeling shapes the DNA replication landscape. Nat Struct Mol Biol 15:477–484. doi:10.1038/nsmb.1419

82. Walter J, Schermelleh L, Cremer M, Tashiro S, Cremer T. 2003. Chromosome order in HeLa cells changes during mitosis and early G1, but is stably maintained during subsequent interphase stages. J Cell Biol 160:685–697. doi:10.1083/jcb.200211103

83. Waseem NH, Lane DP. 1990. Monoclonal antibody analysis of the proliferating cell nuclear antigen (PCNA). Structural conservation and the detection of a nucleolar form. J Cell Sci 96 (Pt 1):121–129. doi:10.1242/jcs.96.1.121

84. Yan S, Michael WM. 2009a. TopBP1 and DNA polymerase alpha-mediated recruitment of the 9-1-1 complex to stalled replication forks: implications for a replication restart-based mechanism for ATR checkpoint activation. Cell Cycle 8:2877–2884. doi:10.4161/cc.8.18.9485

85. Yan S, Michael WM. 2009b. TopBP1 and DNA polymerase-alpha directly recruit the 9-1-1 complex to stalled DNA replication forks. J Cell Biol 184:793–804. doi:10.1083/jcb.200810185

86. Yao NY, O’Donnell M. 2010. SnapShot: The replisome. Cell 141:1088, 1088.e1. doi:10.1016/j.cell.2010.05.042

87. Yao NY, O’Donnell ME. 2016. Evolution of replication machines. Crit Rev Biochem Mol Biol 51:135–149. doi:10.3109/10409238.2015.1125845

88. Zidovska A, Weitz DA, Mitchison TJ. 2013. Micron-scale coherence in interphase chromatin dynamics. Proc Natl Acad Sci USA 110:15555–15560. doi:10.1073/pnas.1220313110

89. Nichols WW, Murphy DG, Cristofalo VJ, Toji LH, Greene AE, Dwight SA. 1977. Characterization of a new human diploid cell strain, IMR-90. Science 196:60–63. doi:10.1126/science.841339

90. Celikay K, Chagin VO, Cardoso MC, and Rohr K. (2022) DenoiseReg: Unsupervised Joint Denoising and Registration of Time-Lapse Live Cell Microscopy Images Using Deep Learning, Proc. IEEE International Symposium on Biomedical Imaging (ISBI 2022), Kolkata, India, 28-31 March, 2022, IEEE Piscataway, NJ

